# Kinetics of elastic recoil in the wings of the cicada, *Dundubia rufivena*

**DOI:** 10.1101/2025.07.30.667726

**Authors:** Parvez Alam, Puspa Restu Sayekti, Fahrunnida, Colin Robert, Bambang Retnoaji, Catharina Maria Alam

## Abstract

In this paper, we quantitatively characterise the kinetics of elastic recoil in the wings attached to the cicada *Dundubia rufivena*. We use high speed videography to deduce the velocities of recoil motion in both bending (B) and torsion (T), in downstroke and upstroke modes of flapping. Elastic recoil is faster in downstroke (DS) than it is in upstroke (US), reaching average velocities, 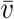, 244 °.*s*^−1^ in DS-B vs 6434 °.*s*^−1^ in US-B, and 5887 °.*s*^−1^ in DS-T vs 4545 °.*s*^−1^ in US-T. Our results also therefore evidence that bending velocities during elastic recoil, are higher to the velocities in torsion, and conclude that this is a result of the necessary geometrical distances that need to be covered over a stroke. The wings do not act alone as biological springs, but rather, we find that the stiffnesses of the wings (1.7-3.6 GPa) are higher when attached to the cicada body, than they are when detached. This evidences thoracical involvement as part of the biological spring enabling elastic recoil, indicating that elastic recoil of flapping wings should be approached from a systems perspective, rather than solely through a localised understanding of wing mechanics.

## 1. Introduction

Energy is required for flight [1], and while insect muscles are efficient, they are not a sole source of input energy [2]. The energy requirements of flight in *Drosophila hydei* for example, are minimised through thoracical elastic elements, which recover excess energy during deceleration in the first half stroke of a flapping cycle [3]. Similar mechanisms of energy saving in flight are evident in wider bodied insects like cicadas [4], which are the world’s loudest insects [5], producing sound through striated viscoelastic tymbals [6] that vibrate at extremely high (*>* 3 kHz) frequencies [7]. Cicadas carry relatively heavy bodies using wings that exhibit simpler designs [8] than those of other higher order insects [9]. The estimation of a cicada’s aerodynamic, inertial and mechanical power in a free forward flight indicates that the total mechanical power in a downstroke is negative. As such by extension, this power could potentially be stored and restored in a subsequent stroke [4]. In nature, elastic springs contribute to insect flight efficiency. Weis-Fogh [10] for example, evidenced the presence of passive elastic forces as drivers for acceleration in the wings of dead locust, by lifting wings into stroke positions, and releasing them. They noted that kinematic responses were present, even after all flight muscles were removed from the animal, indicating that residual elastic energy was provided by the extant materials of the wing, some of which is derived from resilin protein in the wing base cuticle [11]. Resilin is a rubber-like material with high energy absorption capabilities [12,13] and respectable tolerances to fracture damage [14]. Passive wing deformation plays a role in aerodynamic force production and is a characteristic influenced by its flexural stiffness. The leading edge of the wing is considered a major contributor to stiffness, as it is the part of the wing that maintains rigidity during flight. Both the tip and trailing edge are contrarily thinner and more flexible regions of a wing [15,16], thus contributing less to wing stiffness and more to wing flexibility.

Although the flexibility of cicada wings are known to be influential to both lift [17] and translational damping [18], the deformation of cicada wings and any implications these may have on energy saving have not been researched. These are therefore areas that are only vaguely understood by means of inference, through aligned works that have been conducted on other insect species. In this paper, we consider the kinetics of passive responses in pre-pitched, and released, cicada wings. This phenomenon is known as elastic recoil (or, pitch recoil) [19–22], which is in essence, a ‘biological spring’, similar to many other springs that are found in a wide variety of insects including cockroaches, springtails, planthoppers, locusts and more. The effects of such springs are typically researched from a mechanical perspective, with sparse attention given to the kinetics of motion. Yet, the kinetics of active insect flapping has been well researched and as such, an additional understanding of the kinetics of elastic recoil will provide insights on important issues such as wing efficiency. Osbourne [23] for example, concluded that insects in general, fly in a way that minimises the mechanical power required by the animal. This may involve specialised wing-wing interaction techniques such as ‘clap and fling’ exhibited by *Drosophila melanogaster* [24], the judicious control of flap-trajectories [25–27], or through the use of elastic elements incorporated into the architectural design of the wing itself, which can give rise to power losses through span-wise flexure [28] and damping related losses [29]. The 3-dimensional structure of the wing controls in essence, the way in which a wing deforms during flapping and through pitch recoil. This affects its aerodynamic functionality, with morphological features such as venation, functional gradation and flexion lines being of particular importance in deformation control [30].

To enable our study on wing kinetics and stiffness, we employ high speed videography, capturing the elastic recoil of wings in two translational phases (downstroke and upstroke). We use the video output to calculate the angular velocities of the recoiling wings to deduce and compare the kinetics in bending and torsion for both upstroke and downstroke, and furthermore, to contrasts these against stored elastic energy, which we measure directly using dynamic mechanical tests.

## 2. Results

### 2.1 Morphological identification of cicada species

Through morphological identification, we found that all cicadas captured are of the species, *Dundubia rufivena* (Walker, 1850). [31] The following key characteristics were identified, which are typical to this species, as described by Overmeer and Duffels [32]. Male wing colouration was hyaline. The body lengths were between 27.3 mm and 30.6 mm. There was an absence of black fascia on the head. The operculum of *Dundubia rufivena* was notably ‘spoon-shaped’ not laterally curved. The operculum length was almost twice as long as it was wide, and its apical segment was approximately two-fold broader than the base. Likewise, female wing colouration was hyaline. There was no black fascia on its head and its body length was between 30. mm and 30.7 mm, with a bright green and ochraceous color. There was an absence of black lines on the mesonotum. Its abdomen had an obtuse tip, which was curvaceous.

### 2.2. Morphometrics

The mean widths (± standard deviations) of the left wings, 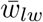, and right, 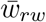, were measured at 11.9 mm (±1.5 mm) and 10.8 mm (±1.3 mm), respectively, with there being a 9.2% difference between the mean widths on the left and right wings overall. The coefficients of variation, *CoV*, for 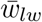 and 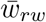 are thus 12.8% and 12.1%, respectively. Mean lengths from base to tip (±standard deviations) of the left wings, 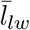 and right, 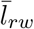, were measured at 36.7 mm (±2.6 mm) and 34.8 mm (± 4.8 mm), respectively, with there being a 5.2% difference between the mean lengths on the left and right wings overall. The *CoV* for 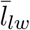 and 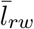 are thus 7.1% and 13.7%, respectively. ^;^Based on these values, the mean aspect ratios of the wings, 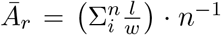, are 3.1 (± 0.2) and 3.2 (± 0.2) for the left and right wings, respectively, where *n* is the number of insect samples used in the sample set (*n* = 4), *l* is the length of a wing and *w* is its width. The *CoV* for the left and right wing aspect ratios are 6.5% and 6.4%, respectively. The average thickness of the leading edges of the wings are 260 *µ*m (±10 *µ*m), while the average body lengths and body widths of the cicadas were measured at 29.3 mm (± 1.8 mm) and 11.0 mm (± 1.3 mm), respectively.

### 2.3. Kinetics of elastic recoil

The velocities of wing bending (B) and torsion (T) during elastic recoil were measured using a Chronos 1.4 (Krontech, Canada) high-speed camera at 2357 frames per second (resolution 1024×576) and a playback speed of one frame per second to enable a frame by frame estimation of time. Left and right wings were filmed individually for each animal, from the anterior to posterior view to map bending kinetics, and from the lateral view to map torsion kinetics. The elastic recoil was measured for both downstroke (DS) and upstroke (US) in line with measured angles reported by Park et al [33] such that the pre-release starting positions for elastic recoil measurements were: Right Wing DS = 340°, Left Wing DS = 20°, Right Wing US = 220°, and Left Wing US = 140°. Figure 1(a) provides a polar plot showing the kinetics for all wings measured in upstroke and downstroke bending (US-B and DS-B, respectively) and torsion (US-T and DS-T, respectively), with colour codes and schematic illustrations provided in Figure 1(b). The results are represented by truncated violin plots in Figure 1(c) for all wings in each of the sample sets of DS-B, DS-T, US-B and US-T. There is a larger range of velocities mapped from elastic recoil in DS-B (20000 °.*s*^−1^) than DS-T (12727 °.*s*^−1^), as well in US-B (8485 °.*s*^−1^) than US-T (1818 °.*s*^−1^), and we note that the highest velocities reached when comparing between these two sets are in bending (in both US and DS). Applying a two-way ANOVA analysis and Tukey’s posthoc test, we report a significant difference (p≤0.001) between DS-B and DS-T, as well as for between US-B and US-T (p≤ 0.05). DS-B also shows a significant difference between DS-B and US-B (p ≤0.0001), while no significant difference is noted to occur between DS-T and US-T sample sets. When comparing the mean velocities, 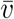, for each group, we note that DS-B 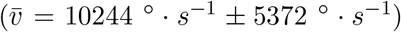 while DS-T 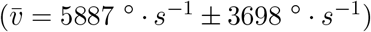 and US-B 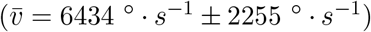 and US-T 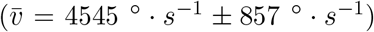, where each 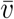 is ± the standard deviation. We note that the 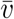 of both DS-B and DS-T are more than one standard deviation apart, as are those for US-B and US-T.

**Figure 1.**
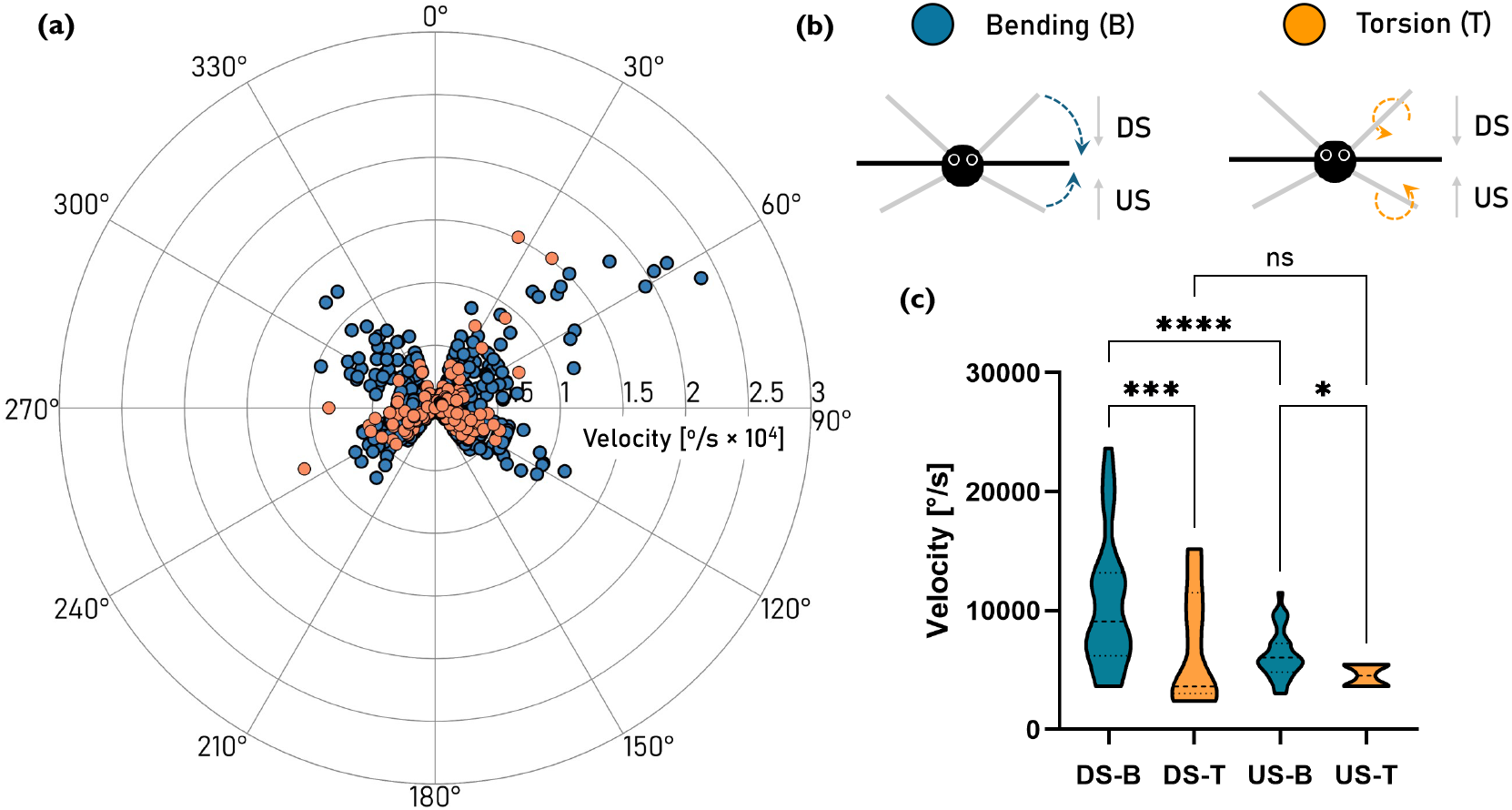
(a) Cicada wing kinetics presented in a polar plot mapping all measured bending and torsional velocities in both downstroke and upstroke (b) colour codes and schematic illustrations (c) Truncated violin plots for all wings collated showing downstroke bending (DS-B), downstroke torsion (DS-T), upstroke bending (US-B) and upstroke torsion (US-T). Significance bars from a two-way ANOVA analysis and Tukey’s posthoc test are provided where *n* = 4, *r* = 32 and *n* = 4, *r* = 32 for downstroke and upstroke bending tests, respectively, while *n* = 4, *r* = 7 and *n* = 4, *r* = 8 for torsion tests in downstroke and upstroke, respectively. Here, *n* is the number of insects used, *r* is the number of replicates, ns indicates there is no significant difference, * indicates that p ≤ 0.05, *** indicates that p ≤ 0.001, and **** indicates that p ≤ 0.0001.

In Figure 2, we provide kinetics data in bending for both the left and right wings in downstroke and upstroke. These are presented in a polar plot in Figure 2(a) and as box and whisker plots in Figure 2(b). Since the data was not normally distributed, we used the Kruskal-Wallis test as well as Dunn’s multiple comparisons test. The mean velocity was compared for each sample set (*n* = 4, *r* = 40 left wing, *n* = 4, *r* = 24 for right wing) where *n* is the number of insects used and *r* is the number of replicates. We note that while there are no significant differences between the downstroke of the left vs right wing (DS Left vs DS Right), or for the upstroke of the left vs right wing (US Left vs US Right), there are significant differences between the downstroke and upstroke of the left wing (p ≤ 0.05) and of the right wing (p ≤ 0.05).

**Figure 2.**
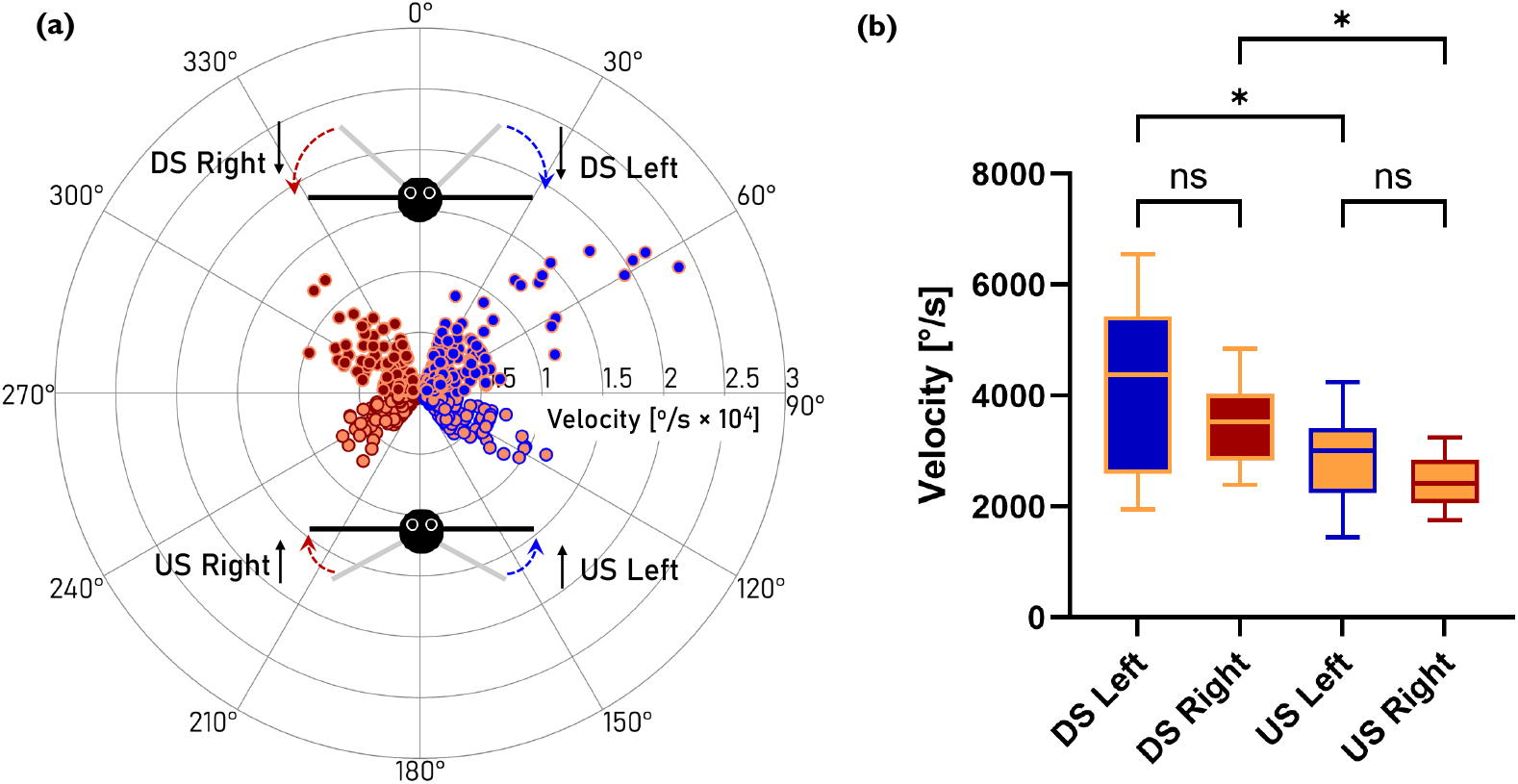
(a) Cicada wing bending kinetics presented in a polar plot showing values of both downstroke (DS) and upstroke (US) for both left and right wings (b) a box and whisker plot comparing the velocities using Kruskal-Wallis test alongside Dunn’s multiple comparisons test. Here, ns indicates there is no significant difference, * indicates that p ≤ 0.05. Here, *n* = 4, *r* = 40 for left wing measurements, and *n* = 4, *r* = 24 for right wing measurements.

Figure 3(a) shows a polar plot of wing kinetics in torsion for both the left and right wings in downstroke and upstroke. These are plotted as box and whisker diagrams in Figure 3(b) for each sample set (DS and US for both left and right wings). Significance bars are not shown in this figure as there were no significant differences founds between the four groups following a non parametric Kruskal-Wallis test alongside Dunn’s multiple comparisons test. The mean velocity was compared for each sample set (*n* = 4, *r* = 8 left wing, *n* = 4, *r* = 7 for right wing).

**Figure 3.**
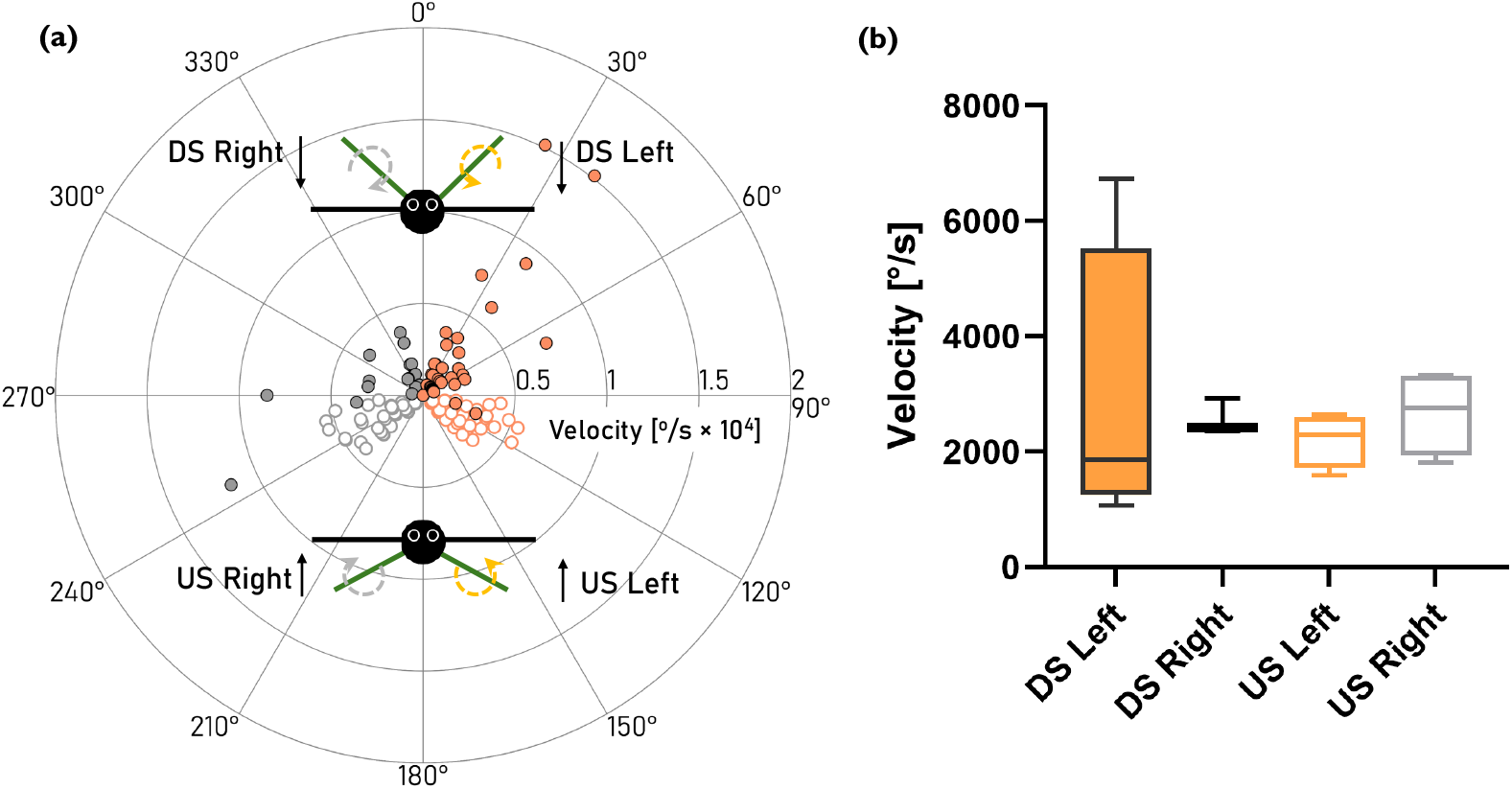
(a) Cicada wing torsion kinetics presented in a polar plot showing values of both downstroke (DS) and upstroke (US) for both left and right wings (b) a box and whisker plot comparing the velocities using Kruskal-Wallis test alongside Dunn’s multiple comparisons test. No significant differences were noted between any of the sample sets. Here, *n* = 4, *r* = 8 for left wing measurements, *n* = 4, *r* = 7 for right wing measurements.

### 2.4. Mechanical properties

The flexural moduli, *E*_*f*_, of cicada wings were measured at ramp rates of 1 Hz and 10 Hz, and at ramp amplitudes of 0.1 mm and 0.3 mm. These are shown as DS vs US in Figure 4(a) for the left wing and in (b) for the right wing. Using a two-way ANOVA alongside Sidak’s posthoc test, we note that there are no significant differences between DS and US for either the left or right wings tested (*n* = 3 for each sample set and tested parameter). The loss factor in these structures was extremely low, thus they store more energy than they dissipate. We therefore treat the storage modulus in flexure as equivalent to a flexural modulus and use the term flexural modulus hereinafter.

**Figure 4.**
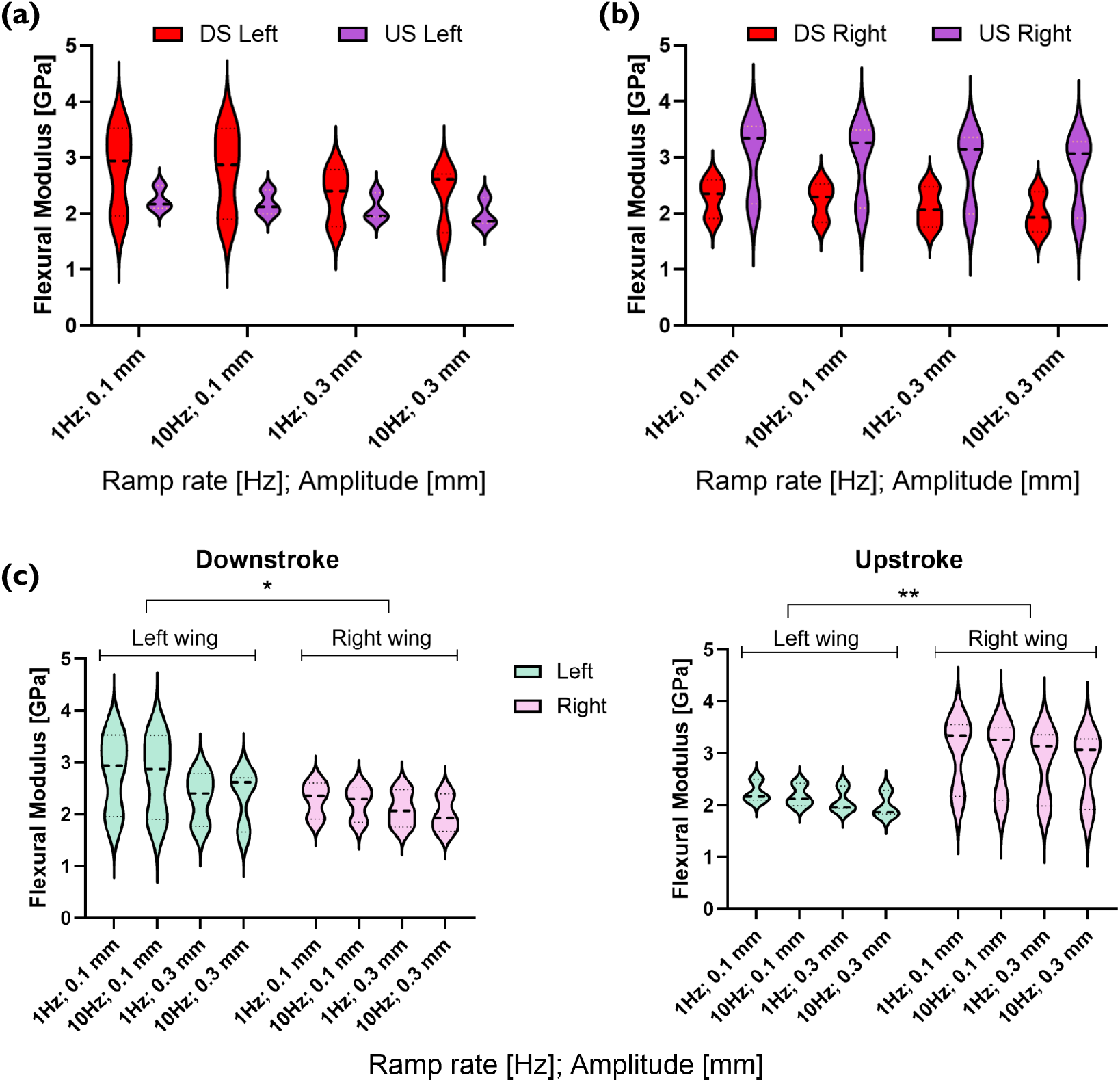
(a) Flexural moduli in downstroke and upstroke compared for left wings at different ramp rates and amplitudes (b) flexural moduli in downstroke and upstroke compared for right wings at different ramp rates and amplitudes and (c) flexural moduli compared of all left wings against all right wings tested in downstroke and in upstroke. Statistical analyses were performed using two-way ANOVA. * = *p* ≤ 0.05 ** = *p* ≤ 0.01. Here, *n* = 3, *r* = 50 wing pairs.

The mean flexural moduli, 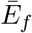, of DS Left and US Left are 2.55 GPa and 2.13 GPa, respectively, indicating a difference of 16.5% between the means of these groups. 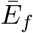 of DS Right and US Right are 2.15 GPa and 2.89 GPa, respectively, indicating a difference of 26.5% between the means of these groups. In Figure 4(c) we present the same data but grouped for the left and right wings in DS and US. Here, we use two-way ANOVA (*n* = 3, *r* = 50 wing pairs). The flexural moduli values were significantly higher (*p*≤ 0.05) in the left wing than in the right wing during downstroke. Contrarily, measuring the same wings in upstroke, we note that the flexural moduli are significantly higher (*p*≤ 0.01) in the right wings.

The values in Figure 4 were based on tests conducted on only the wing. As such, we compare these against stiffness tests conducted on the wings as attached to the body of the cicada, Figure 5 where (a) shows the data for downstroke and (b) shows the data for upstroke. In this figure we refer to the modulus, rather than the flexural modulus. This is because when the wing is attached to the body of the cicada through its joint, a bending load will cause both bending and torsional loading at the joint due to the geometrical construction of the joint. The modulus therefore, while being loaded in flexure, should be considered more generically as a stiffness, or modulus, as used herein. The data is normally distributed, thus we use a two-way ANOVA (*n* = 3, *r* = 50 wing pairs) to analyse the level of difference in DS or in US, between attached wings or wings only. The two-way ANOVA p-value for DS is *p*≤ 0.001 and for US *p* ≤0.05, indicating that the body of the cicada plays a role in the elastic recoil, by increasing the level of elastic energy stored prior to recoil release. In the downstroke, on average there is a 9% improvement in the modulus for attached wings as compared to detached wings, while in upstroke there is an 14% enhancement in the modulus for attached wings as compared to detached wings.

**Figure 5.**
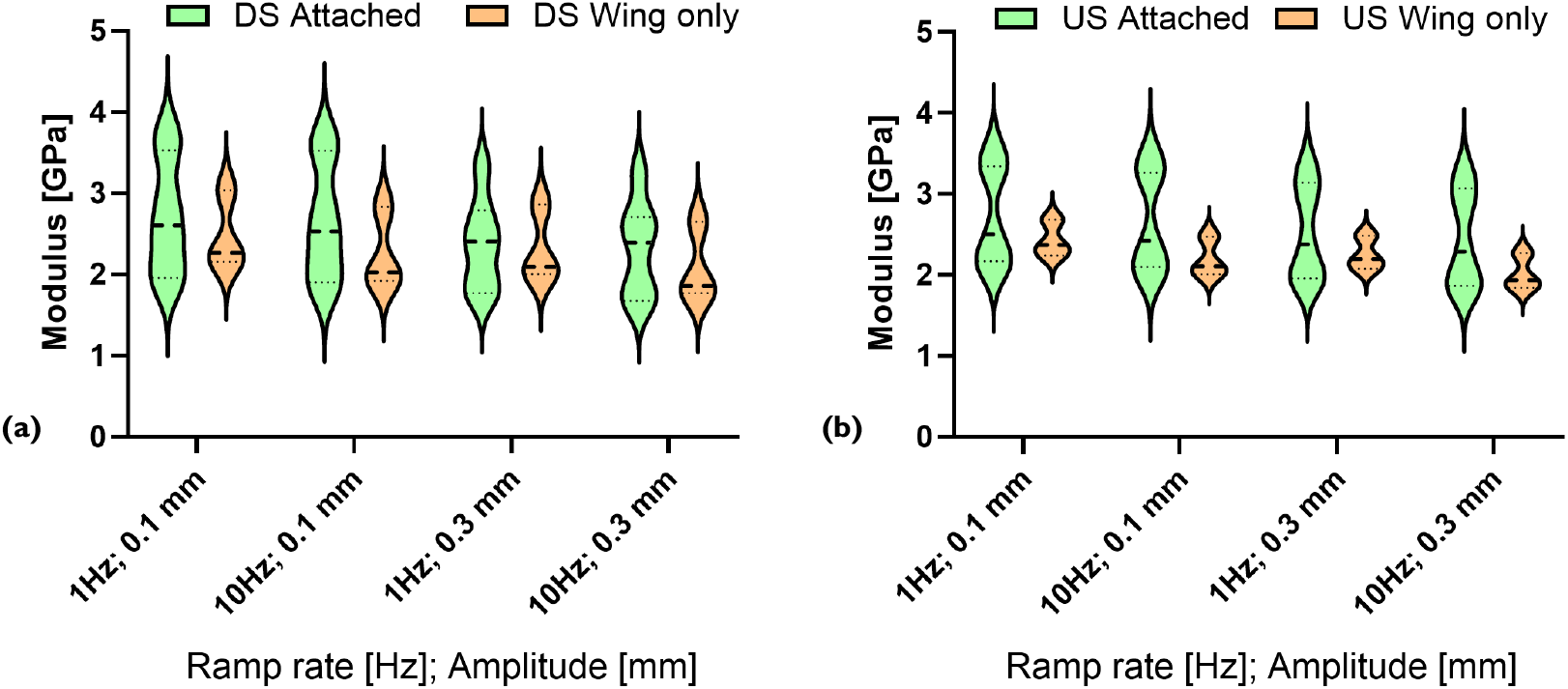
Flexural moduli of both the wing as attached to the cicada body (Attached) and separated from the body (Wing only) at different ramp rates and amplitudes (a) downstroke and (b) upstroke. Statistical analyses for normally distributed sample sets using two-way ANOVA, in DS *p* ≤ 0.001 and for US *p* ≤ 0.05. Here, *n* = 3, *r* = 50 wing pairs.

## 3. Discussion

It has long been understood that elastic energy storage leads to recoil in insect wings, and that this mechanism enhances the efficiency, *ϕ*, of the wing such that in wings where there is perfect elastic energy storage, 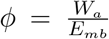, where *W*_*a*_ is aerodynamic work and *E*_*mb*_ is metabolic energy consumption [34]. While torsional flexibility is a pronounced factor [35], essential in enabling rotational elastic recoil [36], other factors such as wing camber and its effect on the biomechanics of lateral wing flexion are also significant contributors [37]. While torsio-flexional wing kinematics have been reported for a range of insects in view of aerodynamic flight [38–41], there are fewer studies that have mapped the kinetics of insect wing dynamics [42,43]. Here, we find that the angular velocities of cicada wings are generally larger as a sample set in downstroke than in upstroke, in both bending and torsion. These are recorded herein as ranging between *∼*3000 and 24000 °.*s*^−1^, which translates roughly to between *∼*52 and 419 rad.*s*^−1^. These are in line with angular velocities reported in the literature for other flapping insects such as hawkmoths (*∼* 20 - 155 rad.*s*^−1^) [44,45], and dragonflies (from *∼*200 to 600 rad.*s*^−1^ (max)) [46]. Of the collated measures for kinetics, we find that downstroke bending had on average, the fastest elastic recoil (*∼*10000 °.*s*^−1^), which was slightly higher than that in upstroke bending (*∼*6500 °.*s*^−1^), both of which exhibited faster elastic recoil velocities than in torsion (*∼*6000 °.*s*^−1^ in DS and *∼*4500 ° *s*^−1^ in US). There is a geometrical logic to this, since wings are longer than they are wide, the distance that needs to be covered during either a downstroke or an upstroke is greater in bending, which works along the length of the wing, than it is in torsion, which works in relation to the width of the wing [39]. While bending is important in lift generation, torsion is used to control airflow, while maximising aerodynamic efficiency [39,47–49] and since as is evident from the polar plot in Figure 1(a), both torsion and bending reach a similar physical end point at null velocity after recoil, indicating that the kinetics of elastic recoil is finely tuned to enable the spatial posturing of the wing for the next phase of kinematic motion.

The wings of the cicada, *Dundubia rufivena*, while slightly different in size between left to right sides 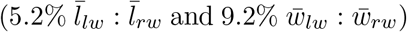 do still reveal similar geometrical profiles, as evidenced by the similar aspect ratios on left and right sides of each cicada (3.1:3.2 for left:right). Aspect ratios in insect wings are indicative of differences in flight style [50], wing loading [51], and ecological as well as environmental pressures [52]. The aspect ratios of *Dundubia rufivena* are notably similar to those of fruit flies, honeybees and hawkmoths [53], which possess a range of values between 2.9 and 3.3. Here, we note that while there are notable differences in the mechanical properties between the left and right sides of the wings in bending (cf. Figure 4(c)), that this does not affect elastic recoil in terms of their kinetics (cf. Figures 2(b) and 3(b)). We suggest therefore, that the while there may be physical differences between wings the similar aspect ratios of the wings enable a degree of consistency in the kinetics of elastic recoil in both downstroke and upstroke.

In this paper, we provide evidence to show that cicada wings act as biological springs [34,54], with an overall modulus value range (1.7-3.6 GPa) in line with other known biological springs such as cockroach legs which are recorded at 1.7-3.1 GPa [54]. The elastic recoil in cicada wings, similar to other insects, will reduce the energetic requirements of flight [43], and changes to the stiffness of the material will have a proportionate effect on elastic recoil [55]. We also show that the ‘spring’ is in fact not solely based on the wing but that the body and joint of the cicada play a vital role in increasing the stiffness of the spring in both upstroke and downstroke (cf. Figure 5). Though spring-like quality of the wings is related to factors such as venation and size [15], the importance of the body in spring-like action has, as far as we know, not been quantified in cicadas or any other insect prior to this study. As such, it would seem that while there is benefit in considering isolated structures and materials in insects to describe biological spring-like characteristics in animals [15,54], that in fact the spring-like characteristics may in many cases be more relevantly described from a systems perspective. This would include several connected body parts for elastic recoil, minimally the wing, the thorax and the joint/interface, which collectively contribute to spring-like characteristics. The application of fixed boundaries at the joint in modelled wings [15,56,57] probably therefore insufficiently describes the deformation-resistance characteristics as compared to more aptly modelled multi-body systems where the wing has been attached to a compliant thorax [58,59].

## 4. Conclusions

Elastic recoil is the release of stored energy during insect flight. This mechanism is important as it reduces muscular power generation burdens, enabling longer durations of flight through energy conservation. Here, we quantitatively characterise the elastic recoil kinetics of the cicada *Dundubia rufivena* using high speed videography, and further characterise the stiffness properties of the wings, both as separated from the cicada body, and as attached to the cicada body. We find that the elastic recoil kinetics in a downstroke, is superior to that of an upstroke, and conclude that this may be because downstrokes control lift and propulsion more than do upstrokes, and there is a greater requirement for power generation in a downstroke. We further find that the kinetics of bending have an overall larger range than the kinetics of torsion (wing twisting) and suggest that this may simply be due to the geometrical requirements of the wing (with shorter travel distance requiring lower velocities to complete a single stroke. We finally find that stiffness properties of the wings align with those of other biological springs reported in the literature, while duly noting that the stiffness properties of wings still attached to the cicada thorax are higher in stiffness than those that have been detached from the body. This indicates there is a thoracical contribution to elastic recoil, which has been characterised both kinematically and mechanically in this work.

## 5. Methods

### 5.1. Sample collection

Cicadas were collected in Tamantirto, Kasihan, Bantul Regency -7.8256389 110.3263697 (05°16’12.2” S latitude; 110°32’63697” E longitude) at night using a light trap until a required number of samples were captured (10 individuals). The cicadas were divided into two groups for preservation using different methods, either chemical preservation or drying. Chemical preservation was conducted using 70% ethanol, while the drying process was completed in an oven at 40°C.

### 5.2. Morphometrics

Morphometric analyses were conducted using using images of four of the cicadas uploaded into ImageJ software (*n* = 4). We measured forewing length (base to tip), forewing width, body length and body width. Wing thickness was measured using an electronic micrometer with a rated accuracy of 0.001 mm.

### 5.3. Morphological identification of cicada species

Morphological identification was conducted based on identification keys provided by Overmeer and Duffels [32]. The characteristics analysed in males included wing colour, body length, and features of the operculum and head. In females, we analysed the wing colour, body length, and the morphological characteristics of the operculum, head, mesonotum and abdomen tip.

### 5.4. Elastic recoil experiments

The kinetics of elastic recoil from pre-strained and released cicada wings were deduced by high speed videography. Specifically, the velocities of wing bending and torsion during elastic recoil were measured using a Chronos 1.4 (Krontech, Canada) high-speed camera at 2357 frames per second (resolution 1024 ×576) and a playback speed of one frame per second. This enabled a frame by frame estimation of time. Both left and right wings were filmed individually for each animal, from the anterior to posterior view to measure the bending velocities, and from the lateral view to measure the torsional velocities. Elastic recoil was measured for both downstroke and upstroke in line with measured angles reported by Park et al. [33], such that the pre-release starting positions for elastic recoil measurements were: Right Wing DS = 340°, Left Wing DS = 20°, Right Wing US = 220°, and Left Wing US = 140°. Each cicada was positioned and held using a clamp stand. Prior to testing, the cicadas were conditioned by humidification, and tests were conducted in a 21°C environment with a relative humidity of 55%. A total of four cicadas were used (*n* = 4) and replicate kinetic tests, *r*, on the same cicada wings were conducted.

### 5.5. Mechanical testing

The first set of tests considered the flexural stiffness of only the wings (as removed from the cicada body), which were measured by single cantilever bending in a Tritec 2000 DMA (Dynamic Mechanical Analyser), clamping the root end of the wing and loading the wing tip. The second set of tests considered the flexural stiffness of the wings while still attached to the cicada body. These were measured by single cantilever bending in a Tritec 2000 DMA (Dynamic Mechanical Analyser), clamping the body of the cicada between paper pads to prevent damage, and loading at the wing tip. A temperature scan was conducted in single frequency mode, with start and end temperatures of 24°C and 28°C, respectively. Test speeds were varied independently using 1 Hz and 10 Hz for each wing. Test amplitudes (cantilever displacements) were varied independently using 0.1 mm and 0.3 mm for each wing tested. The cross-sectional areas for each wing were averaged for each wing across the gauge length (30 mm) using thicknesses and widths systematically measured at regular intervals. Each wing yielded 50 stiffness results (*r* = 50), which was averaged for each of three insects (*n* = 3), where *r* is the number of replicates and *n* is the number of insects used. Examples of the tests with the wing attached to the body, and with the wing detached from the body, are provided as Electronic Supplementary Videos.

## Supporting information

Electronic Supplementary Materials

## 6. Author Contributions

Parvez Alam conceived the project, wrote the abstract, introduction, results, discussion and conclusions sections, edited the methods section, prepared the plots and videos for use as supplementary materials, analysed and interpreted data, prepared the polar plots, formatted figures 1-3, and supervised the research team. Puspa Restu Sayekti along with Fahrunnida captured the cicada samples, conducted the kinetics testing, conducted the mechanical testing, performed the morphometric studies, morphologically identified the cicada species, packaged the data for later analysis, and wrote the methods section. Colin Robert designed the DMA method used mechanically characterising the cicadas. Bambang Retoaji prepared administrative documentation and secured permits for materials transfer. Catharina Maria Alam prepared the violin plots and the box-and-whisker plots, and conducted statistical analyses for all data outputs from this project.

## 7. Acknowledgements

We wish to acknowledge Sukirno, Ph.D. for teaching light trap methods to Fahrunnida and Puspa Restu Sayekti.

## 8. Electronic Supplementary Materials

The following supplementary materials are provided:

- Electronic Supplementary Video 1: Example of elastic recoil from a downstroke release
- Electronic Supplementary Video 2: Example of elastic recoil from an upstroke release
- Electronic Supplementary Video 3: Example of a dynamic mechanical test on a wing attached to the body of a cicada (1 Hz upstroke)
- Electronic Supplementary Video 4: Example of a dynamic mechanical test on a wing attached to the body of a cicada (10 Hz upstroke)
- Electronic Supplementary Video 5: Example of a dynamic mechanical test on a wing attached to the body of a cicada (1 Hz downstroke)
- Electronic Supplementary Video 6: Example of a dynamic mechanical test on a wing attached to the body of a cicada (10 Hz downstroke)
- Electronic Supplementary folder containing kinetics polar plots for individual wings in bending separated into L and R with DS or US
- Electronic Supplementary folder containing kinetics polar plots for individual wings in torsion separated into L and R with DS or US

## 9. Data Availability

Data is provided as Electronic Supplementary Materials and will also be available from the corresponding author on reasonable request.

## 11. Ethics Statement

All animal care, handling, and experimental procedure in this study were carried out in out in accordance with guidelines available through The Committee of Ethical Clearance for Pre-clinical Research of The Integrated Laboratory of Research and Testing, Gadjah Mada University, Yogyakarta, Indonesia. Indonesian Law does not require an ethical approval arrangement for the capture and post-euthanasia research of the cicadas used in this study. Gadjah Mada University already has legally authorised permission to conduct field work and sample collections. The samples captured are not on the list of protected animals. According to Law of the Republic of Indonesia No. 18 Year 2009 Article 66 about Animal Welfare (Undang Undang Republik Indonesia No. 18 Tahun 2009 Pasal 66 Tentang Kesejahteraan Hewan).

