## Supplementary figures and images for "Kinetics of elastic recoil in the wings of the cicada, *Dundubia rufivena*"

### L_DS_a.tiff

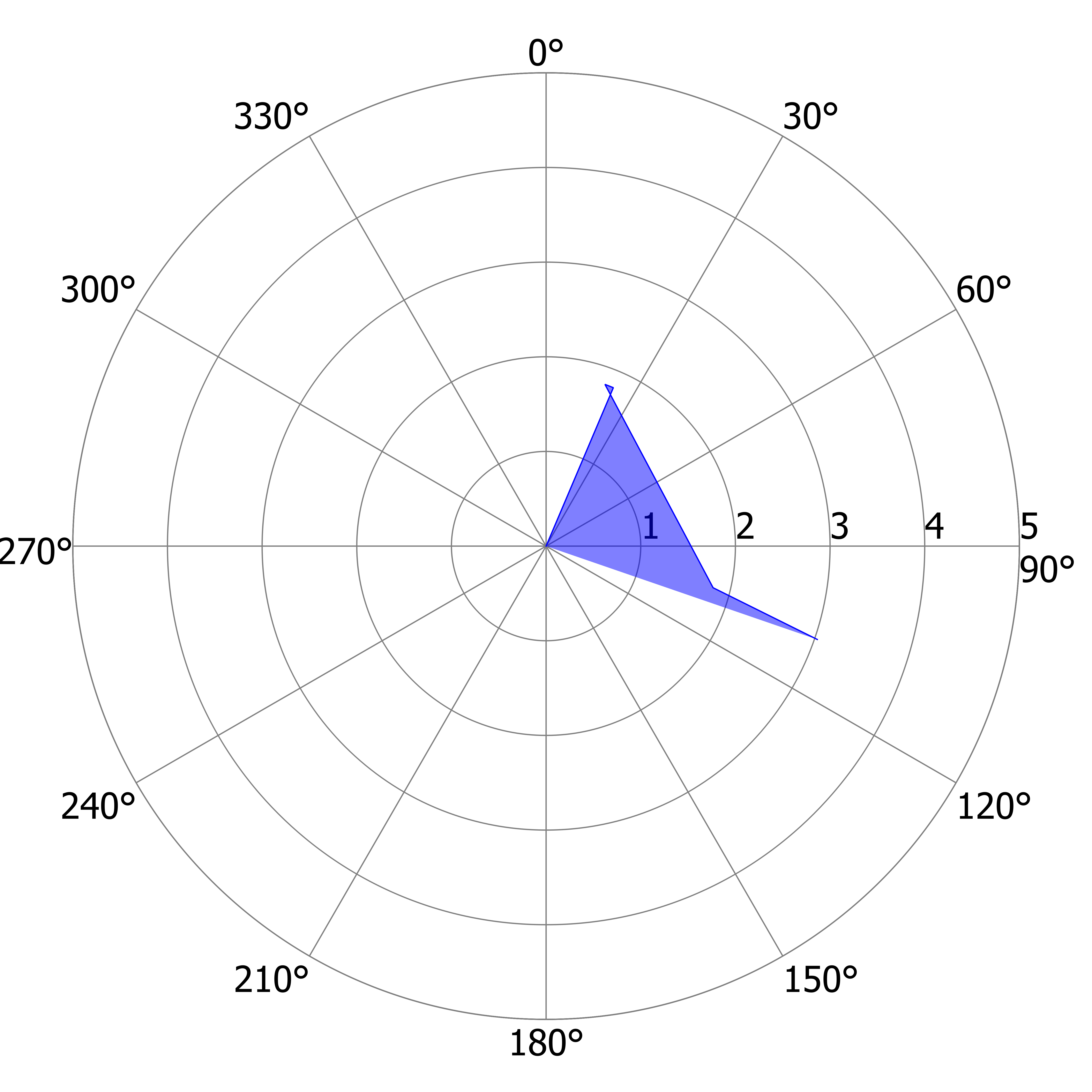

### L_DS_a.tiff

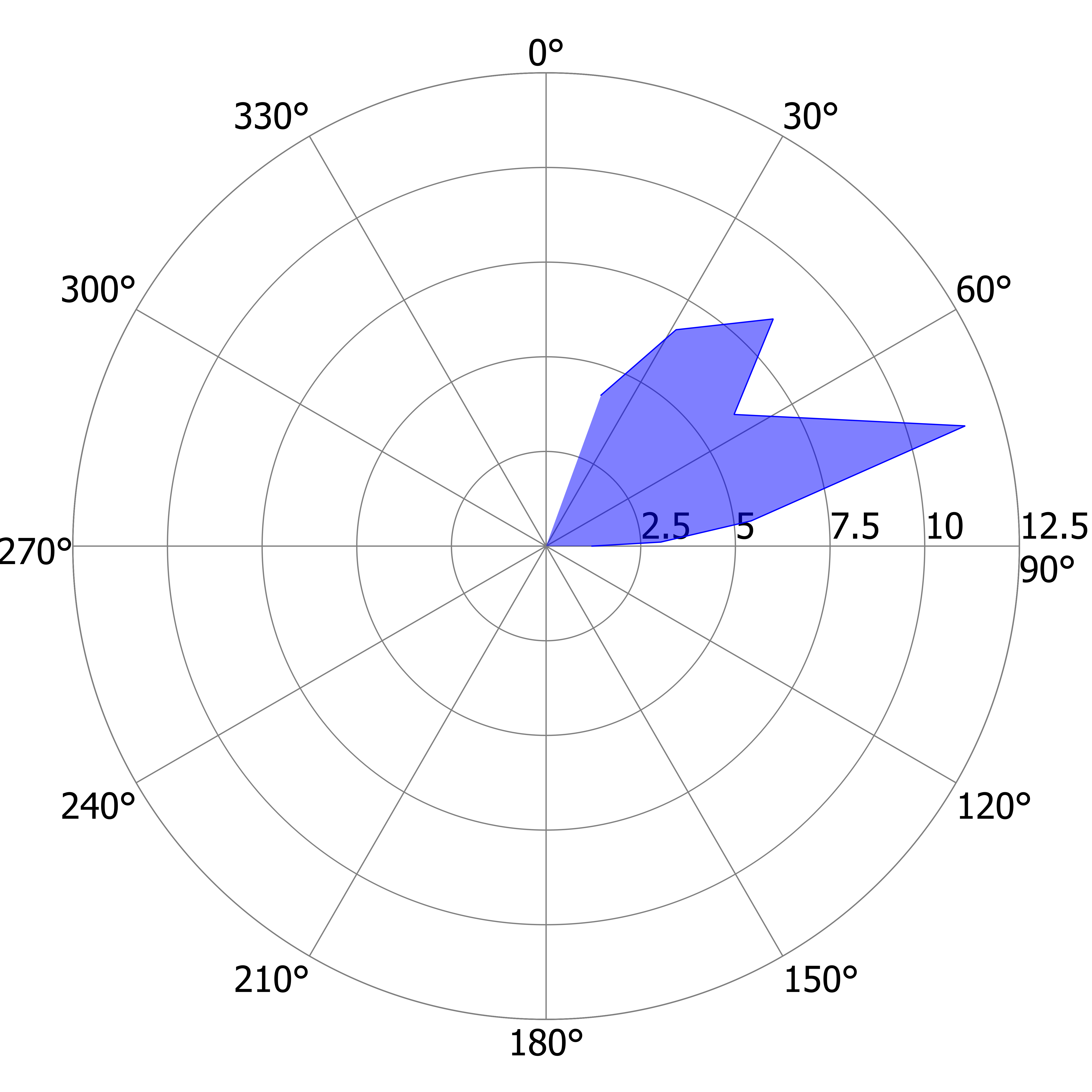

### L_DS_b.tiff

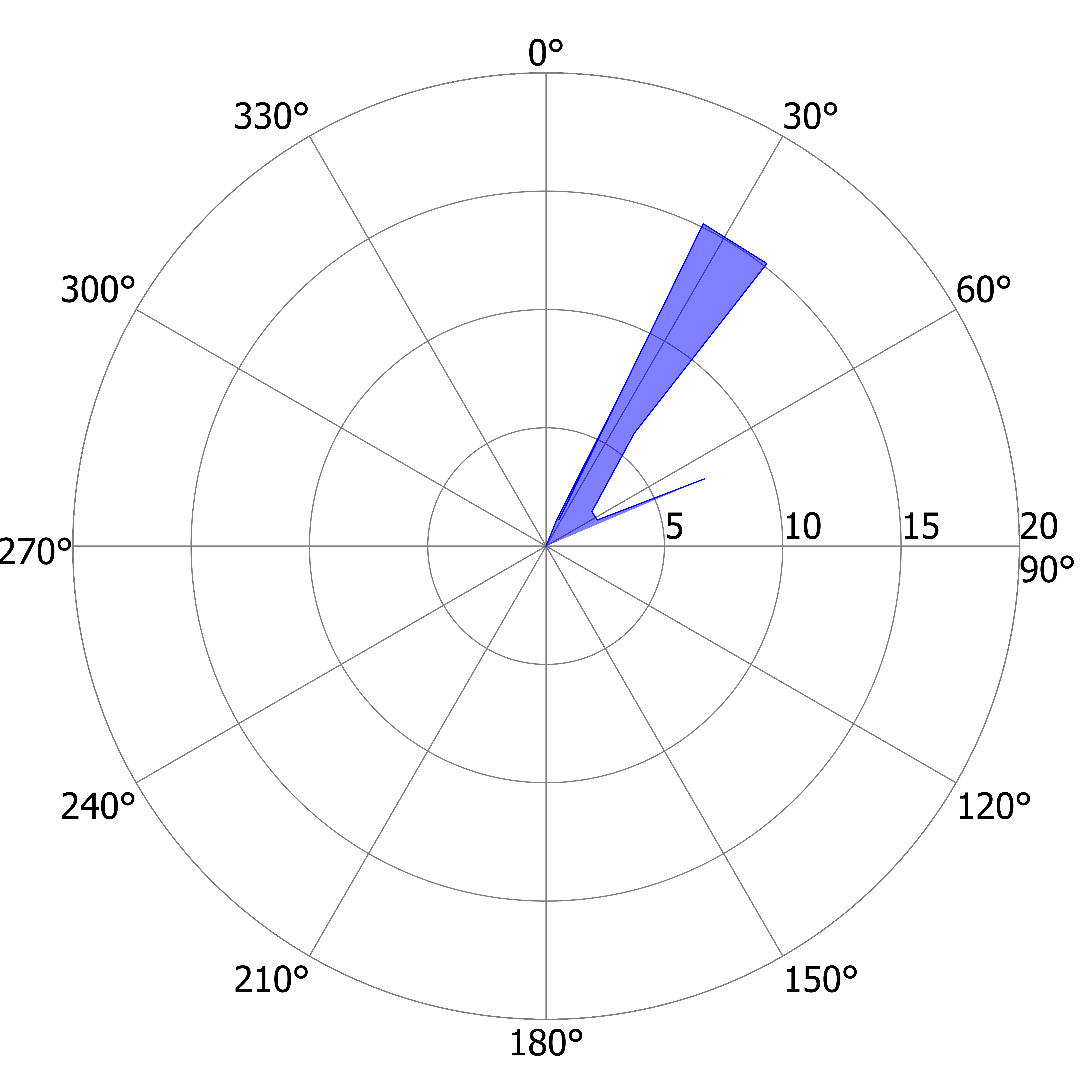

### L_DS_b.tiff

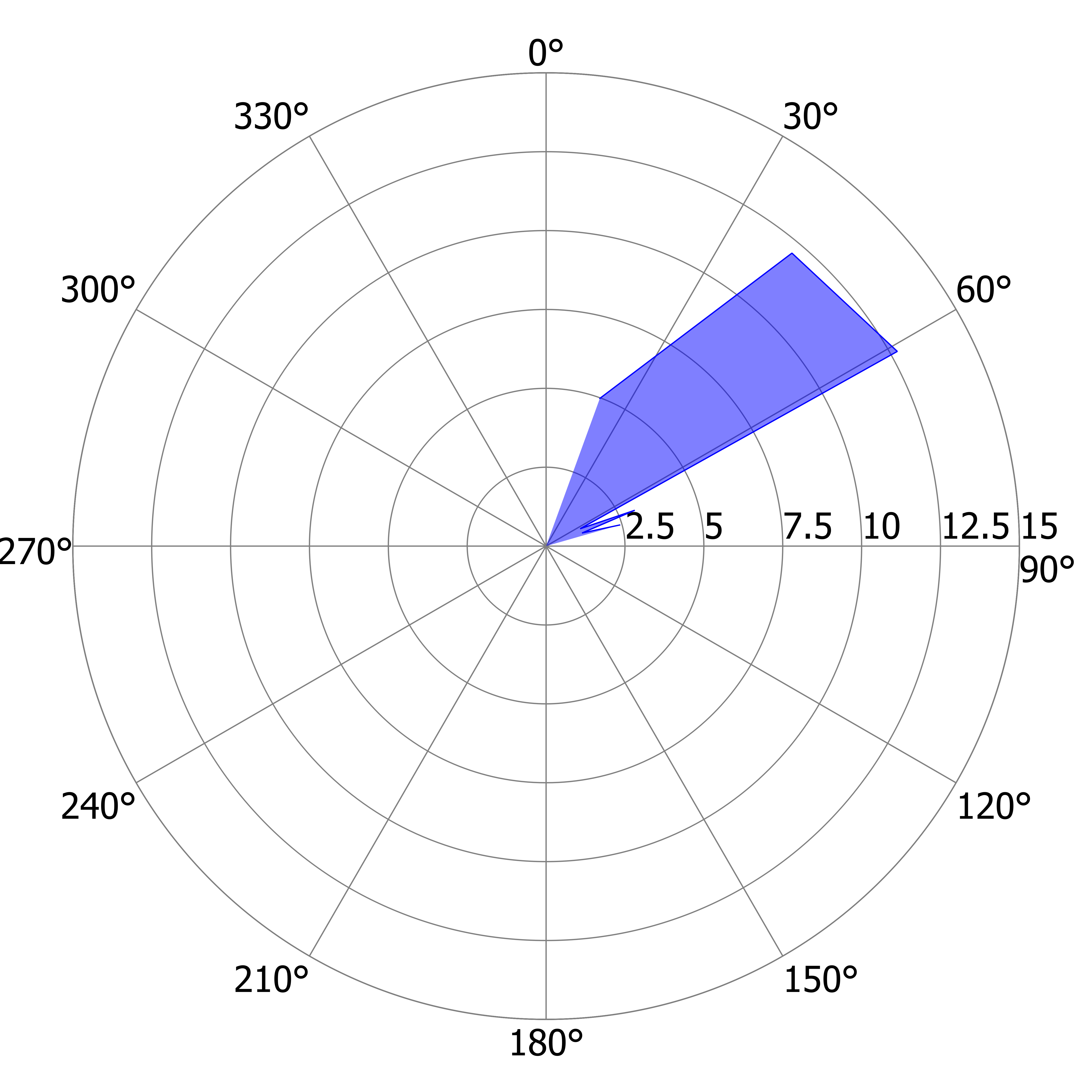

### L_DS_c.tiff

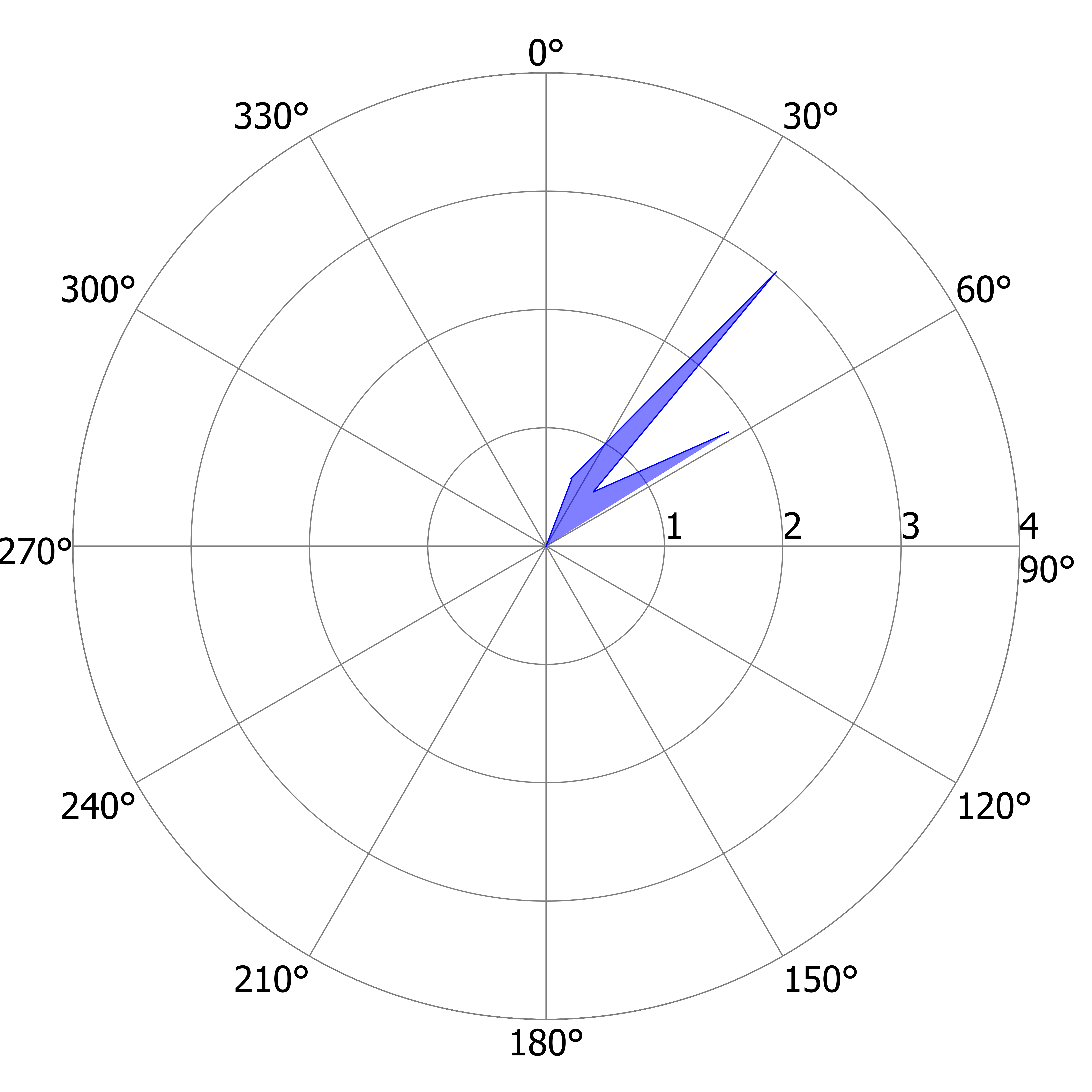

### L_DS_c.tiff

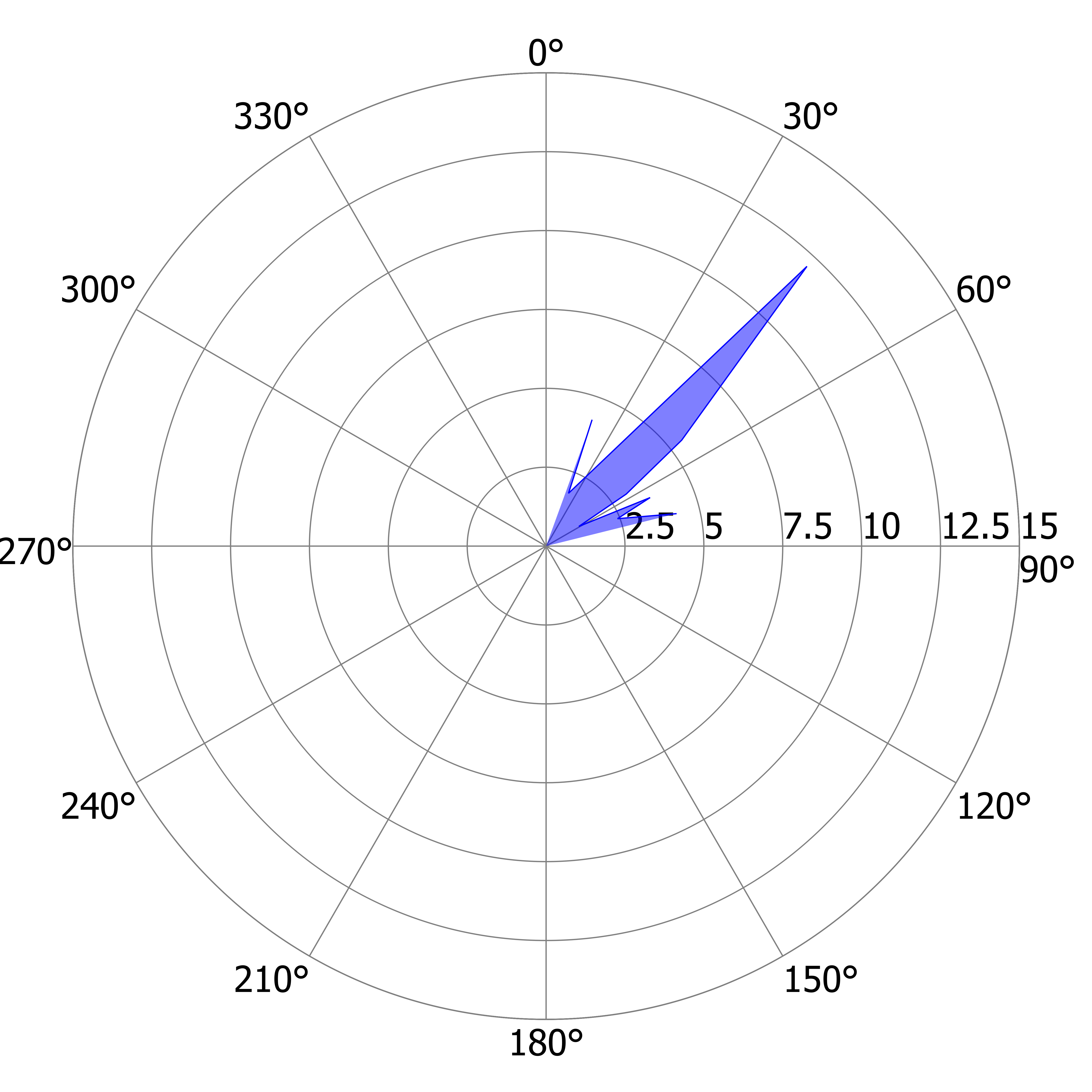

### L_DS_d.tiff

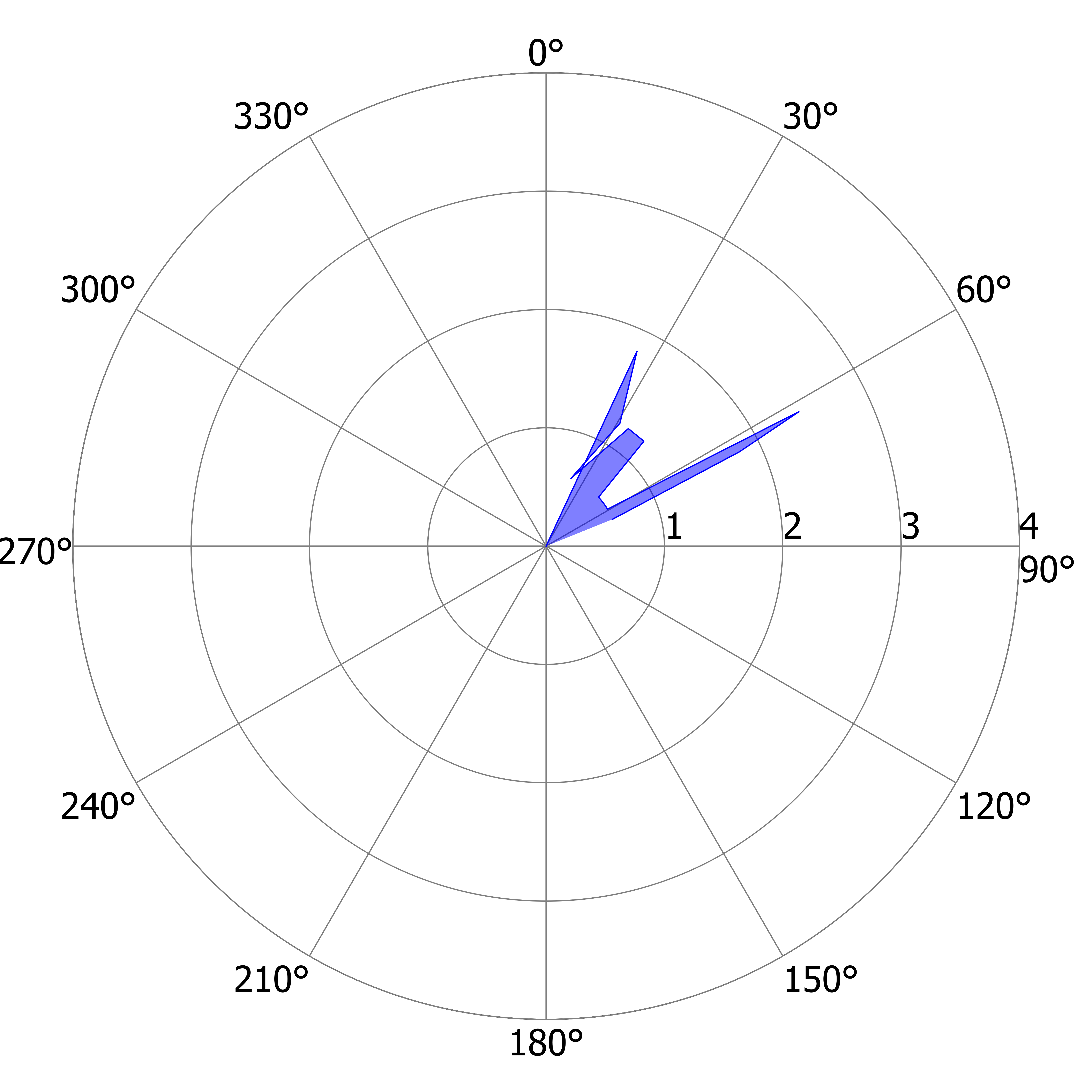

### L_DS_d.tiff

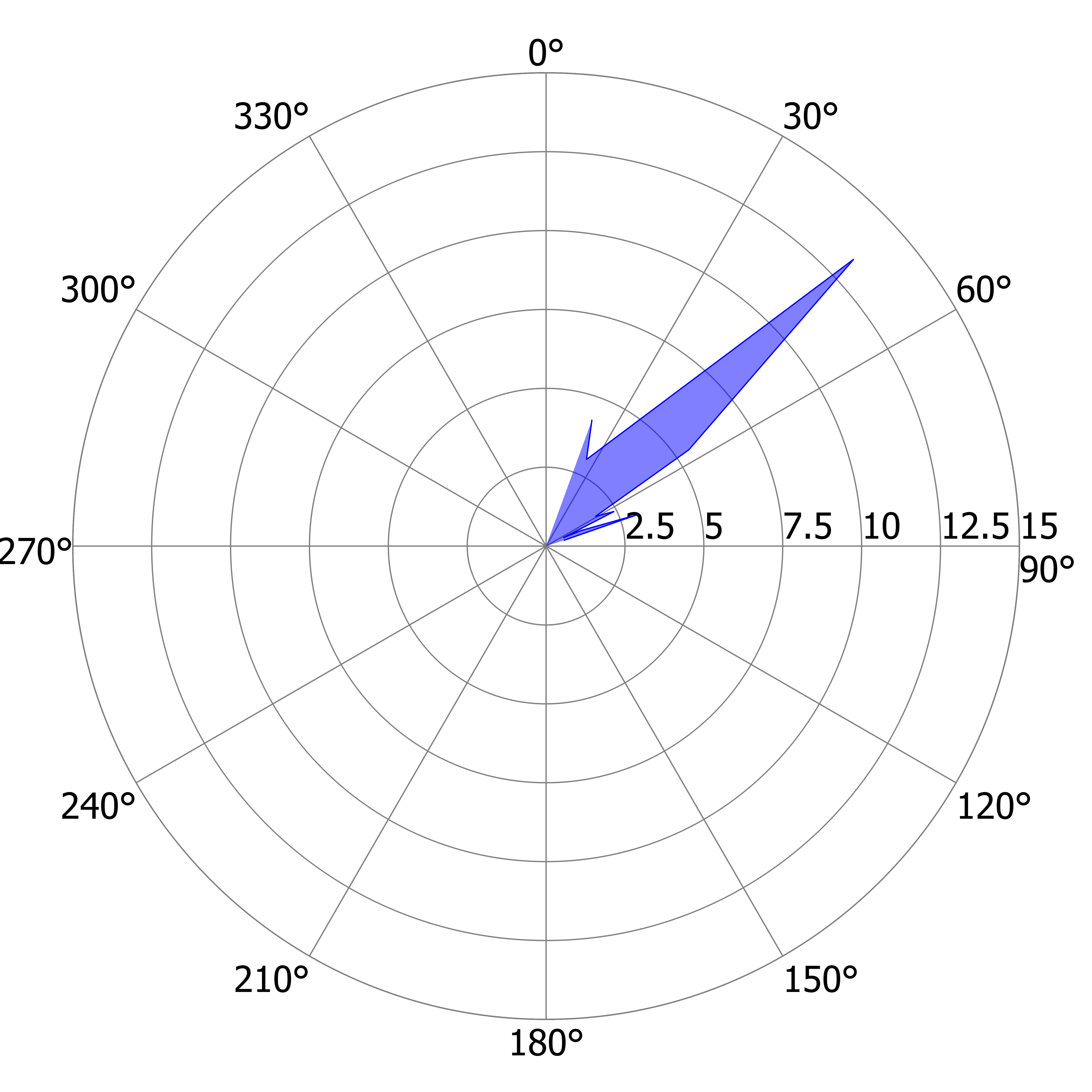

### L_DS_e.tiff

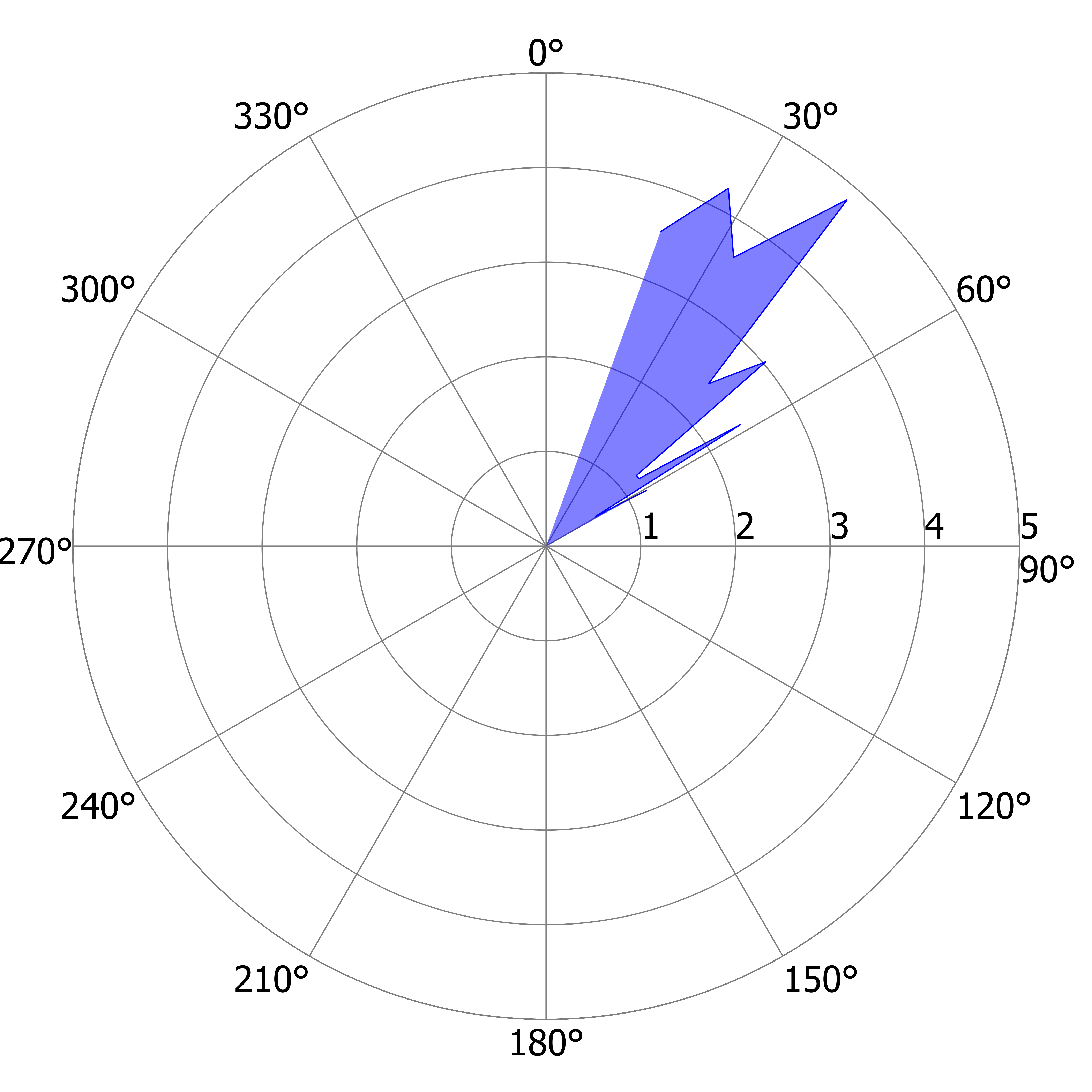

### L_DS_f.tiff

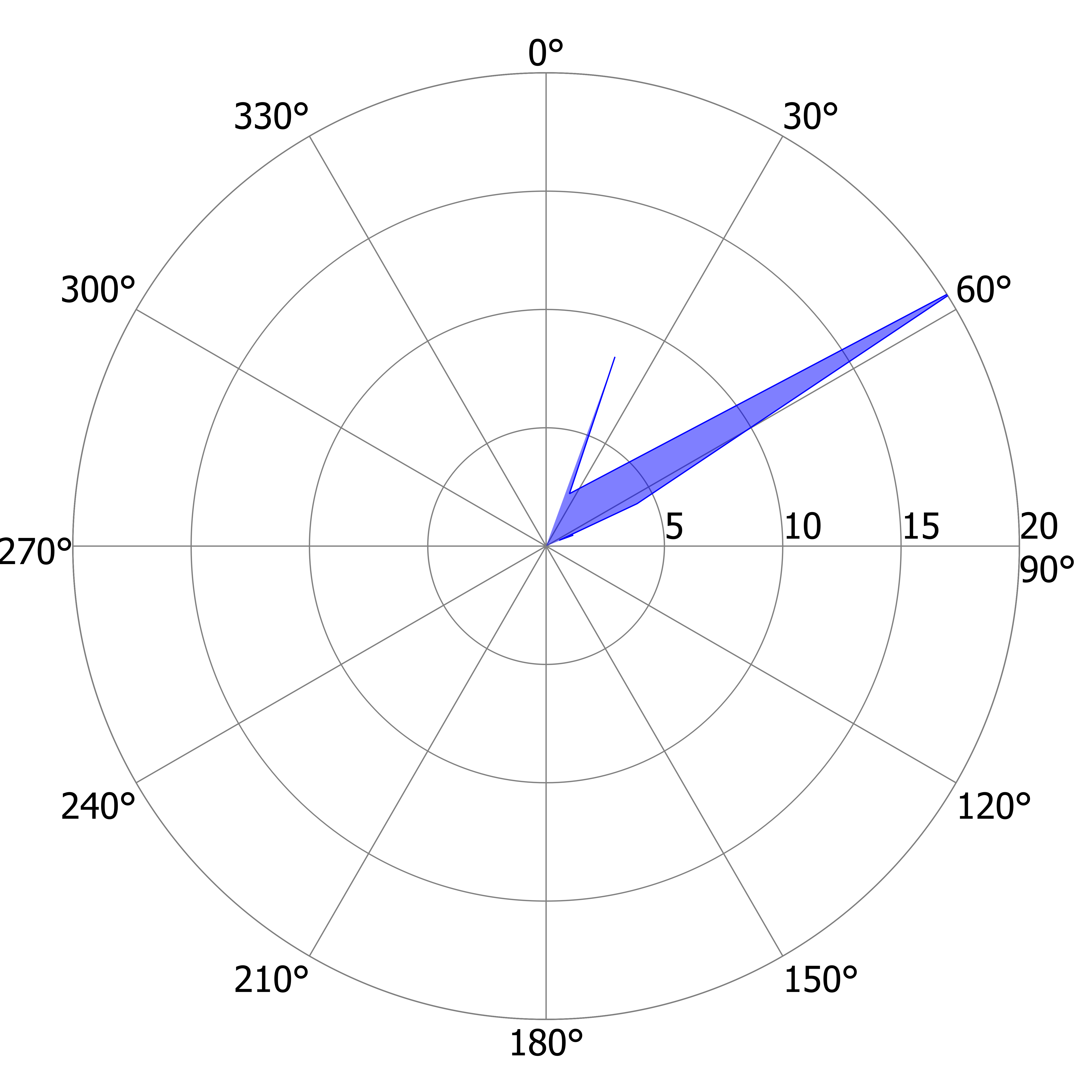

### L_DS_g.tiff

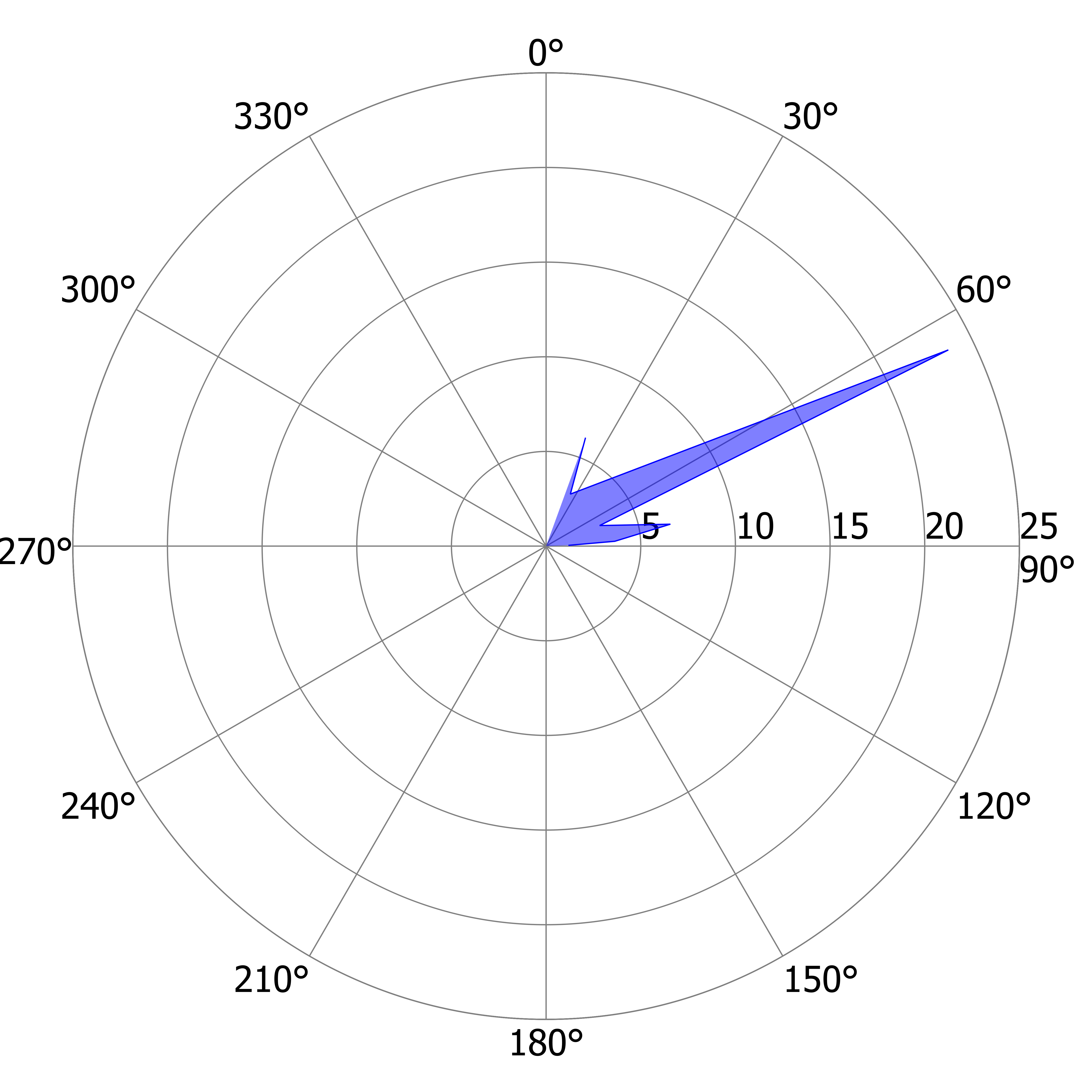

### L_DS_h.tiff

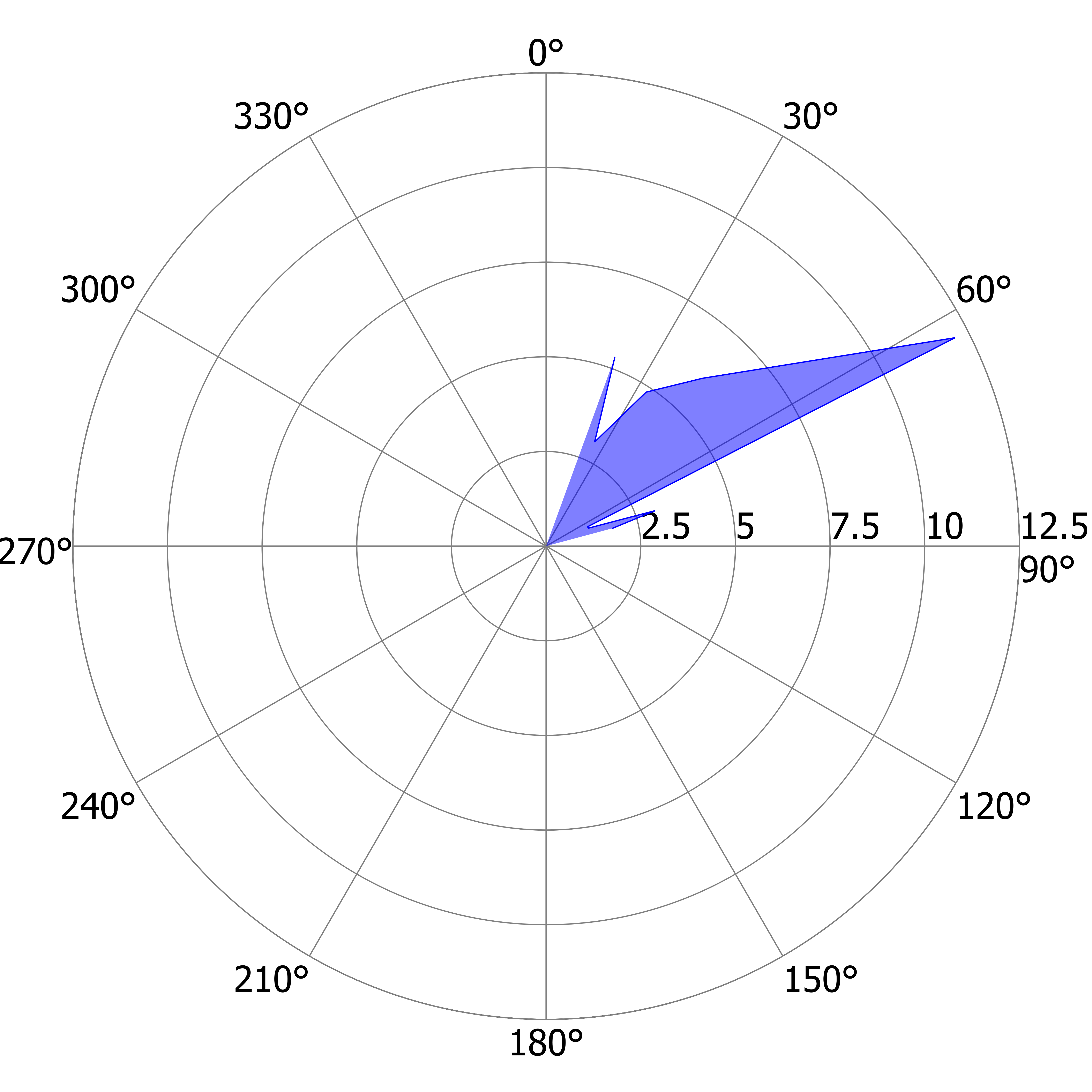

### L_DS_i.tiff

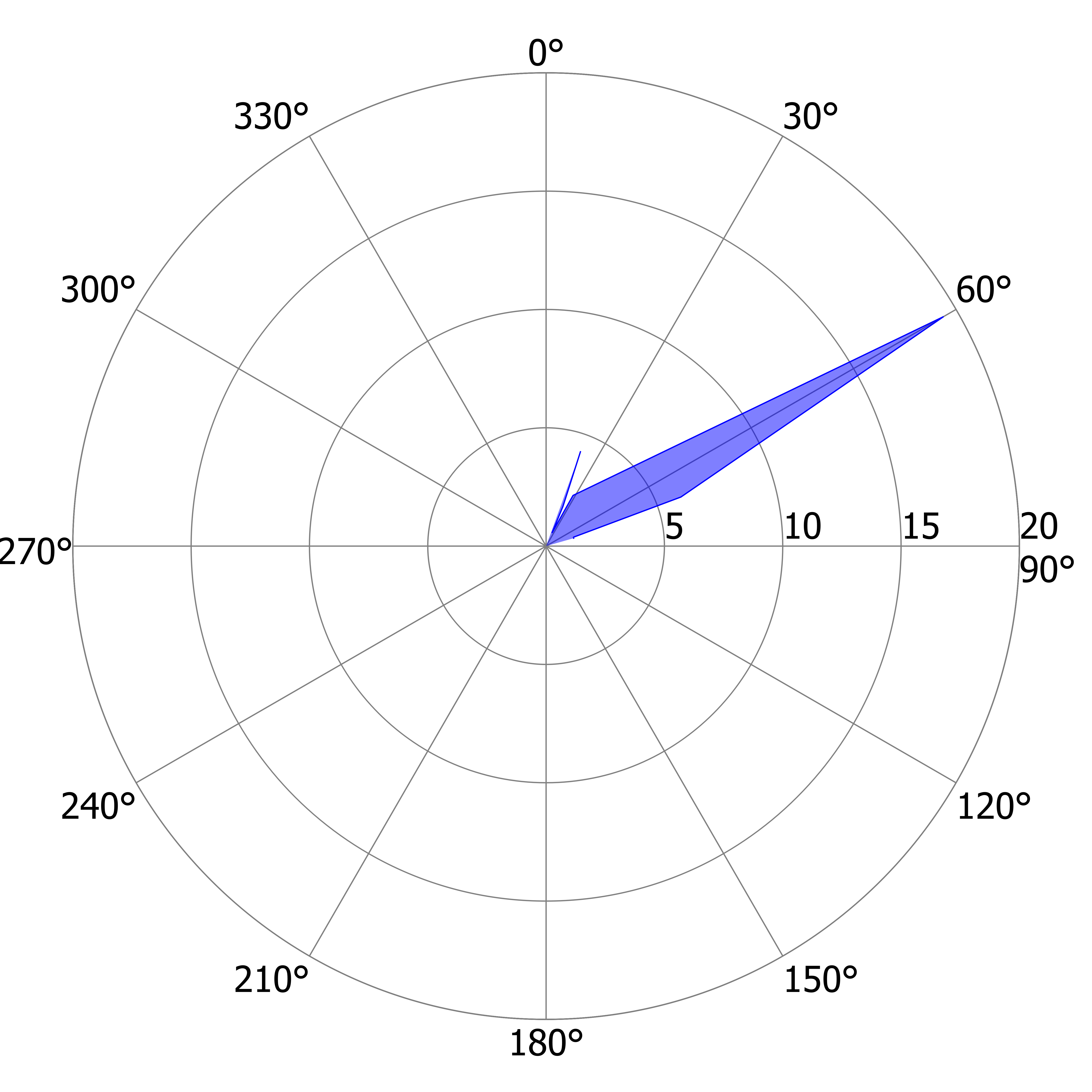

### L_DS_j.tiff

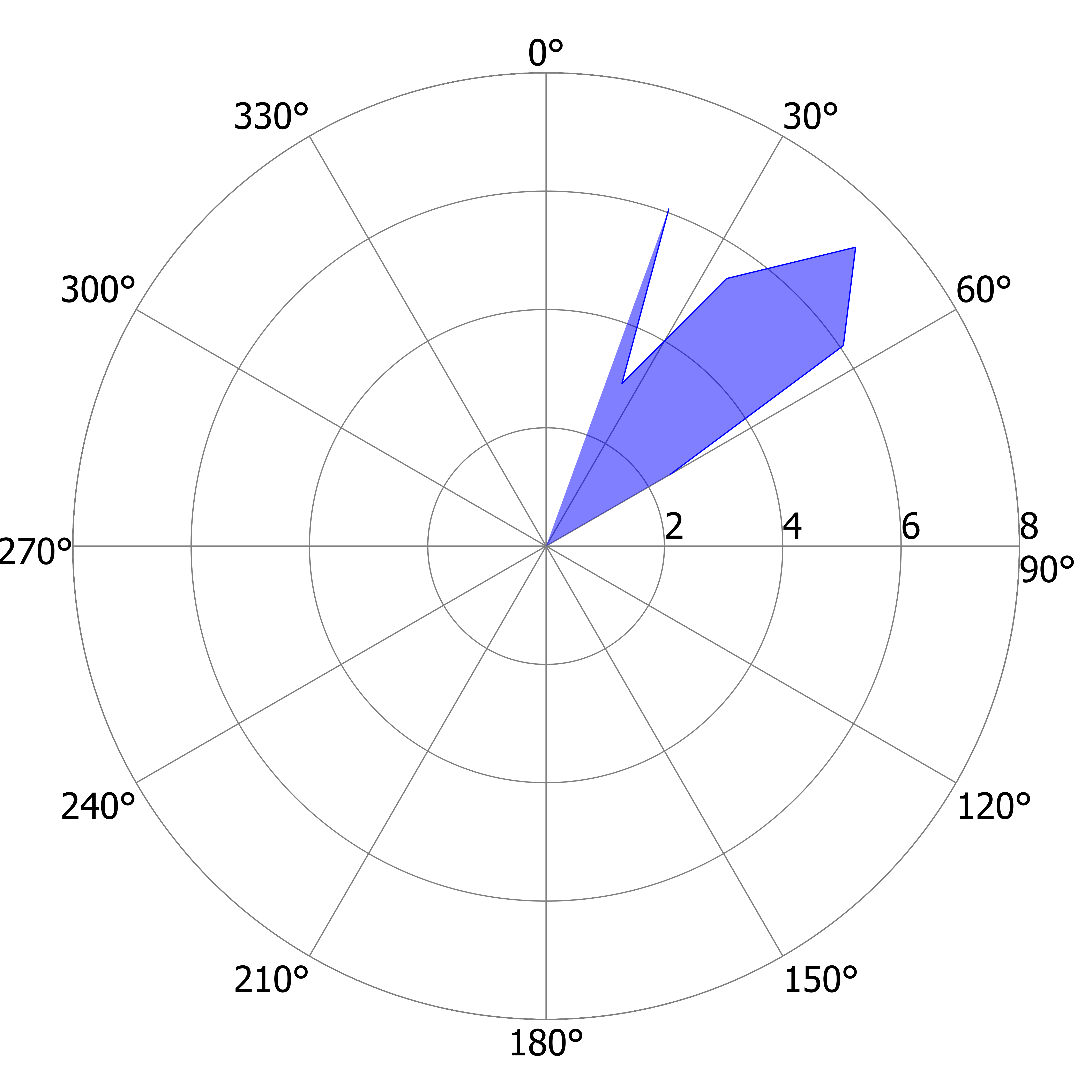

### L_DS_k.tiff

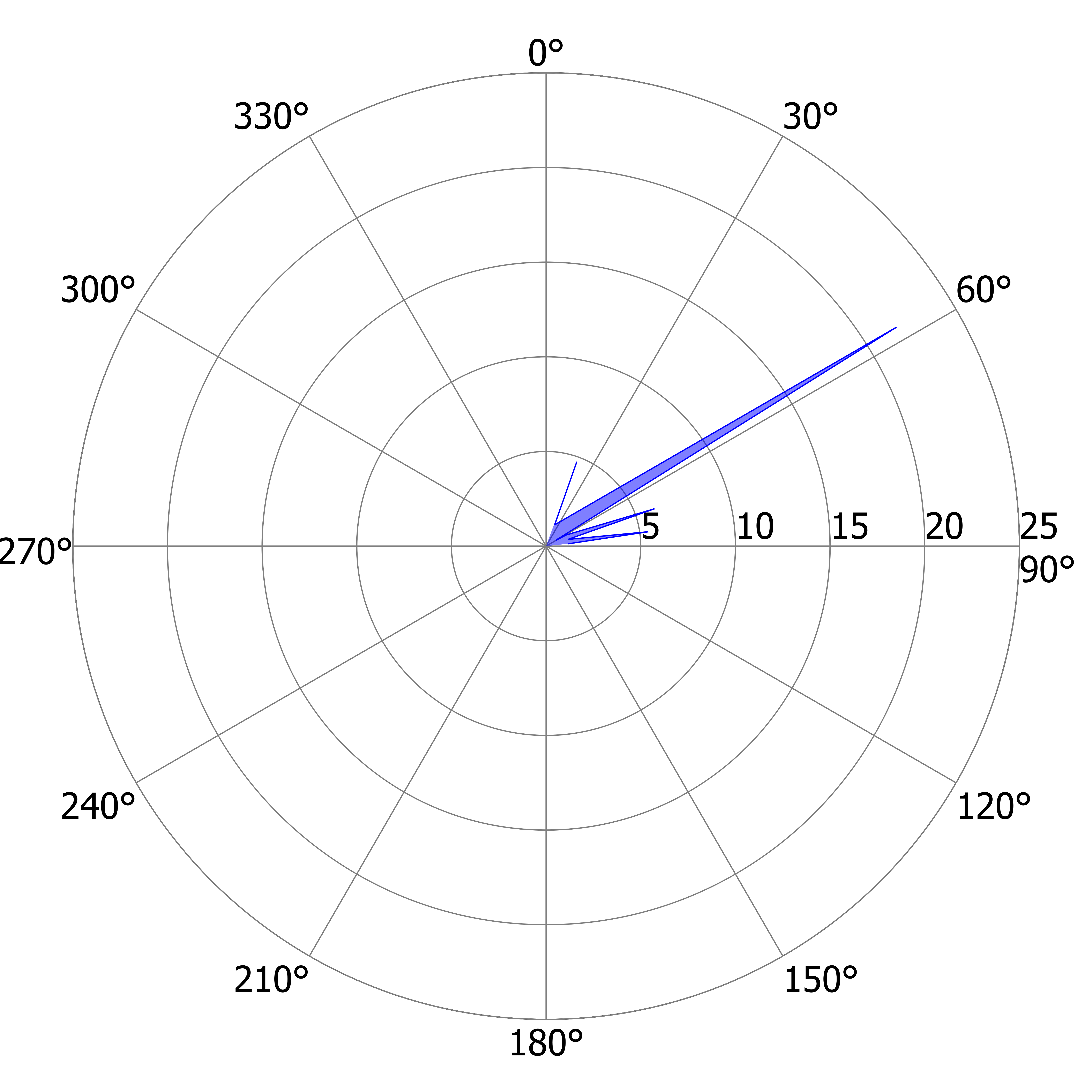

### L_DS_l.tiff

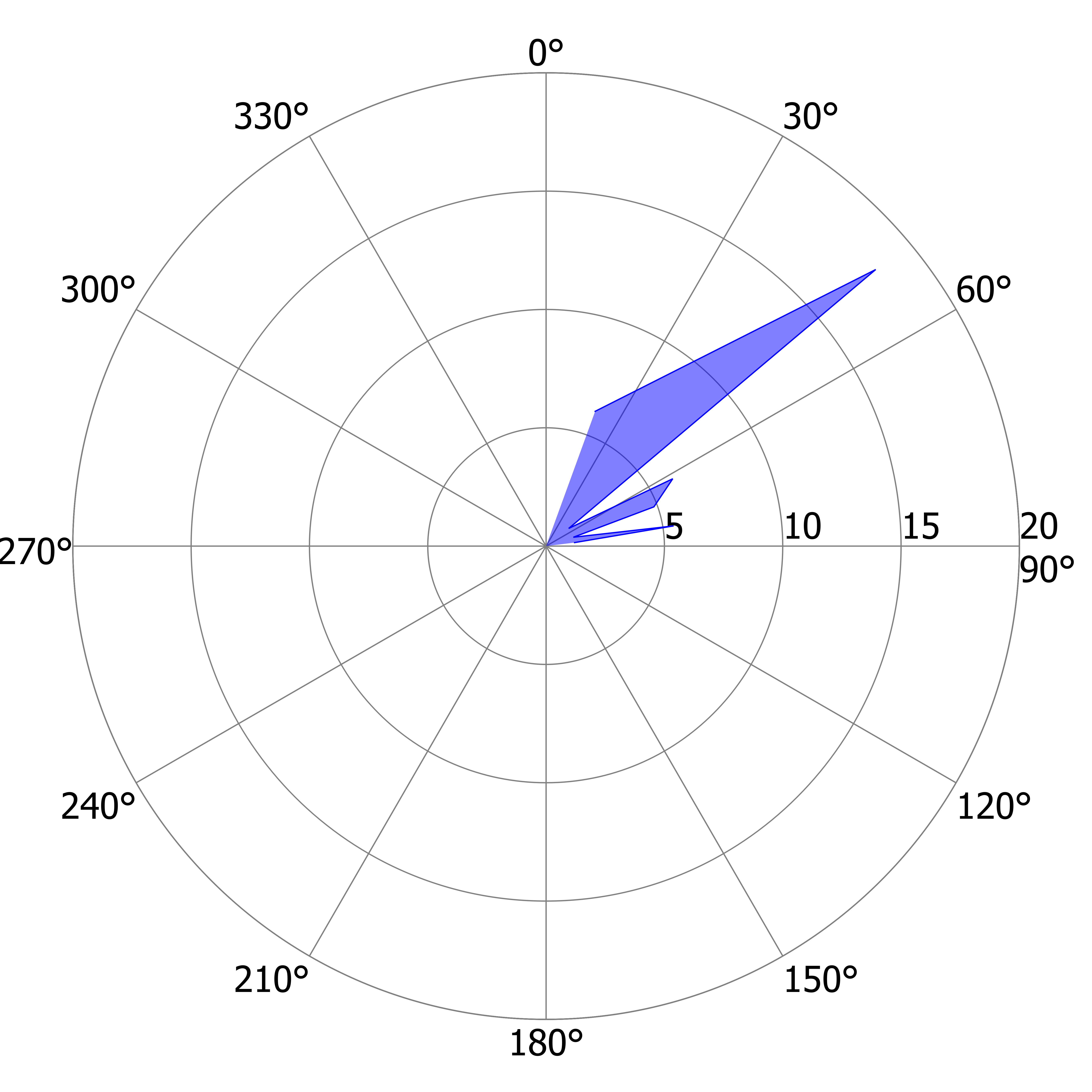

### L_DS_m.tiff

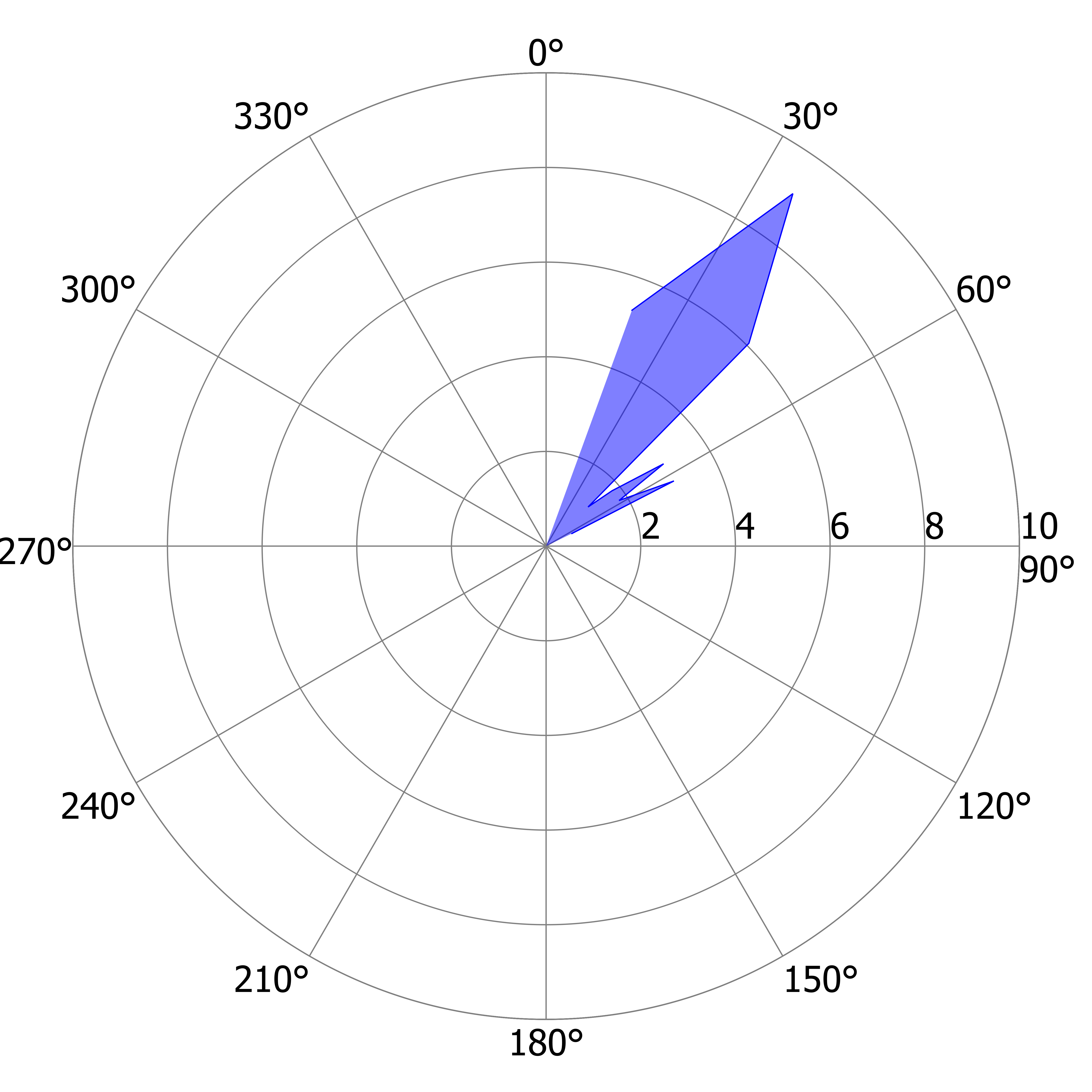

### L_DS_n.tiff

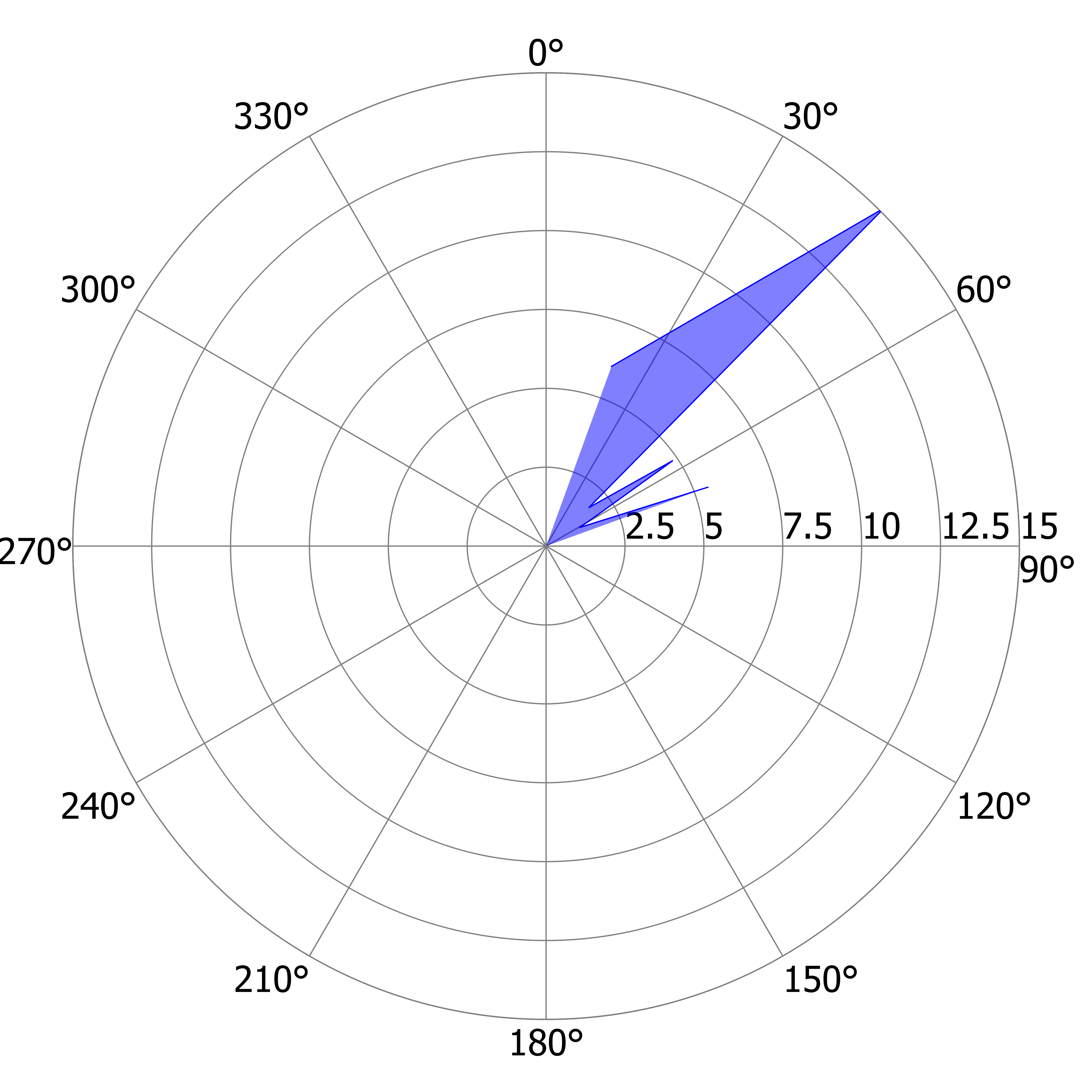

### L_DS_o.tiff

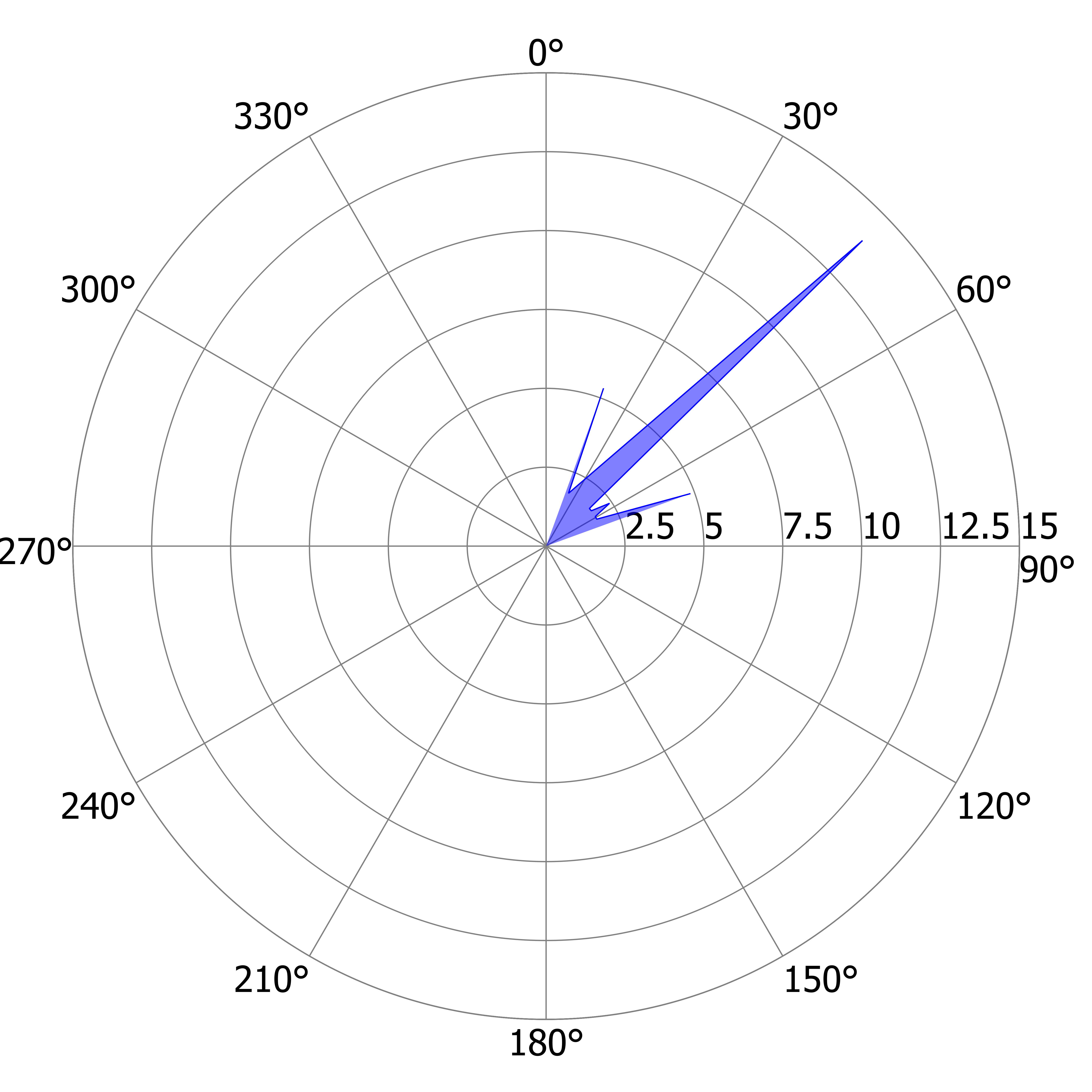

### L_DS_p.tiff

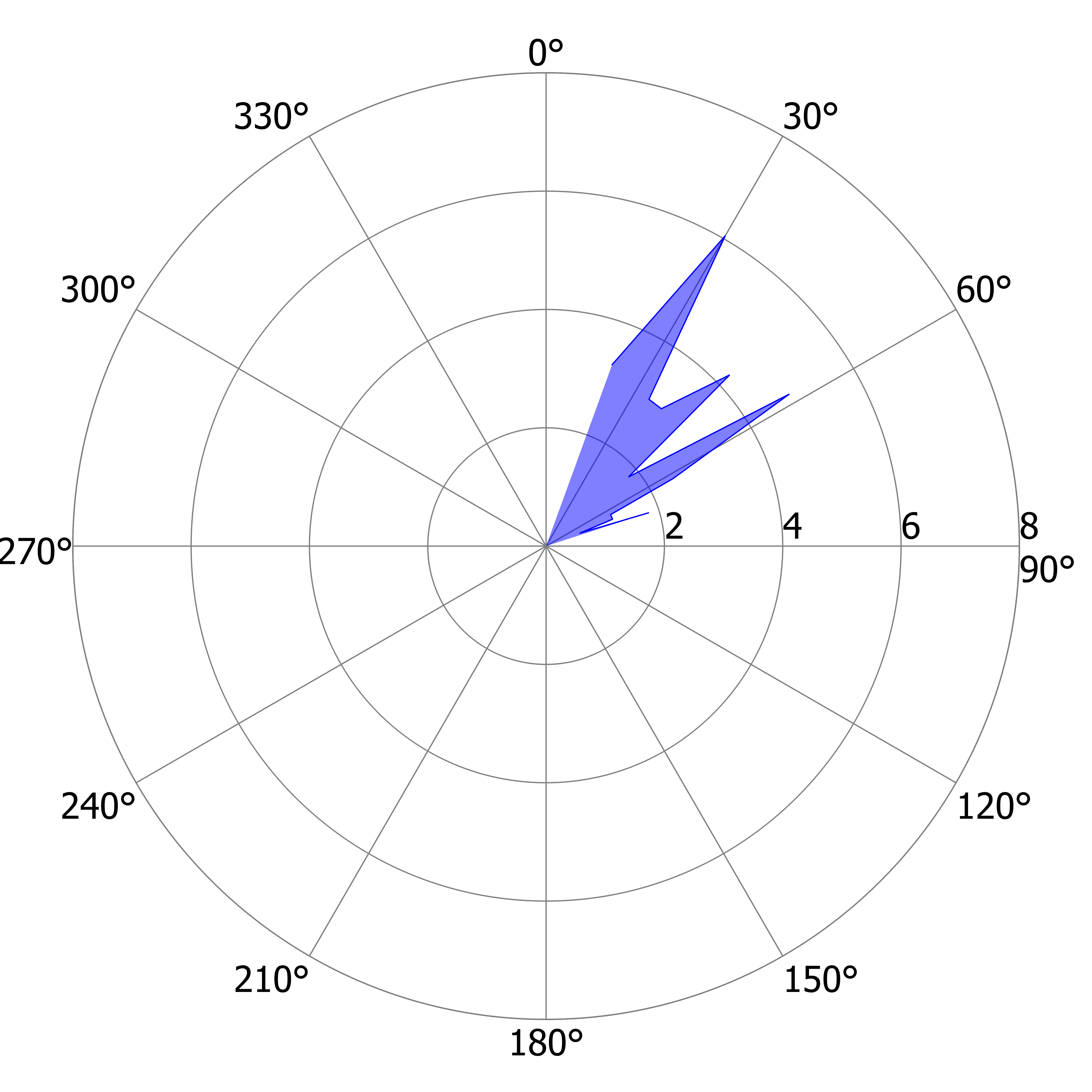

### L_DS_q.tiff

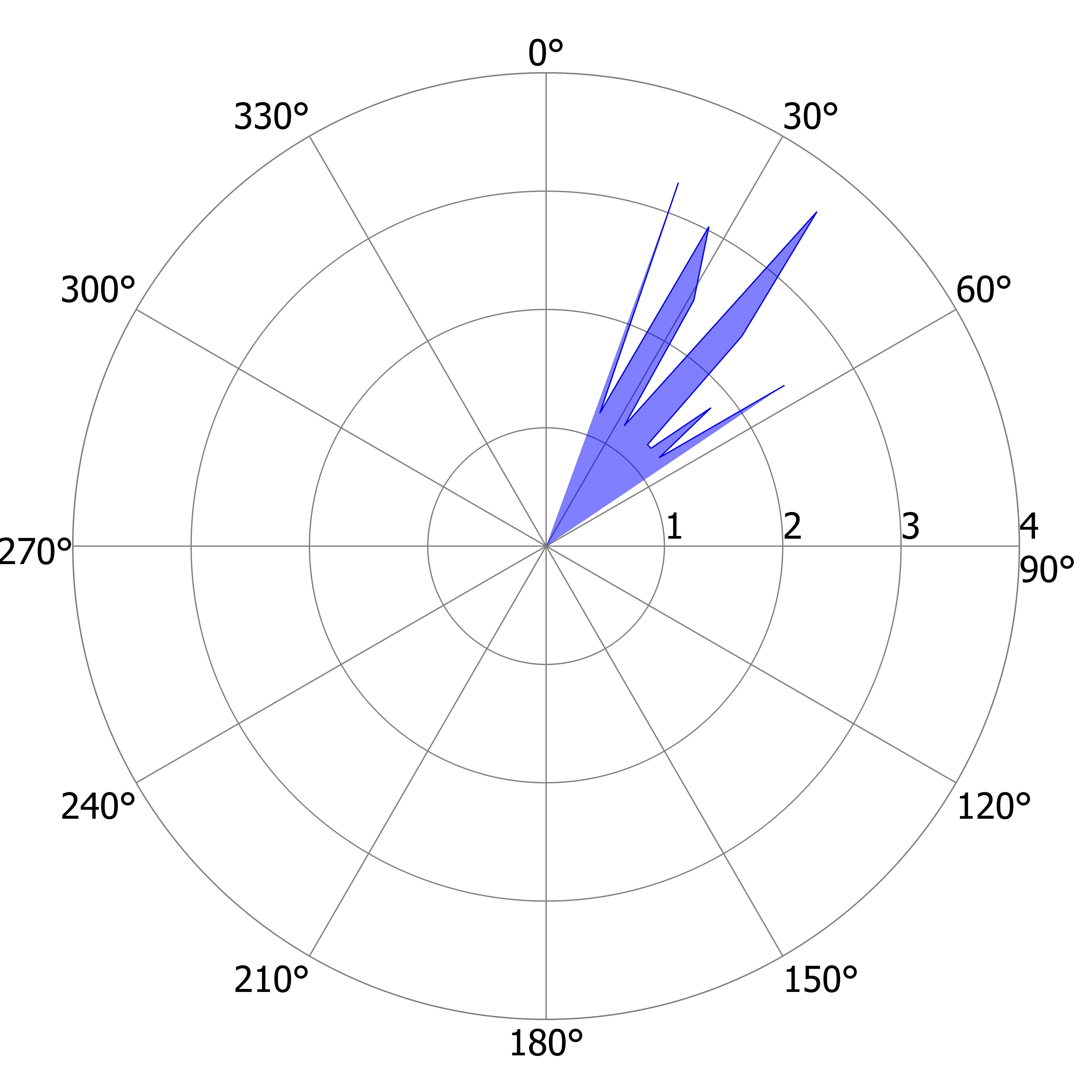

### L_DS_r.tiff

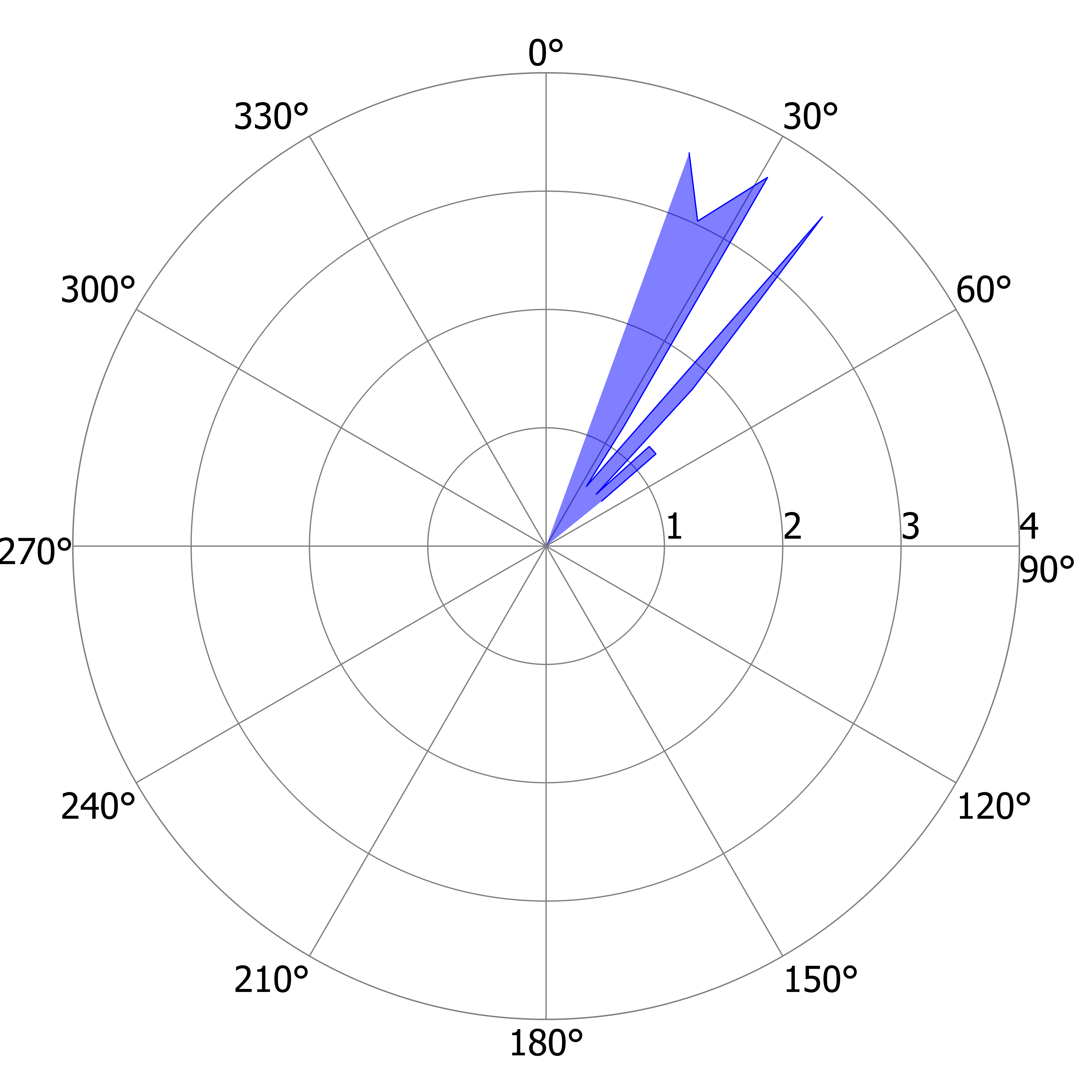

### L_DS_s.tiff

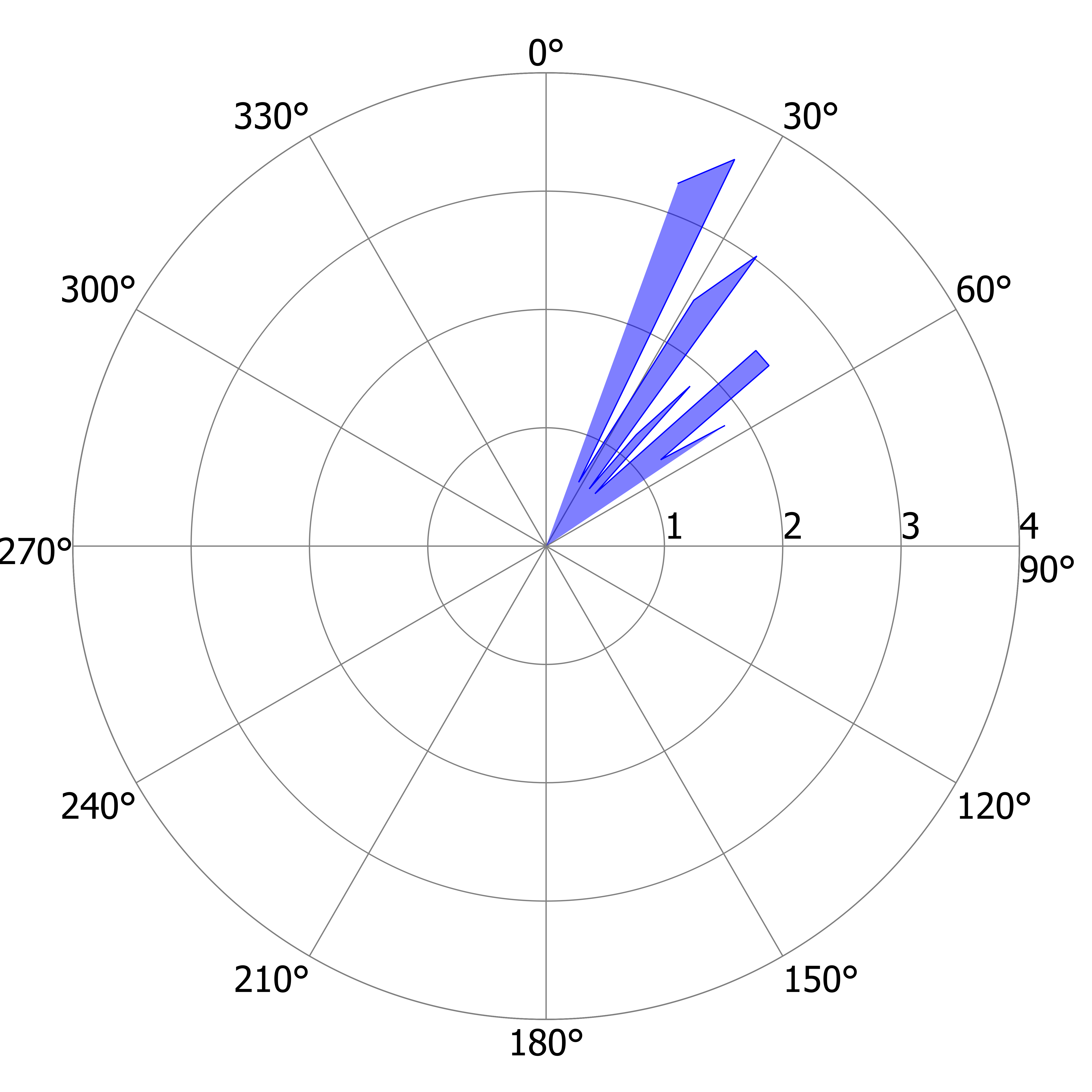

### L_DS_t.tiff

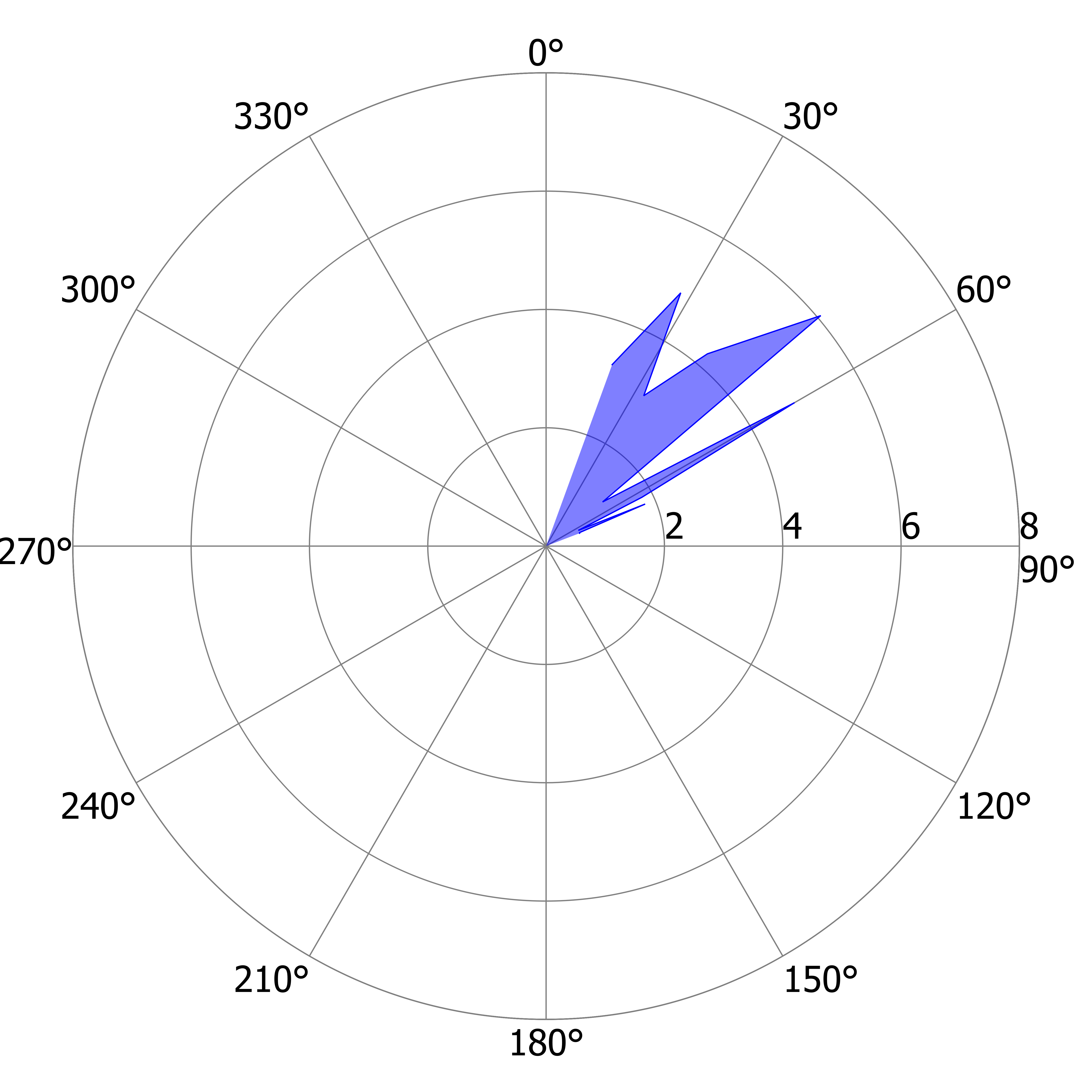

### L_US_a.tiff

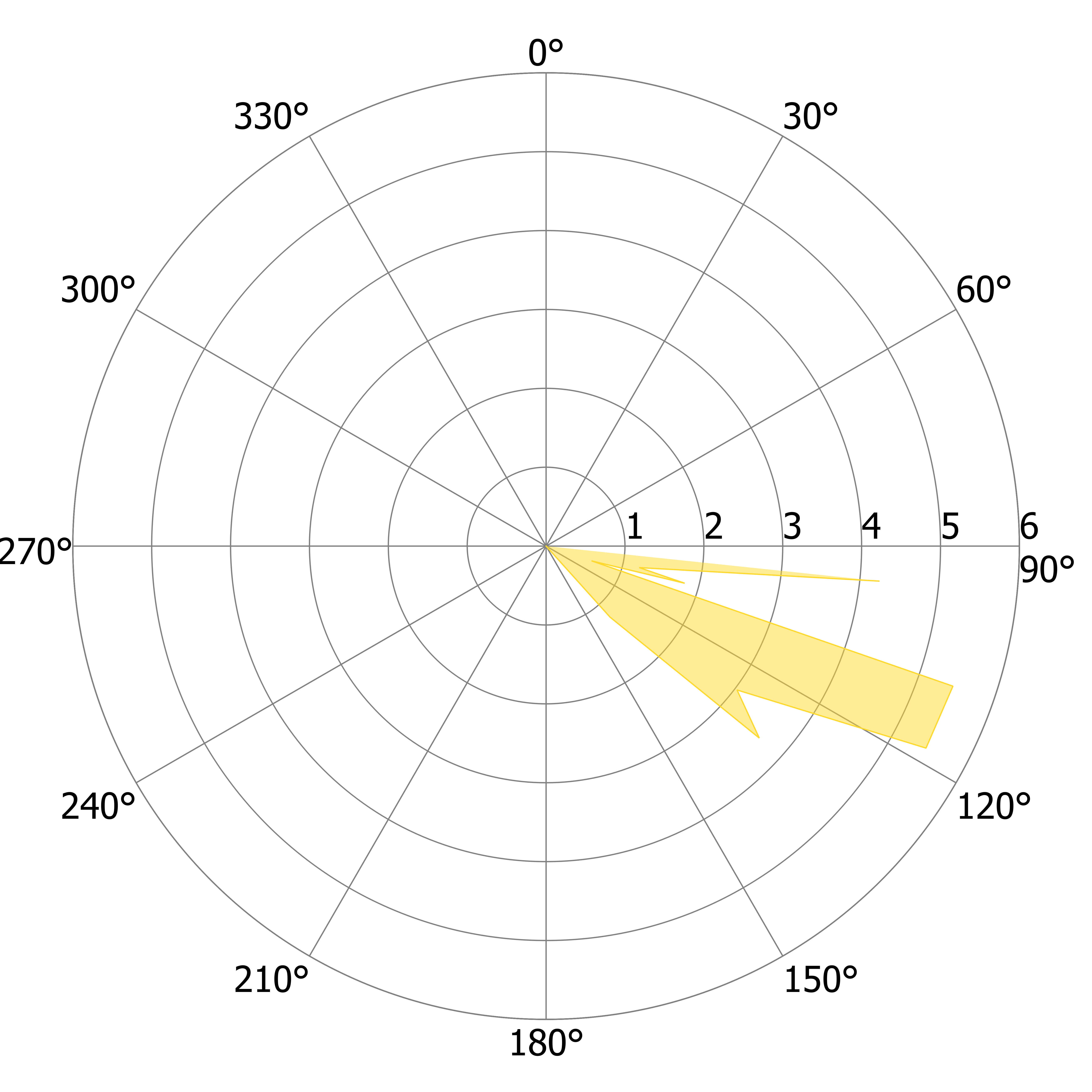

### L_US_a.tiff

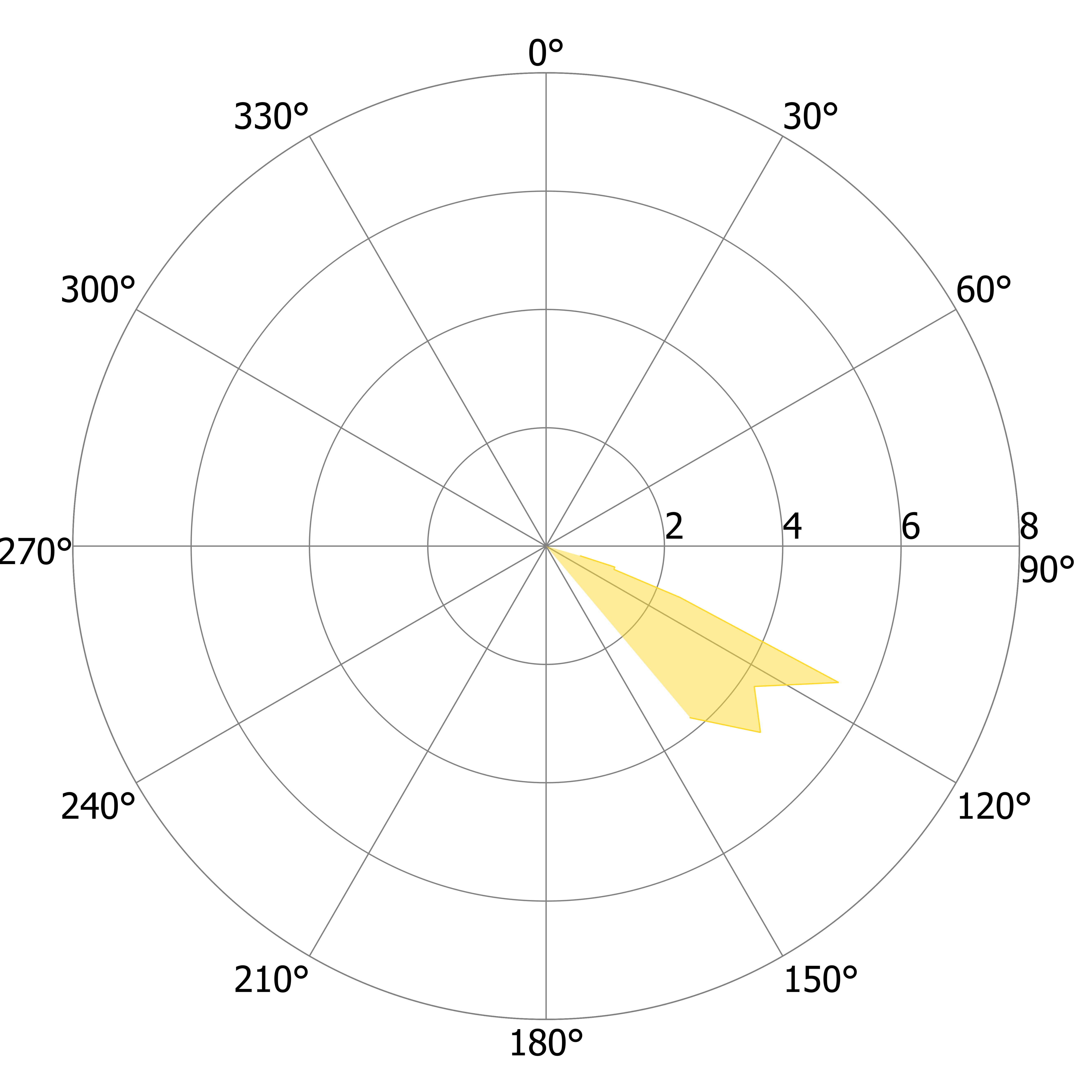

### L_US_b.tiff

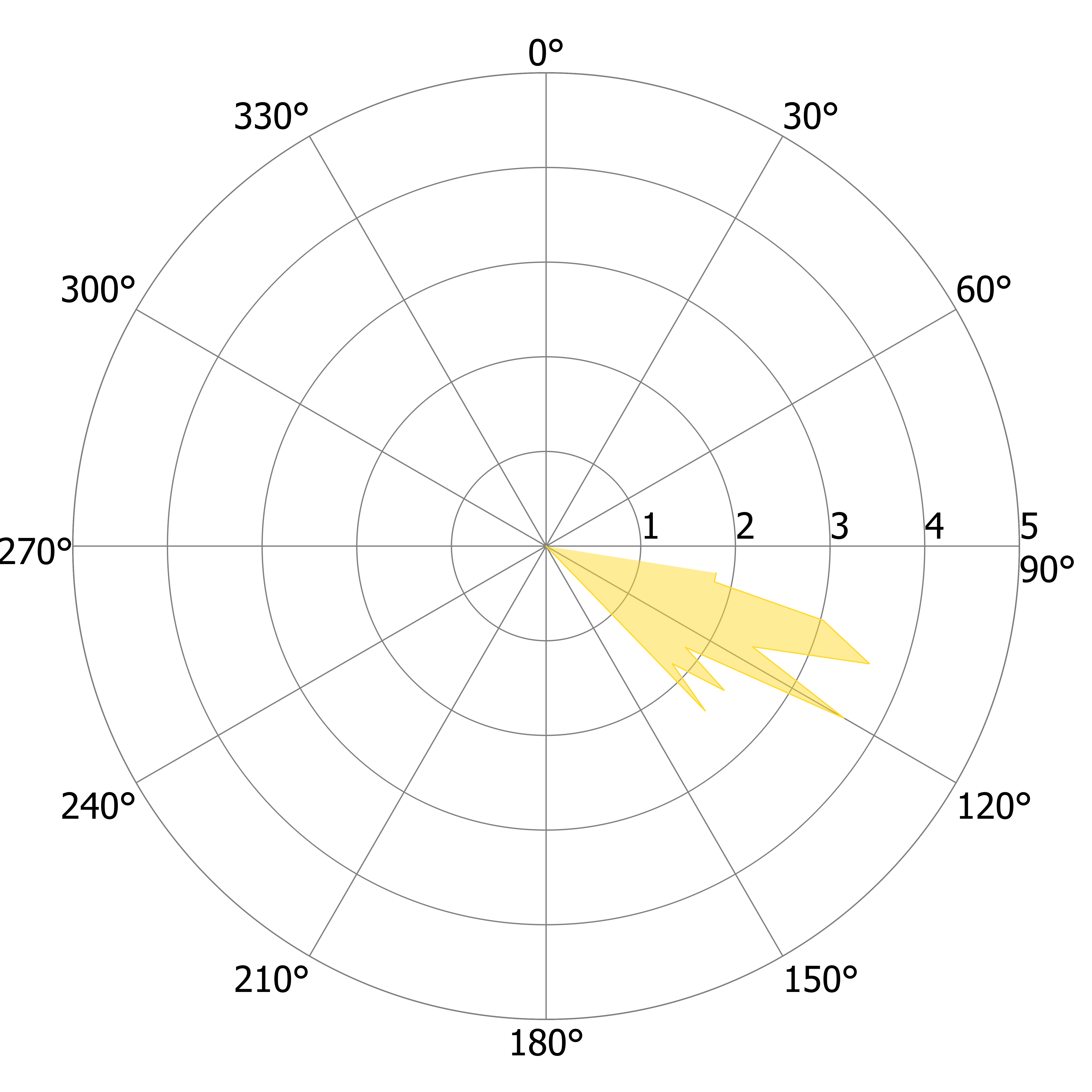

### L_US_b.tiff

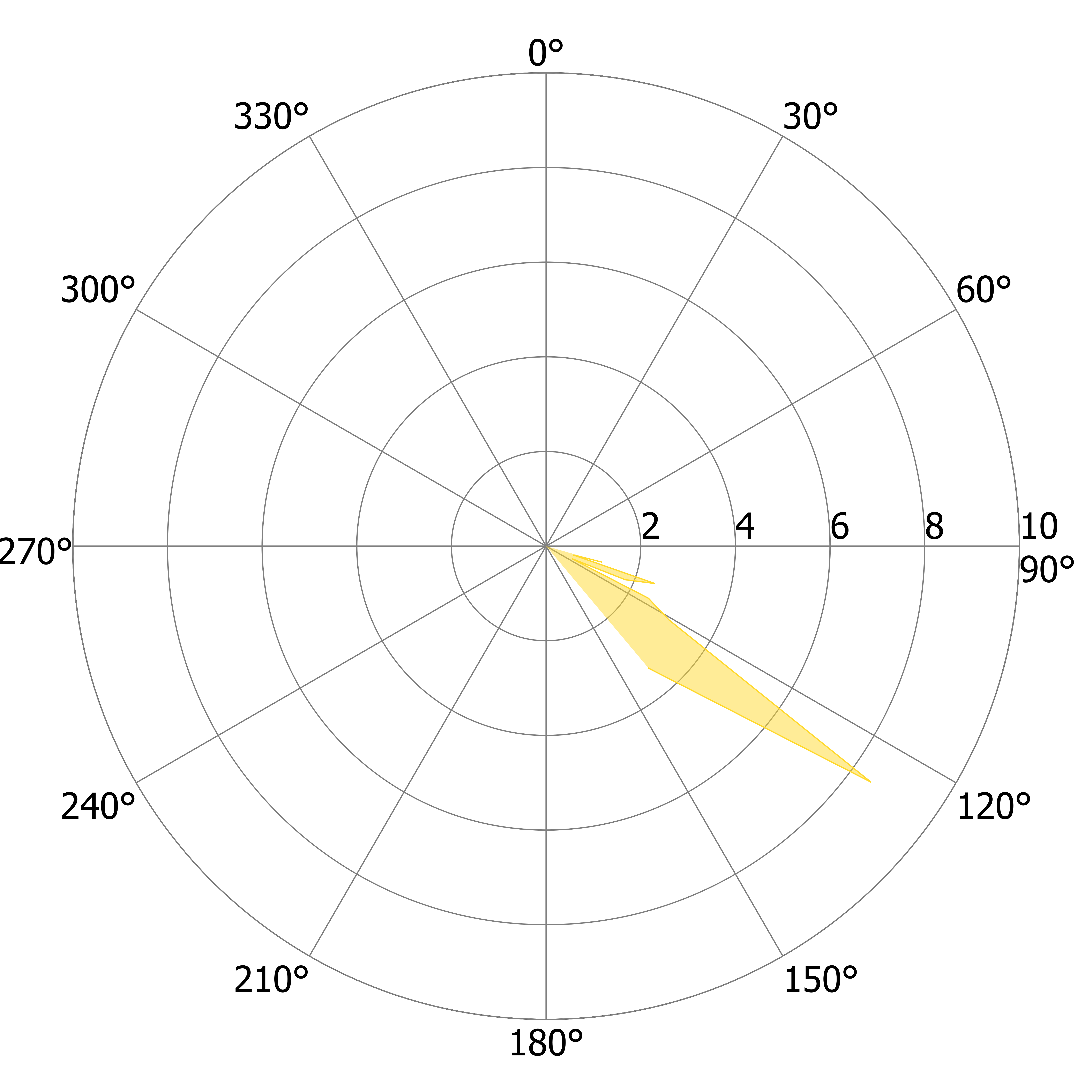

### L_US_c.tiff

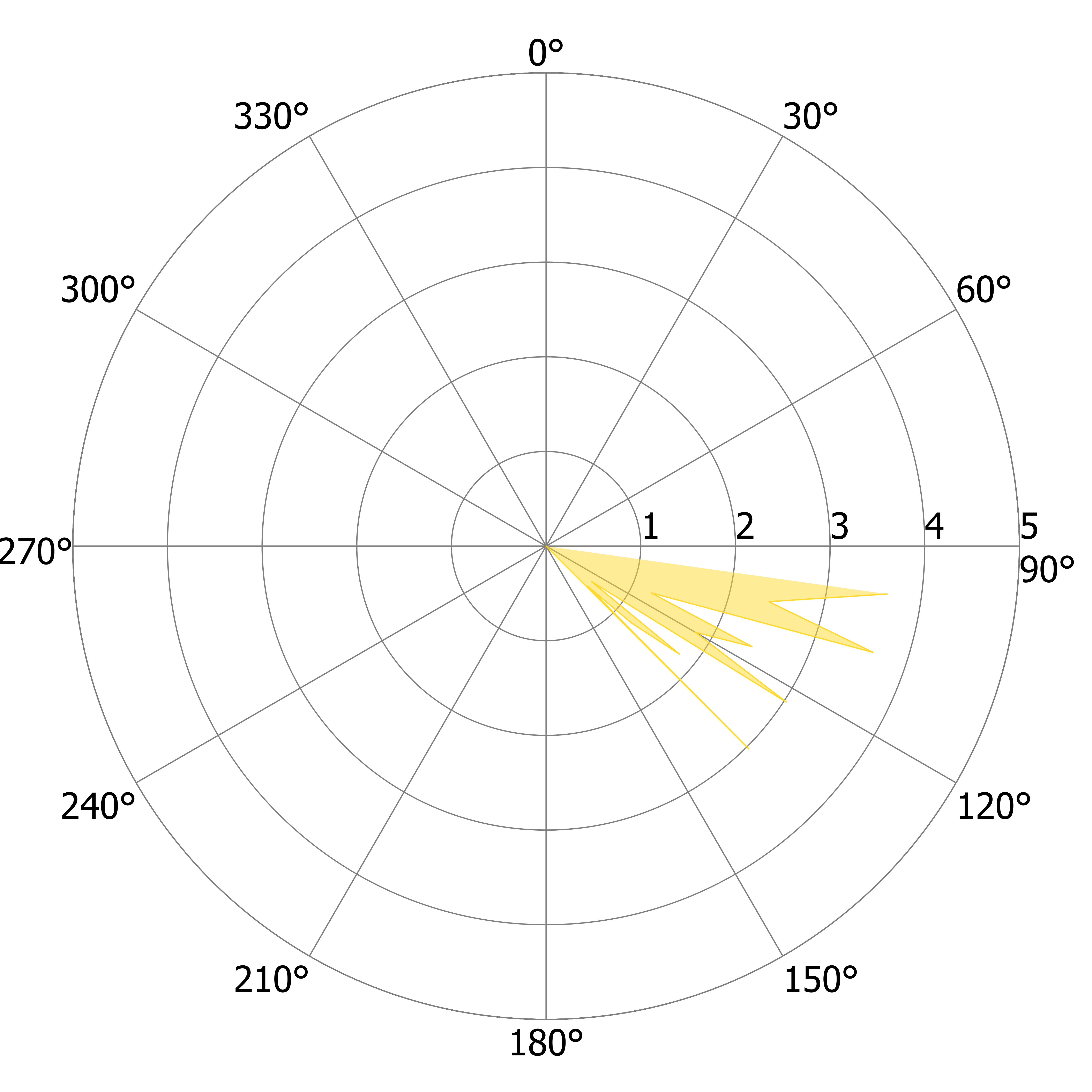

### L_US_c.tiff

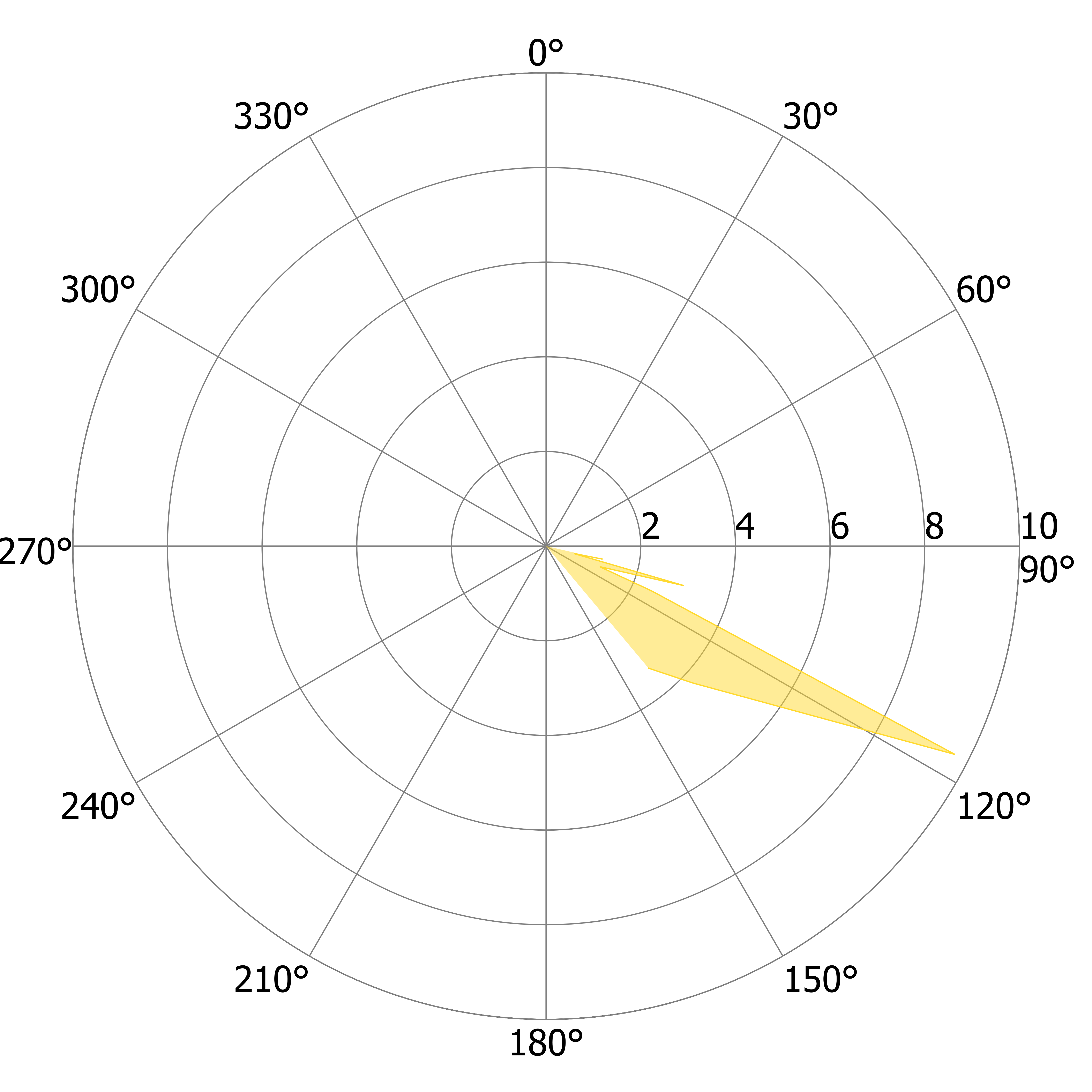

### L_US_d.tiff

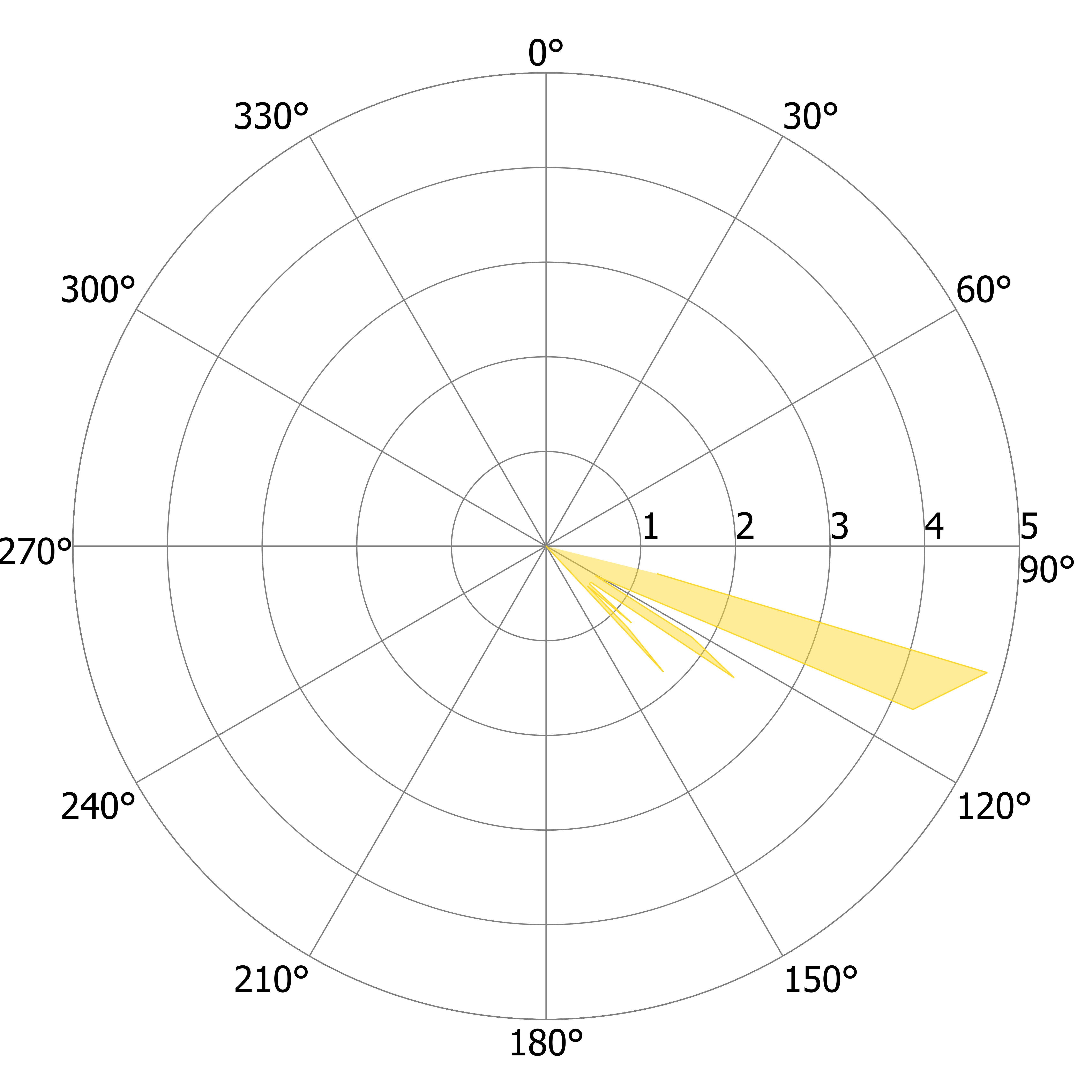

### L_US_d.tiff

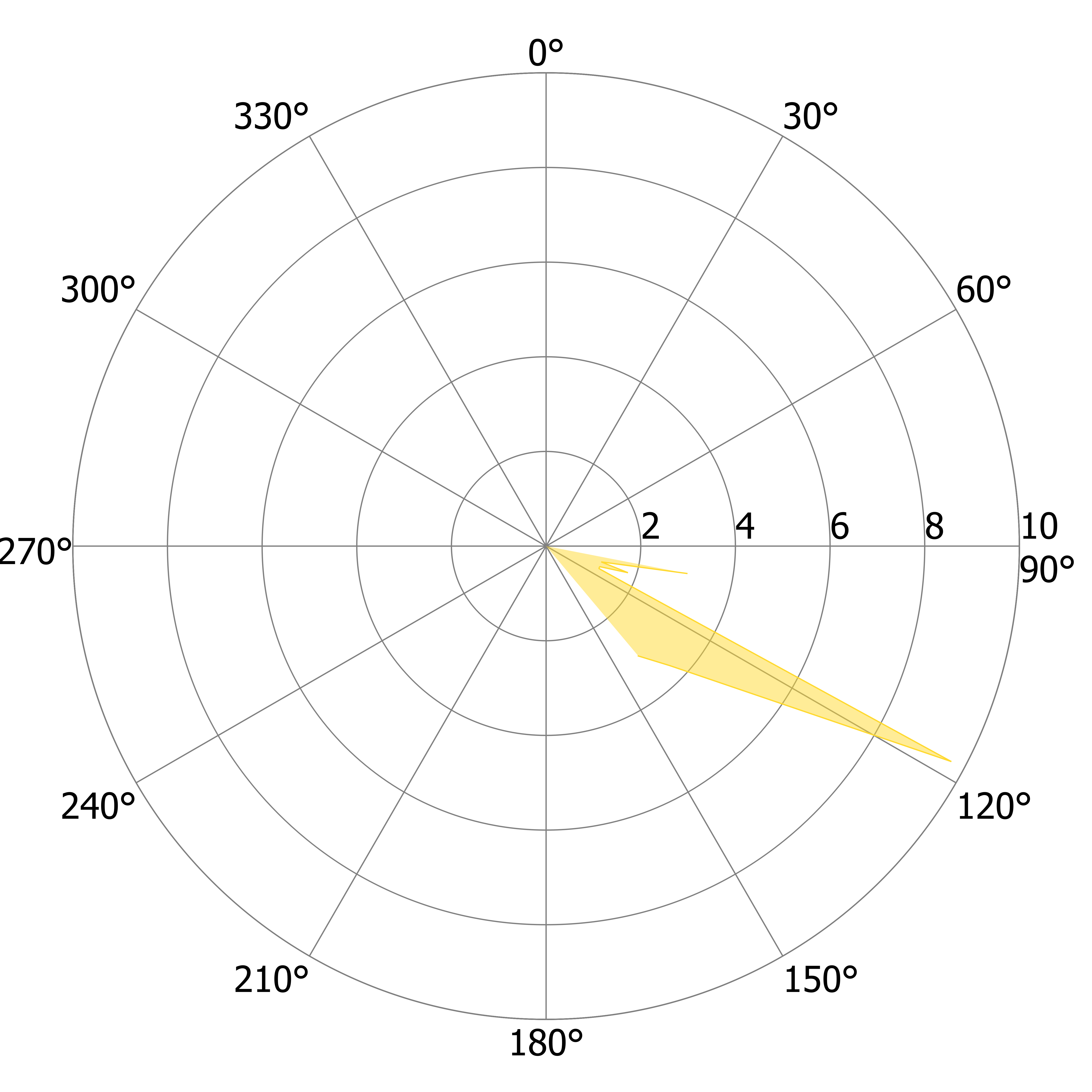

### L_US_e.tiff

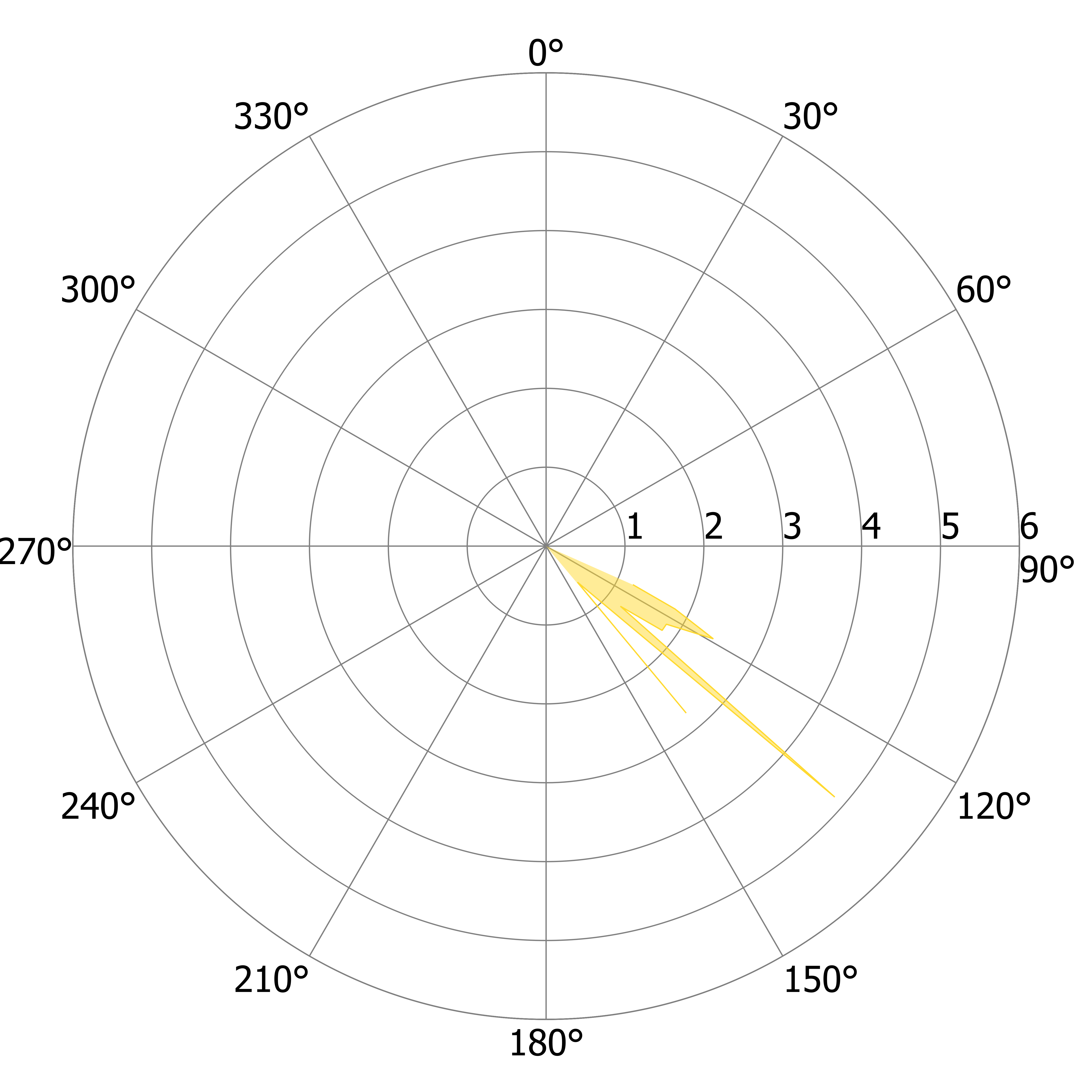

### L_US_f.tiff

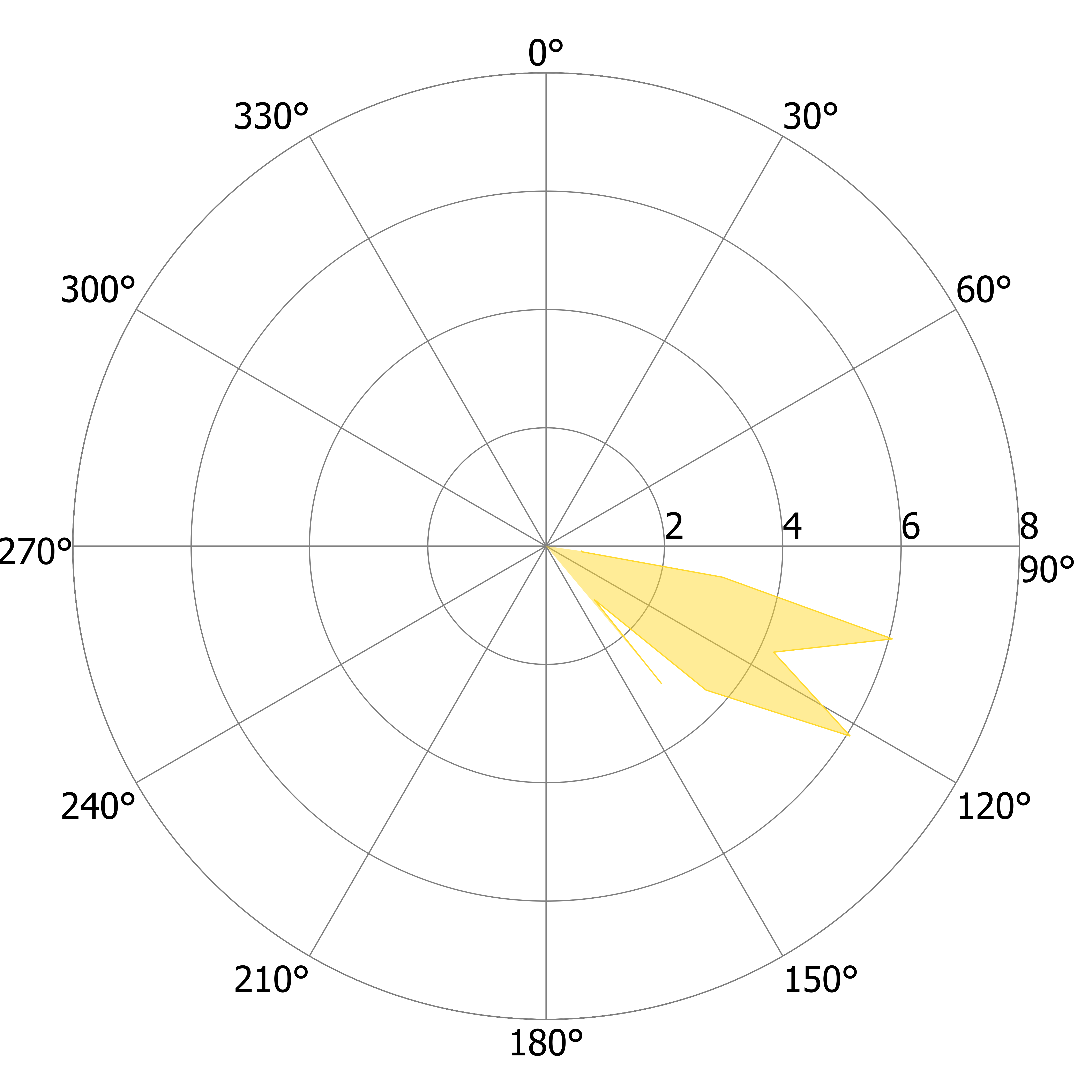

### L_US_g.tiff

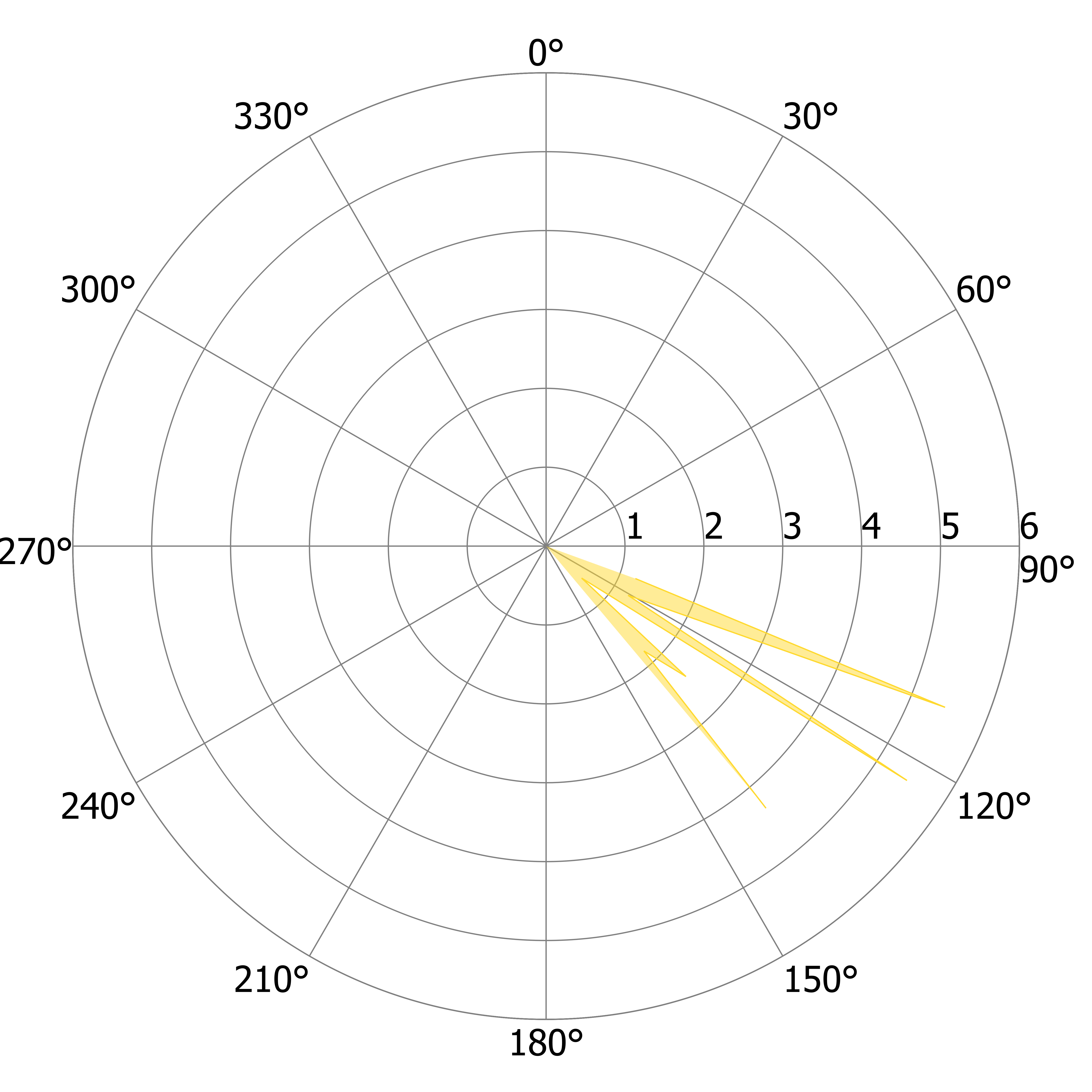

### L_US_h.tiff

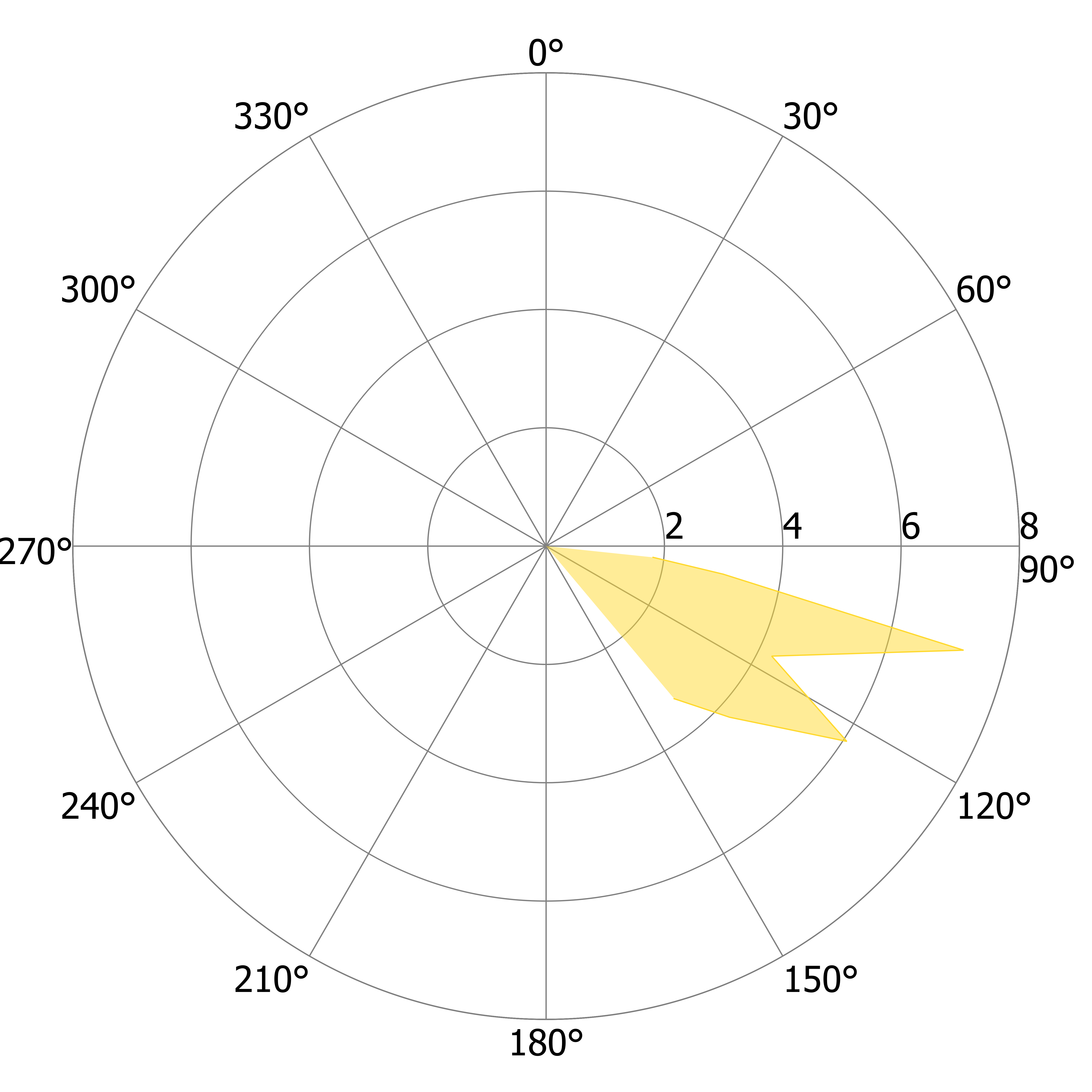

### L_US_i.tiff

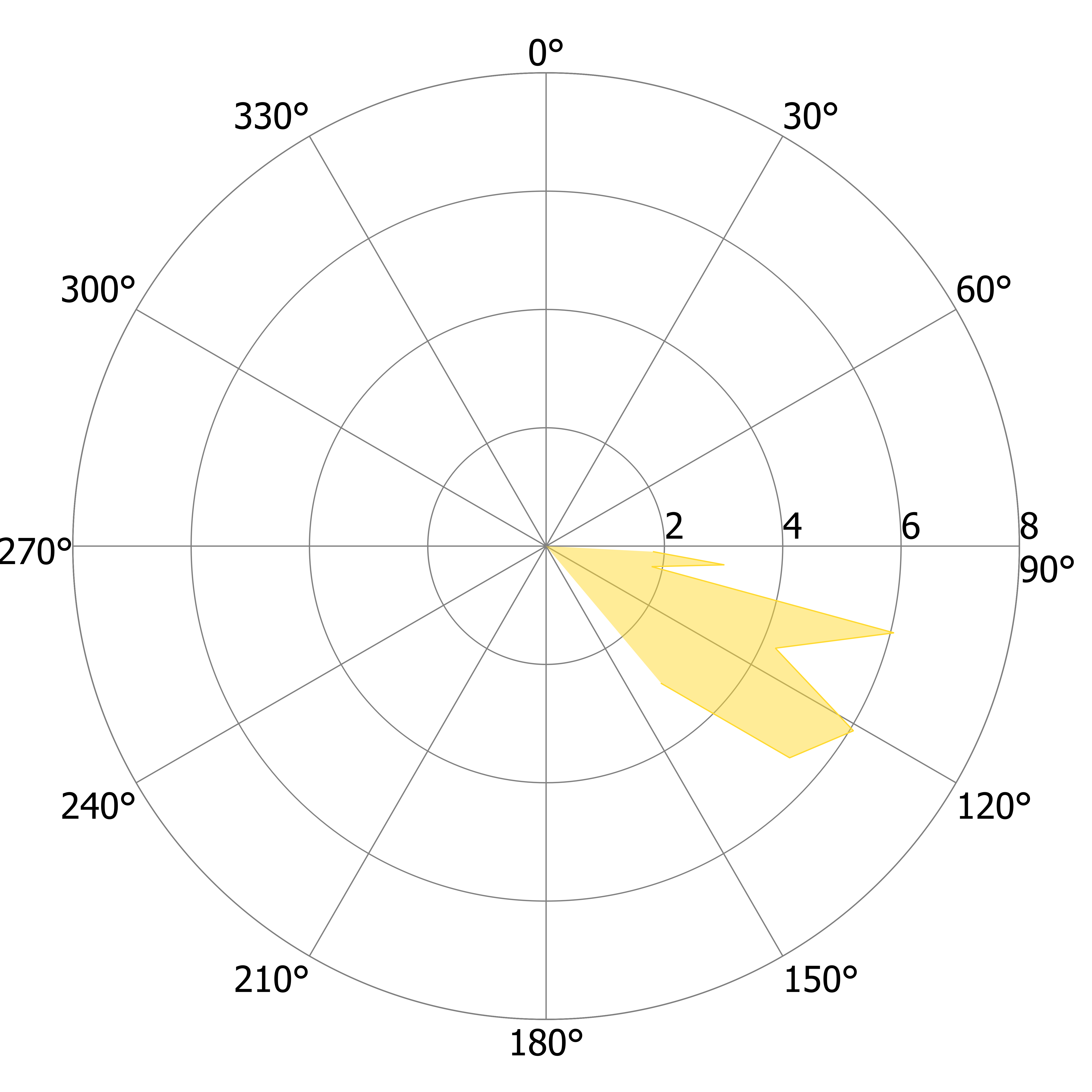

### L_US_j.tiff

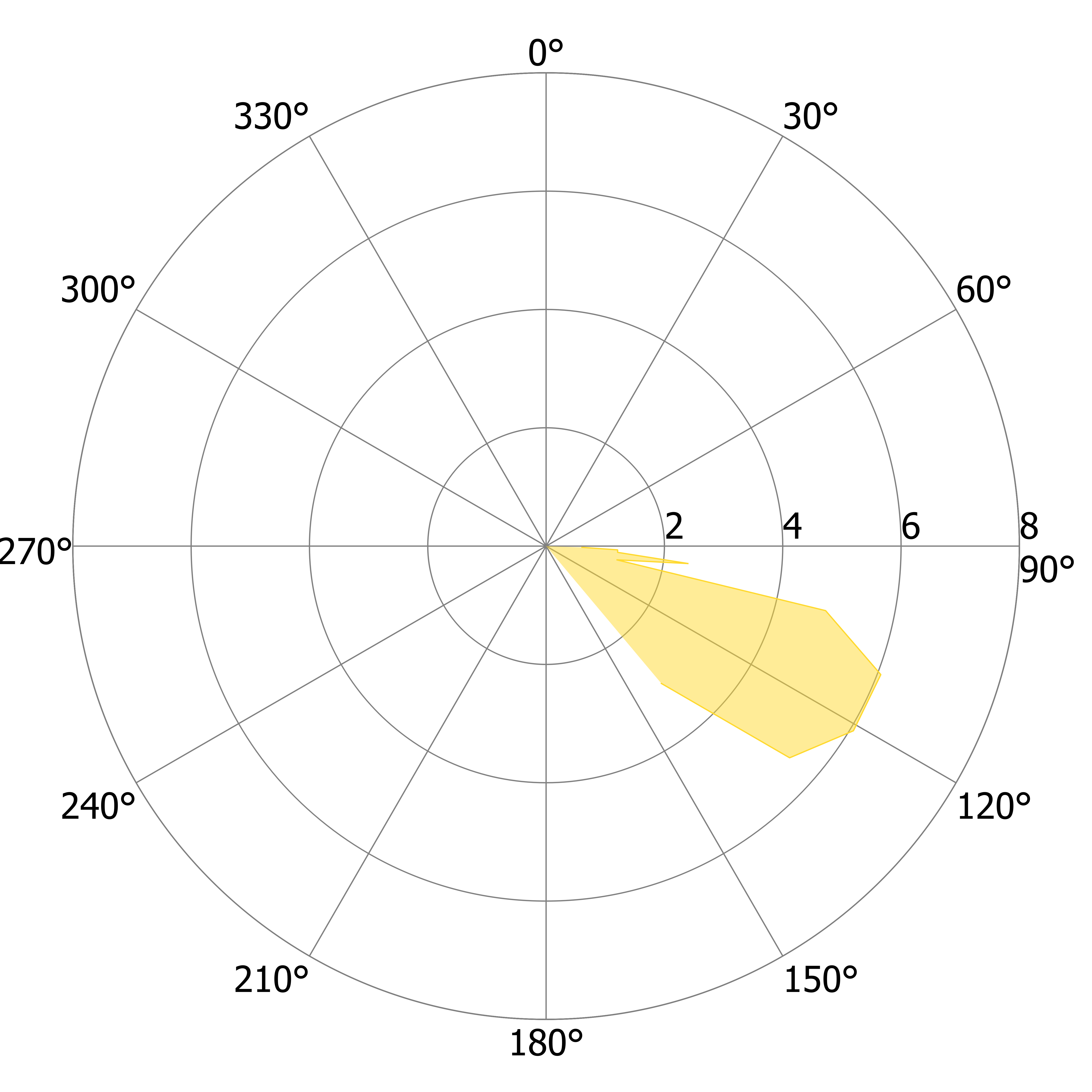

### L_US_k.tiff

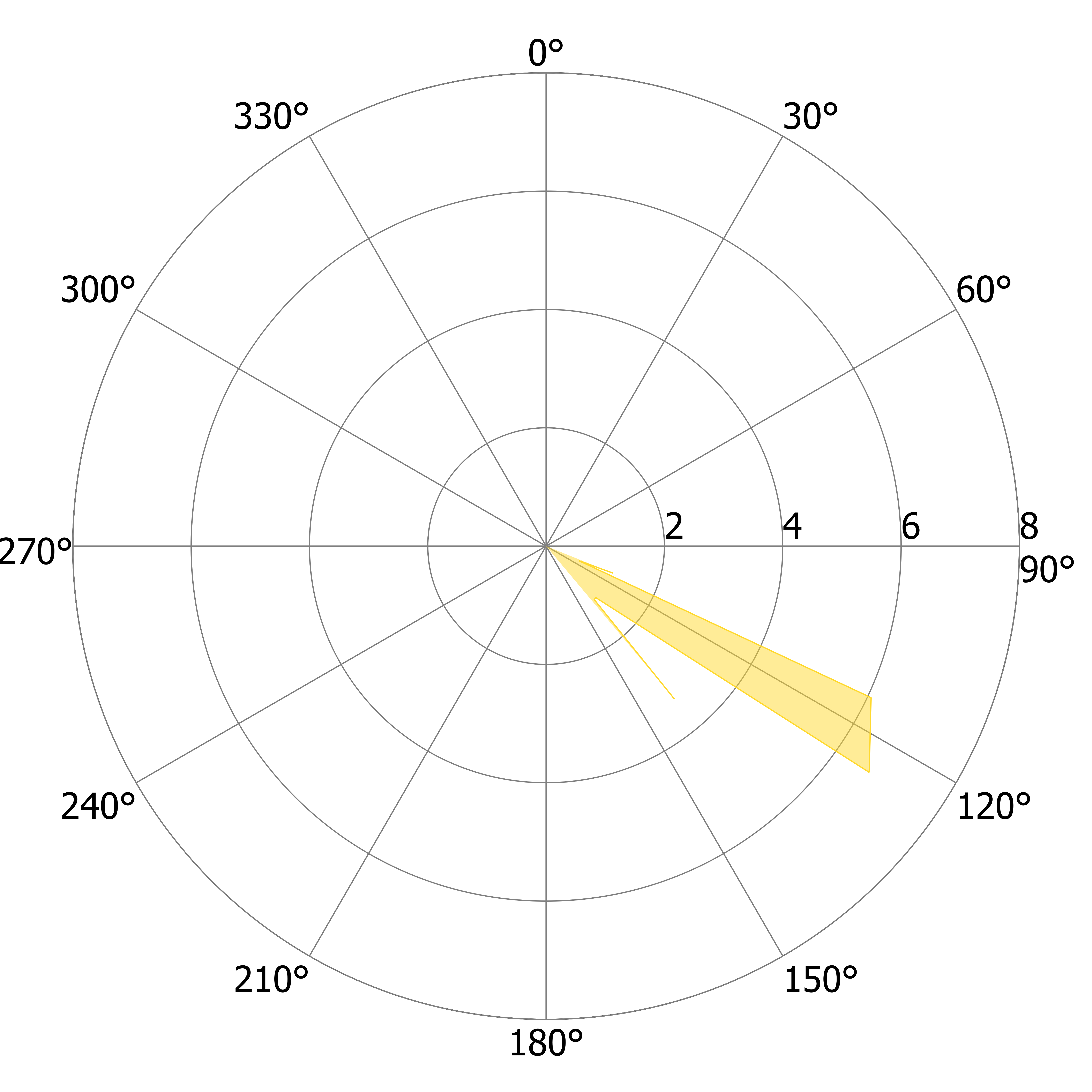

### L_US_l.tiff

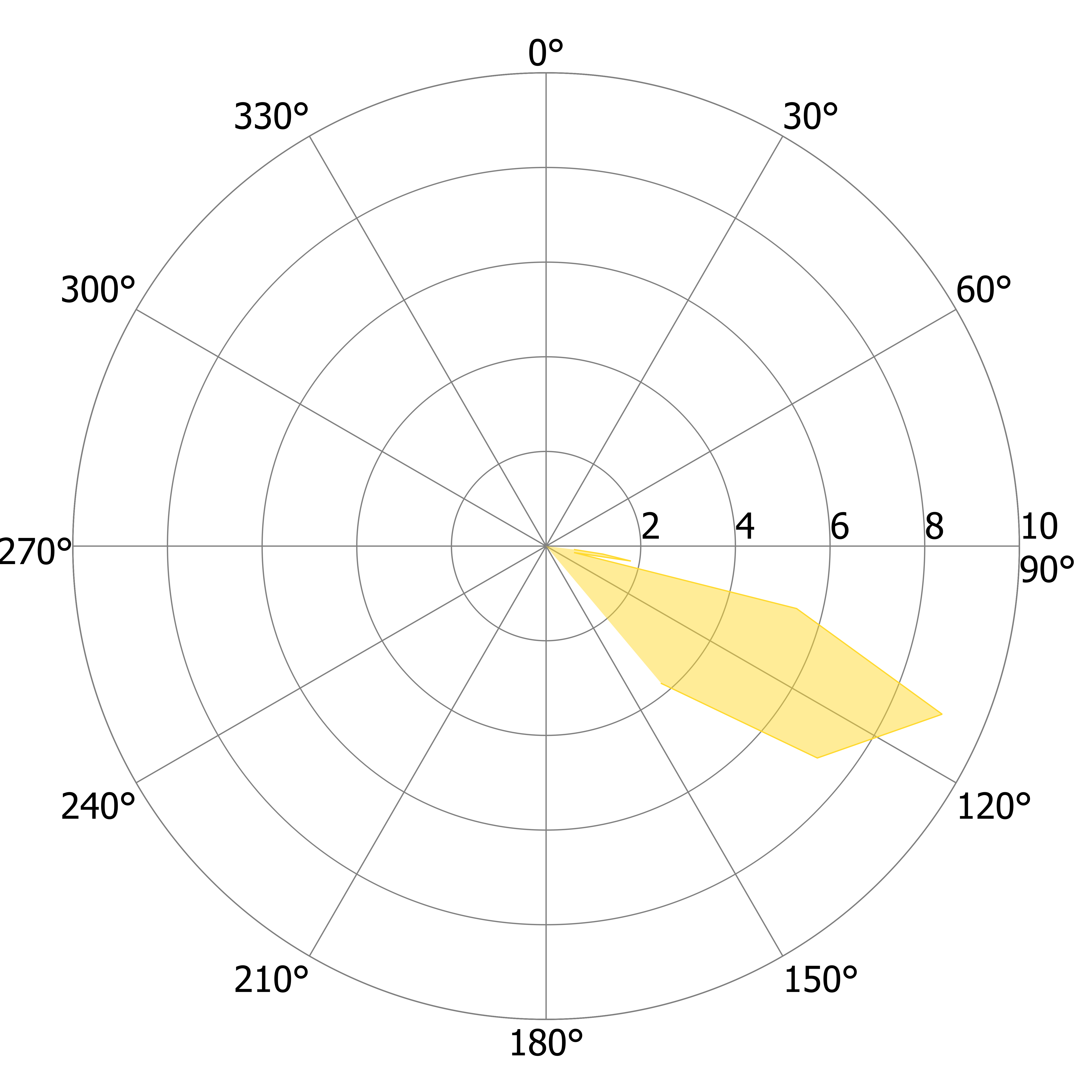

### L_US_m.tiff

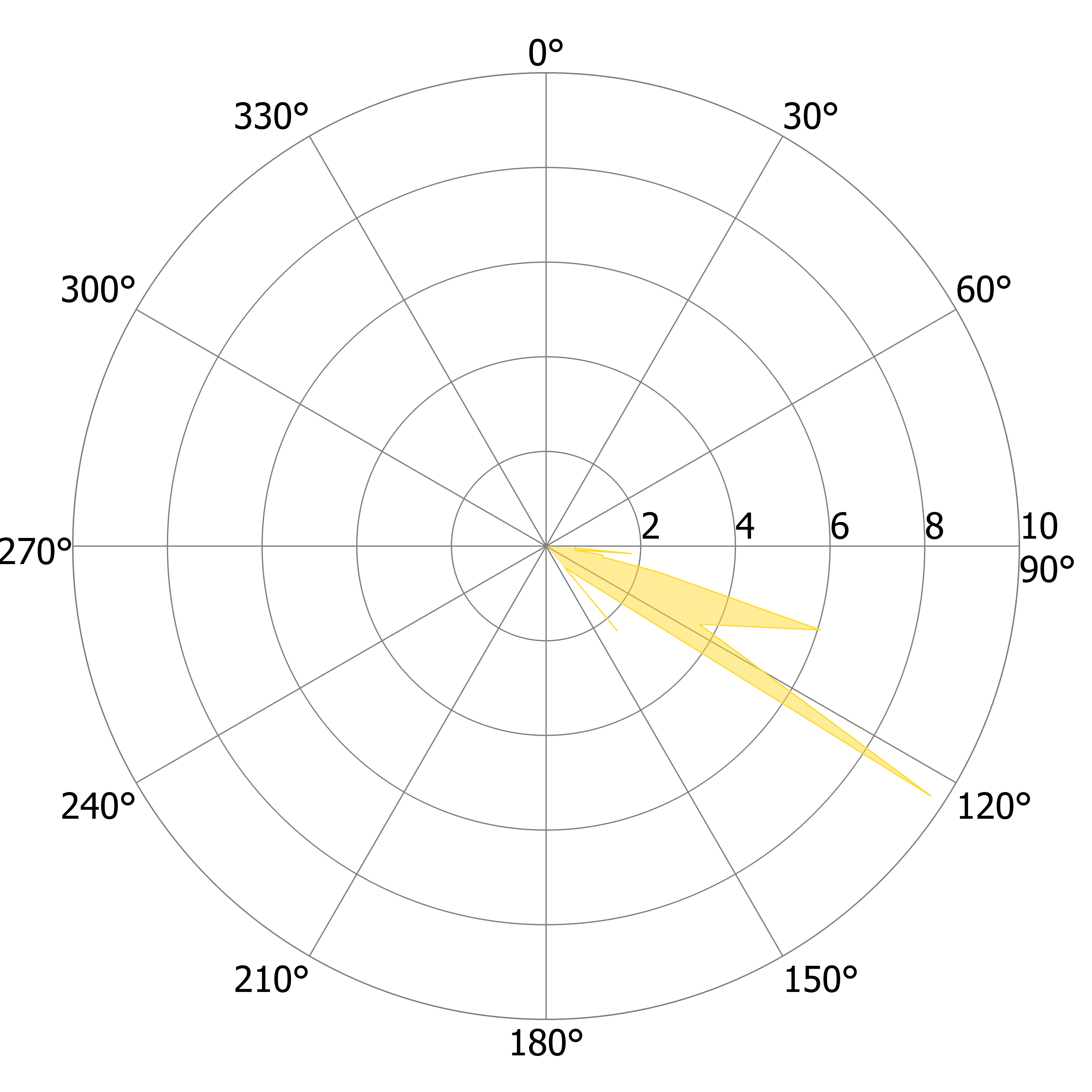

### L_US_n.tiff

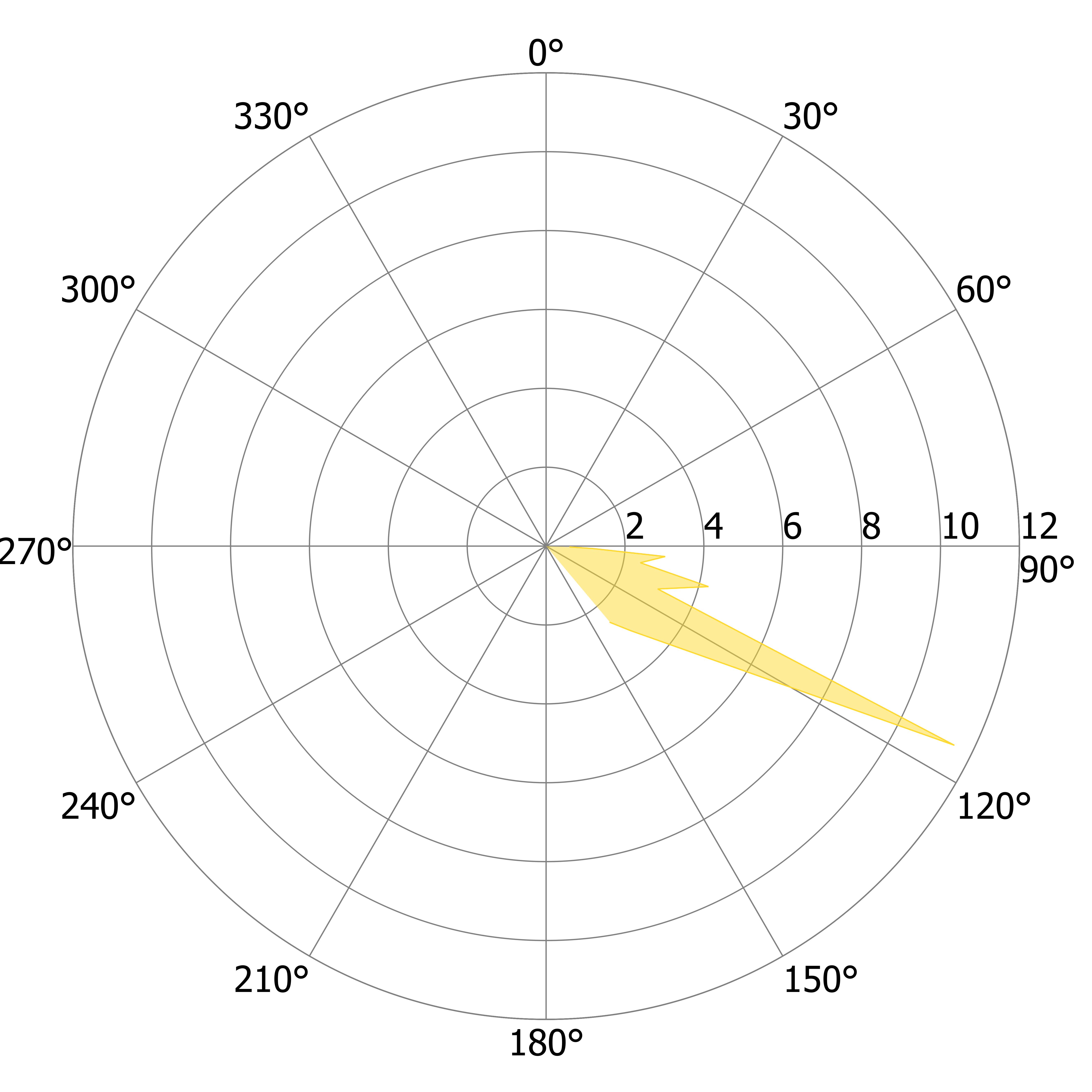

### L_US_o.tiff

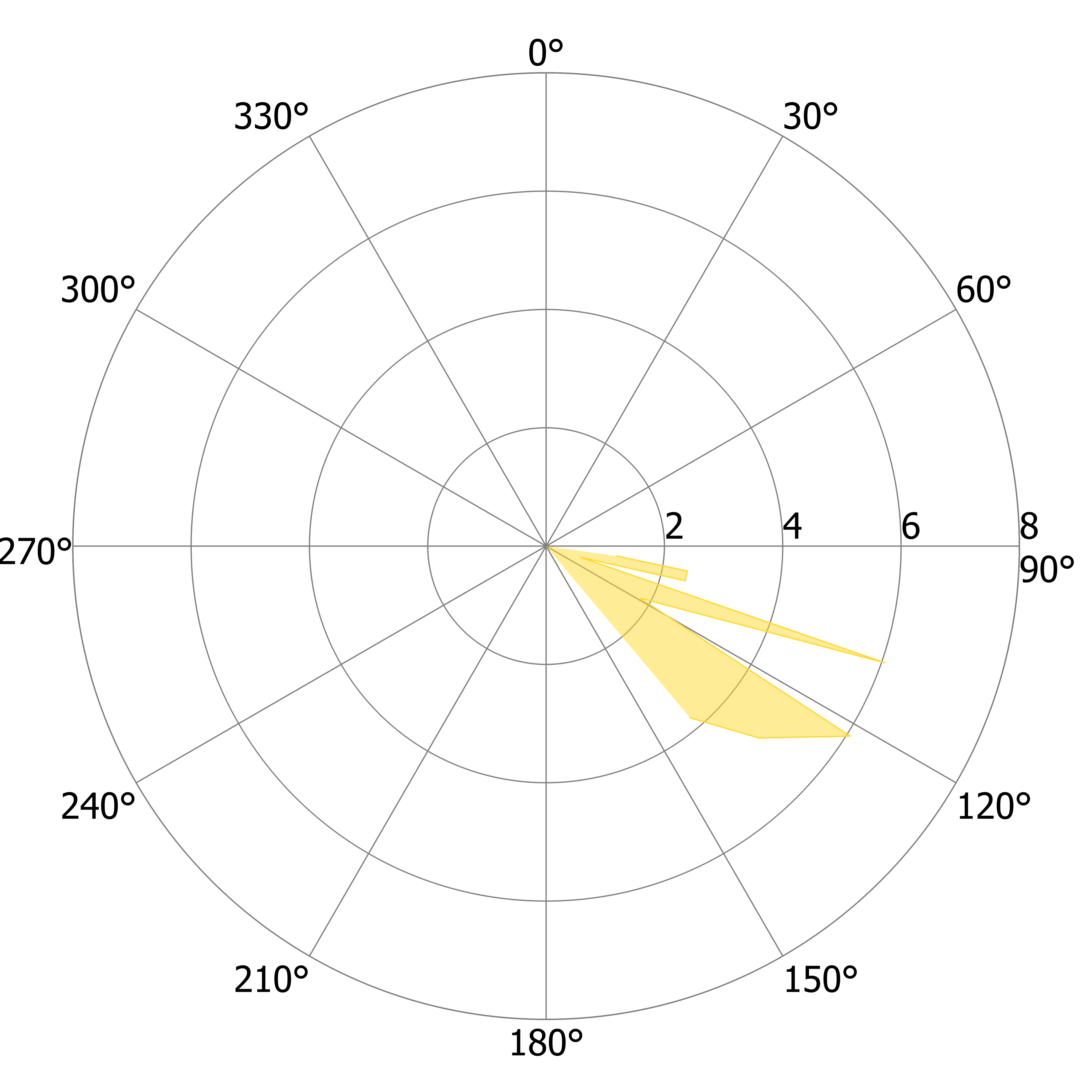

### L_US_p.tiff

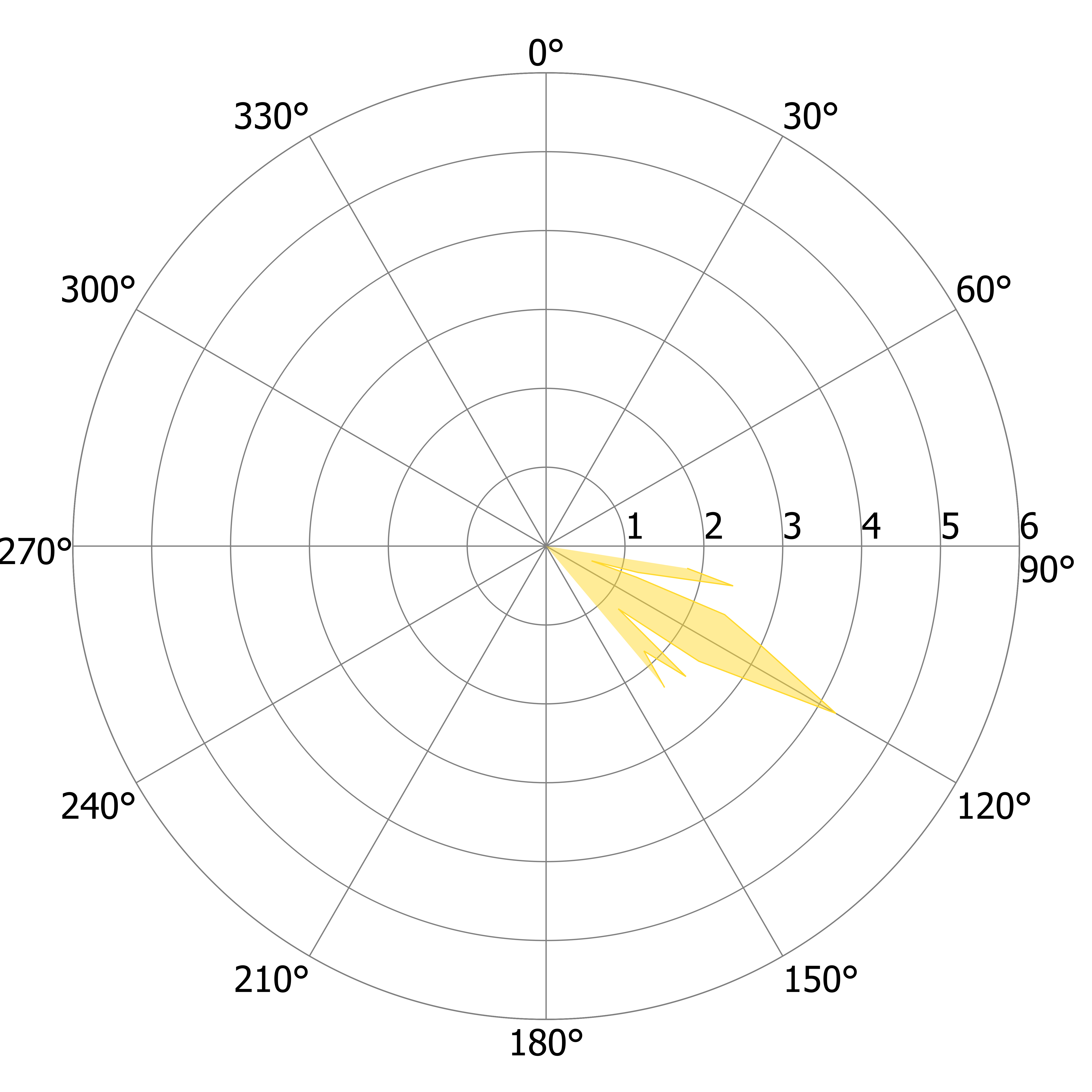

### L_US_q.tiff

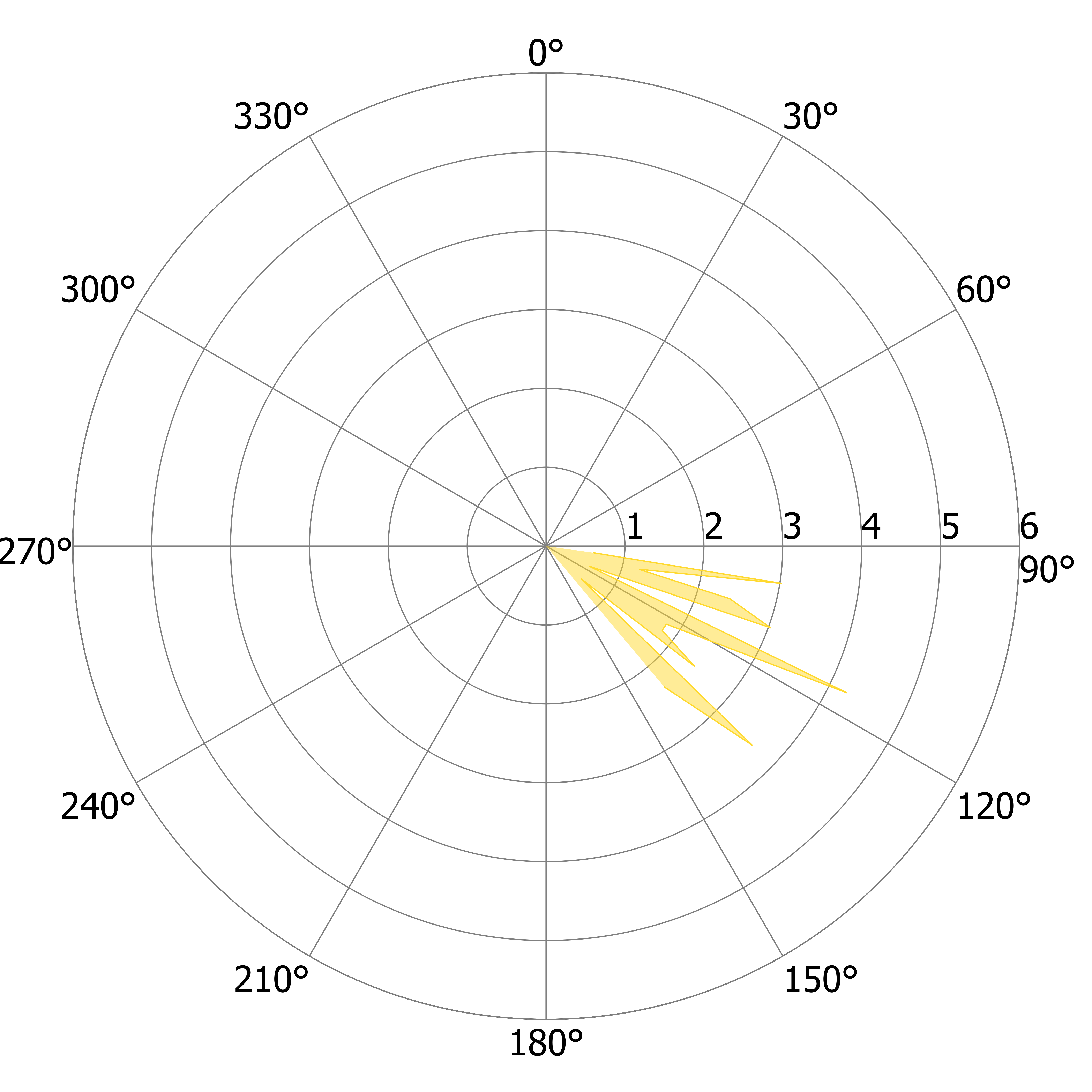

### L_US_r.tiff

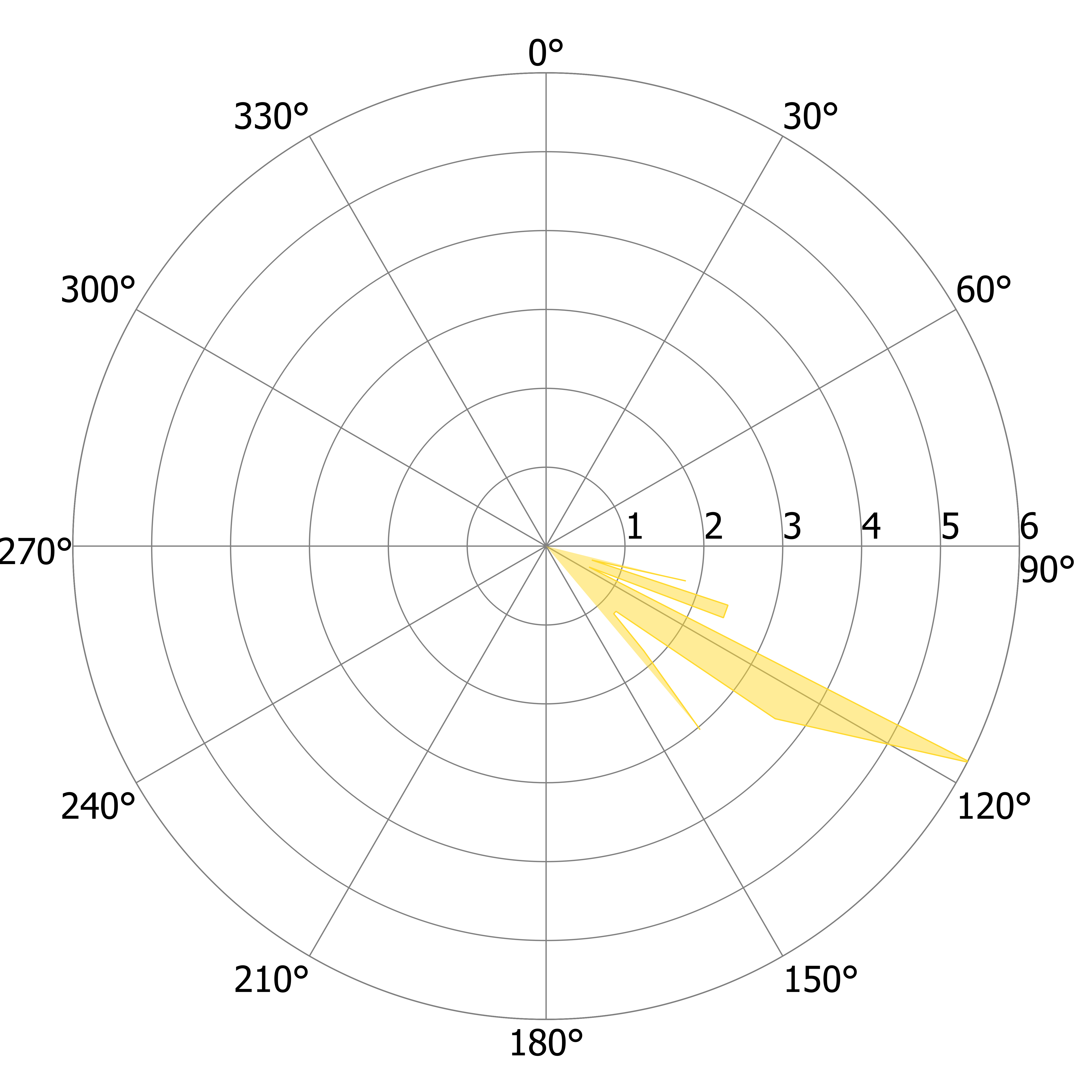

### L_US_s.tiff

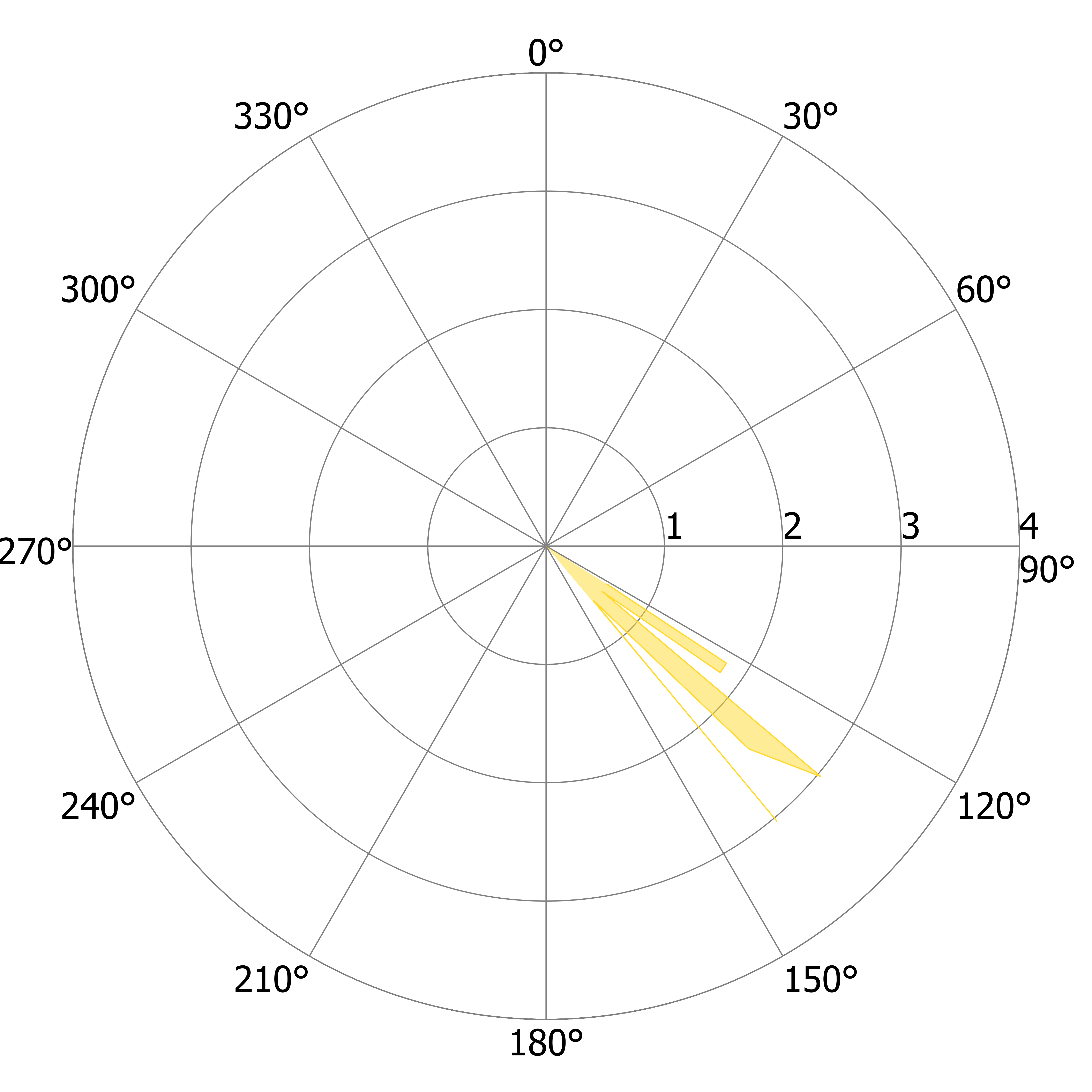

### L_US_t.tiff

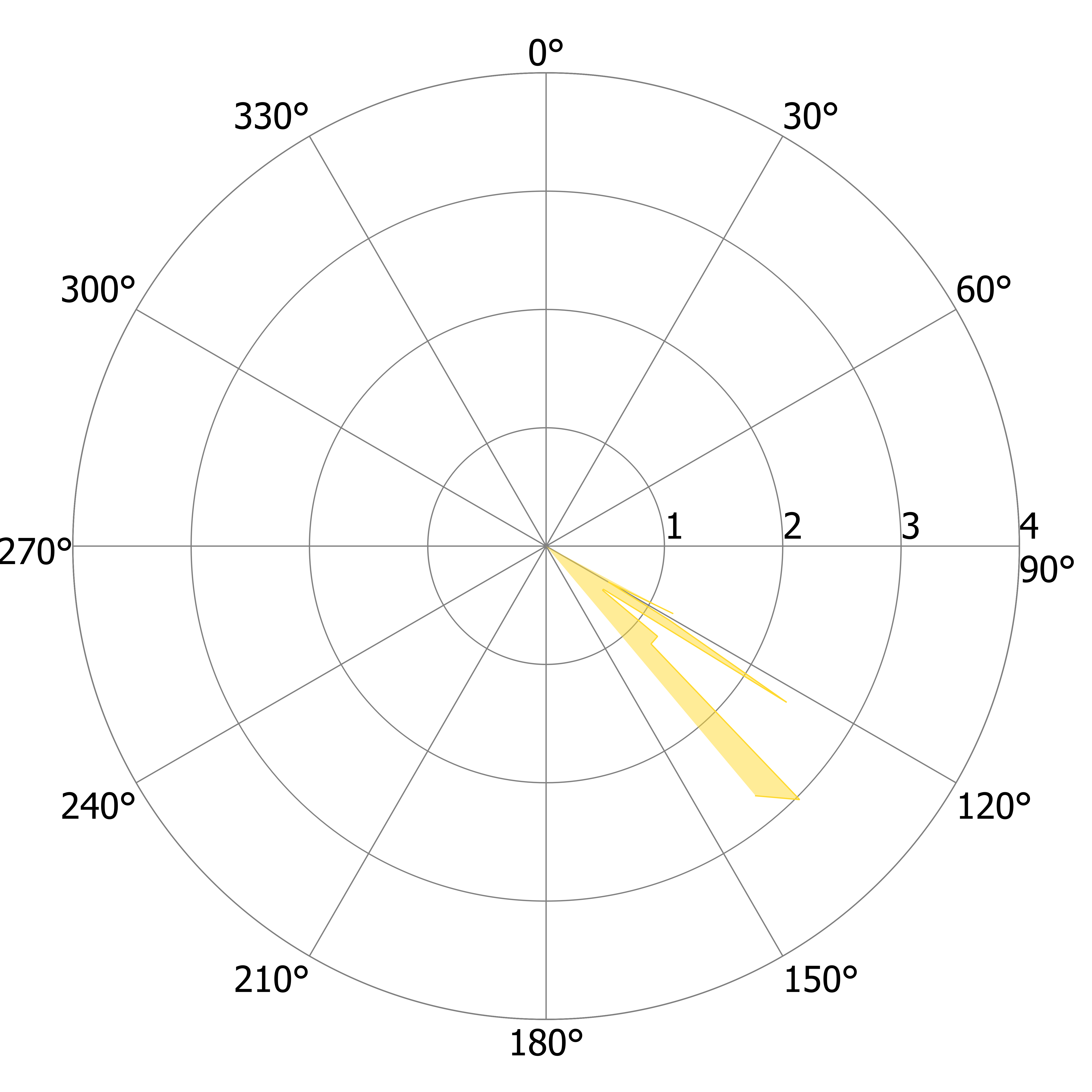

### R_DS_a.tiff

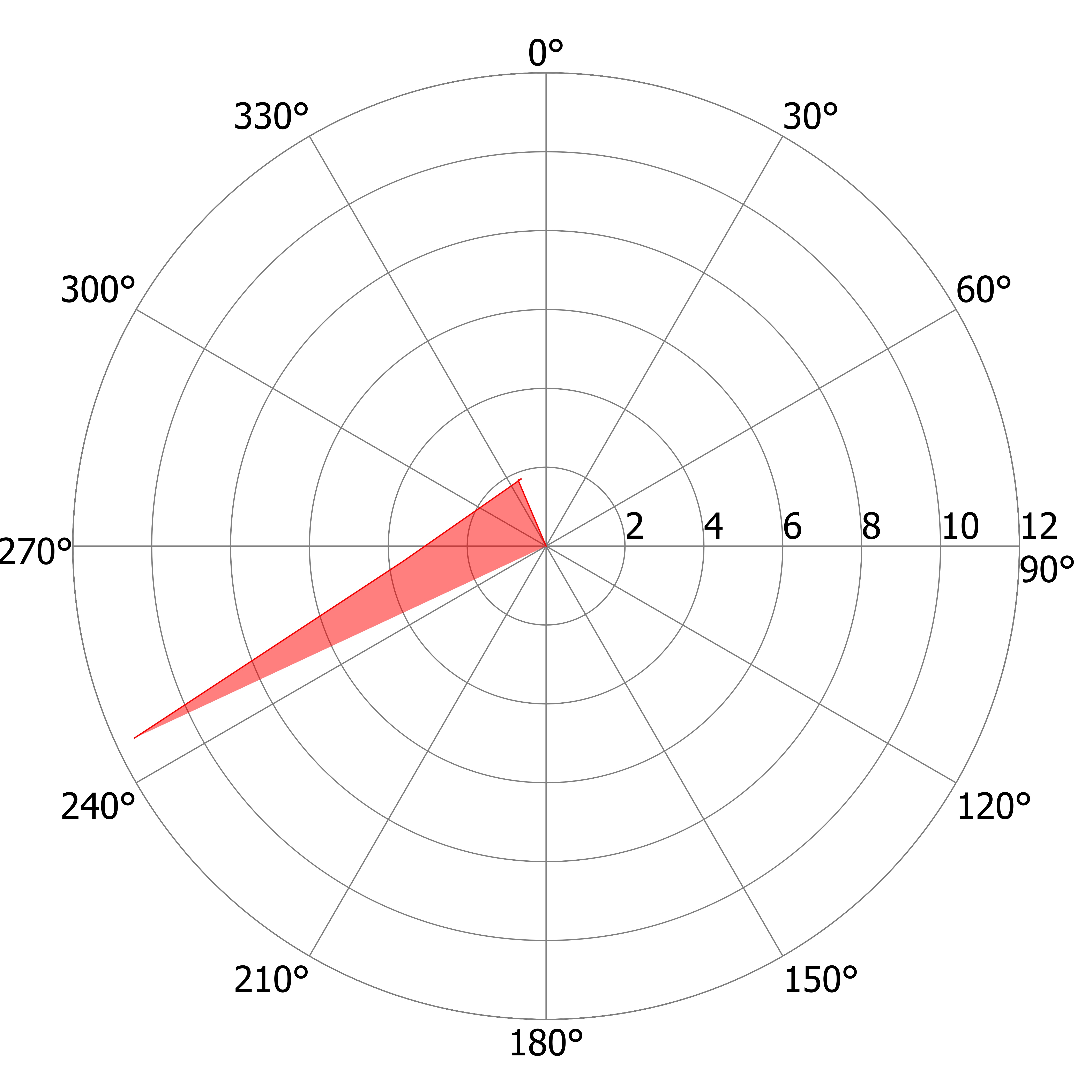

### R_DS_a.tiff

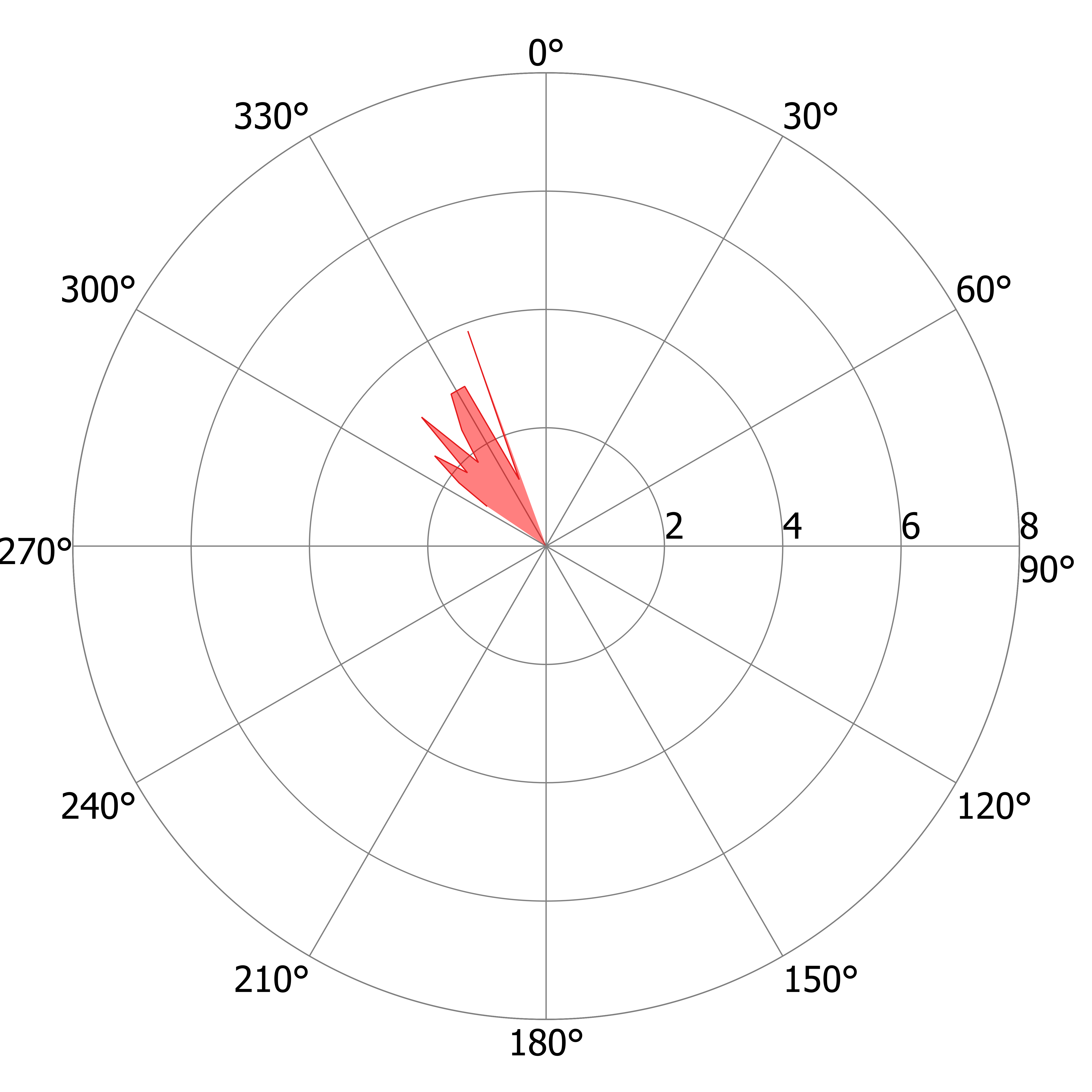

### R_DS_b.tiff

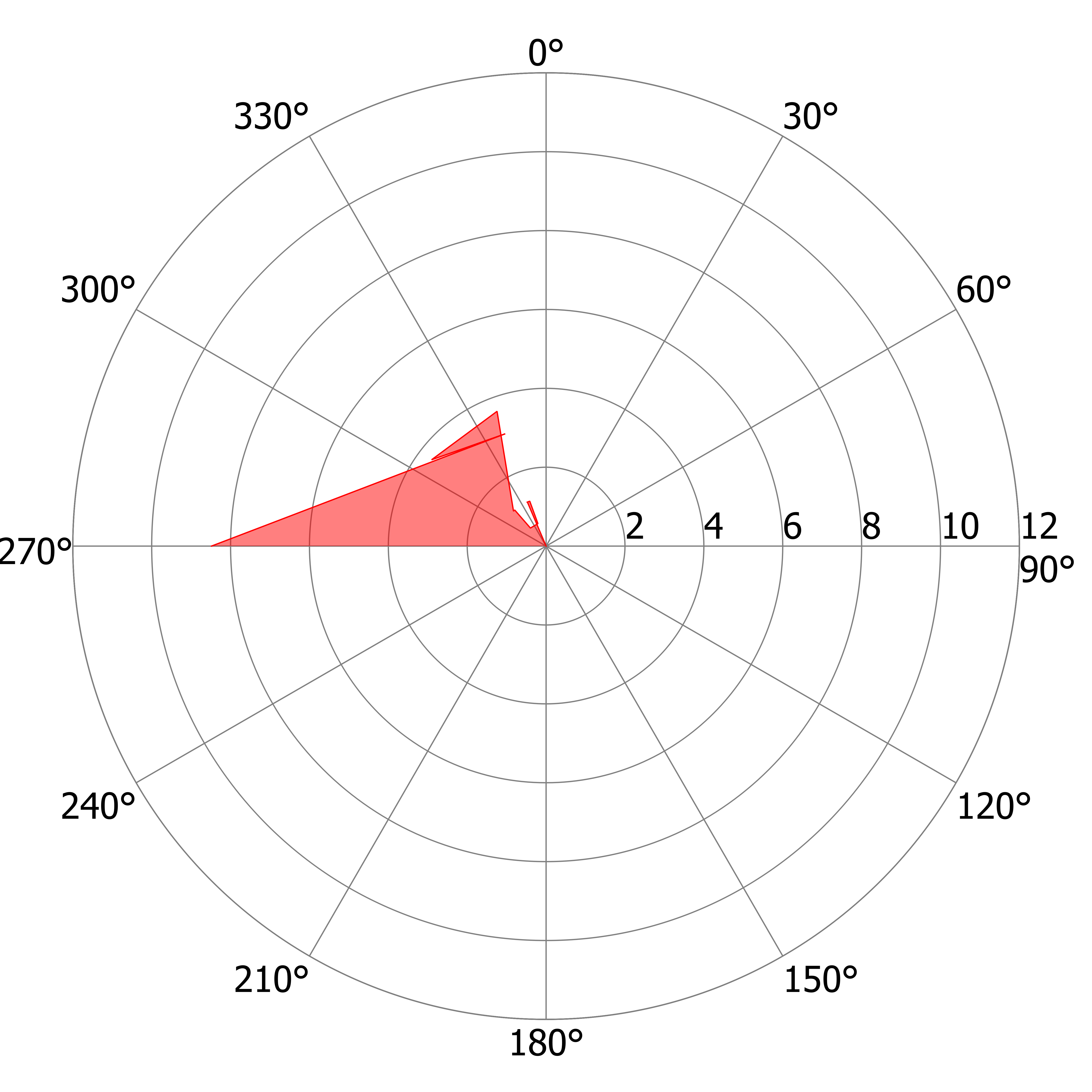

### R_DS_b.tiff

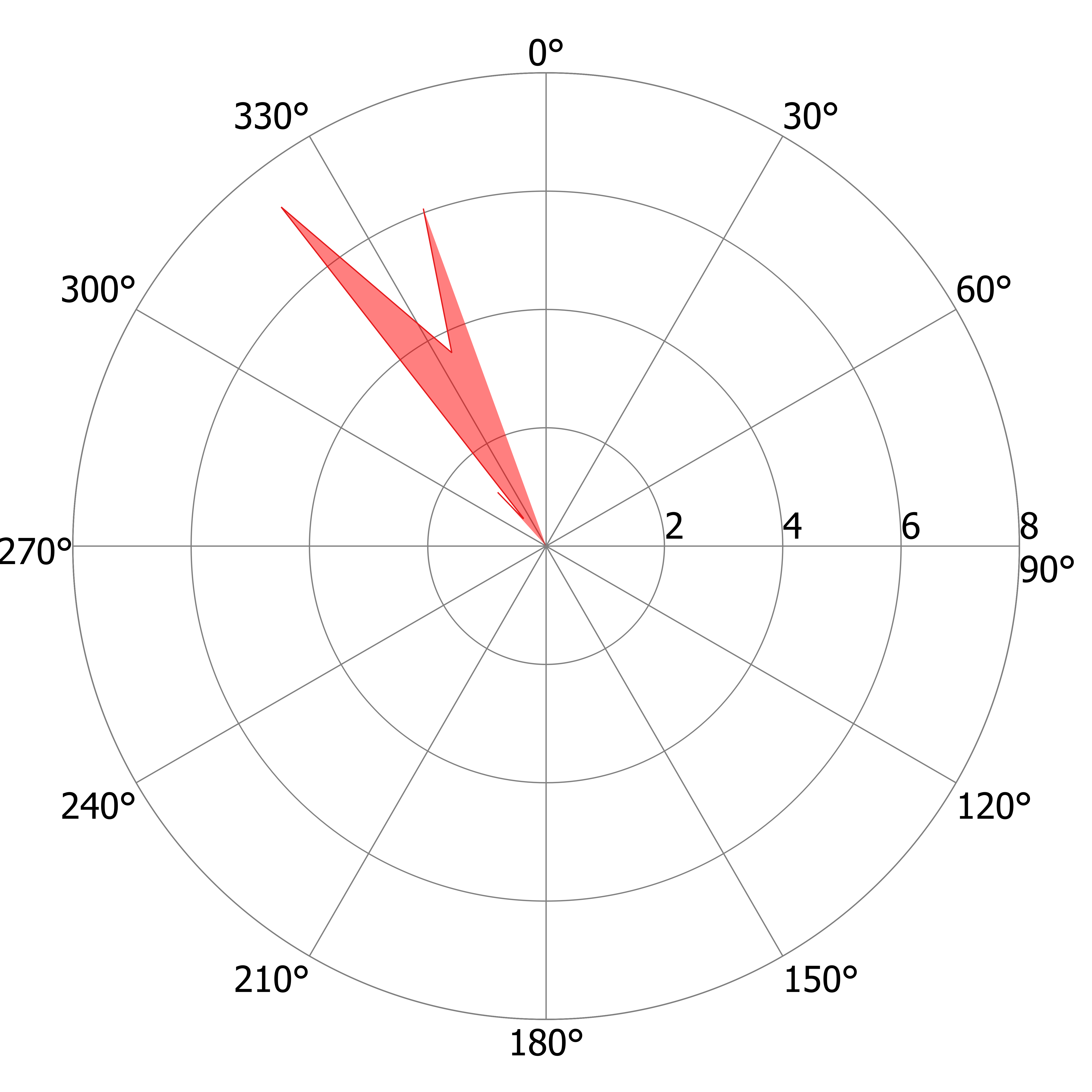

### R_DS_c.tiff

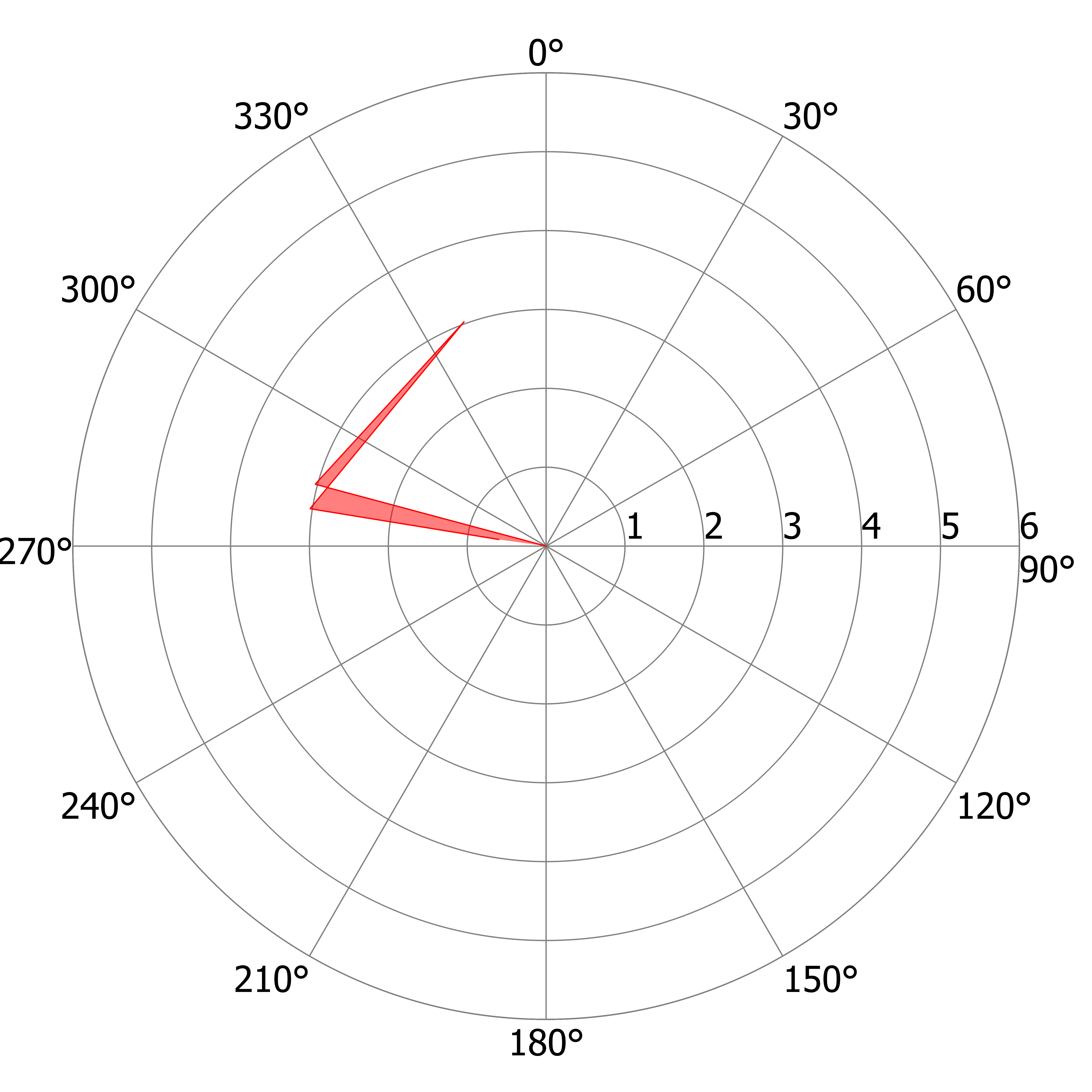

### R_DS_c.tiff

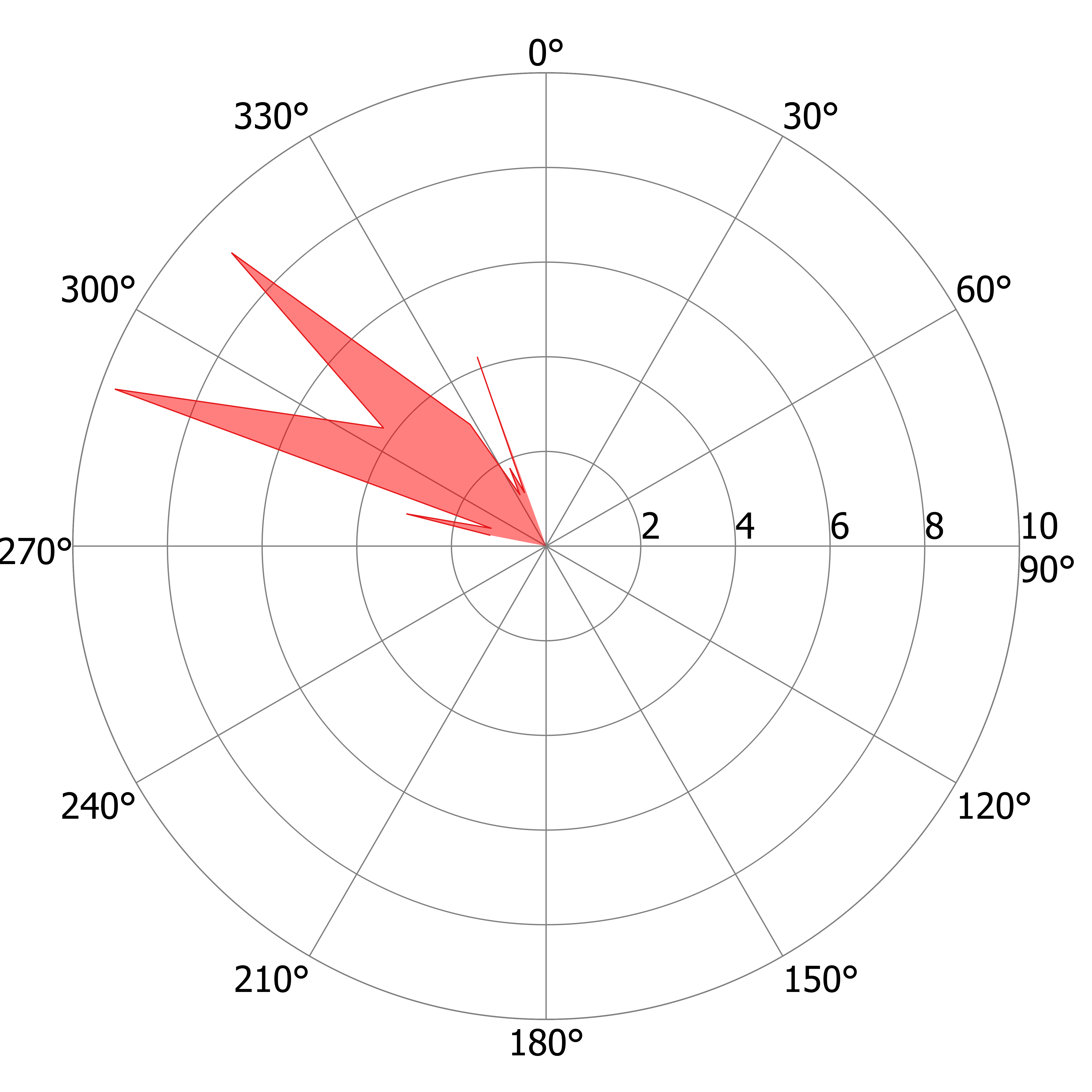

### R_DS_d.tiff

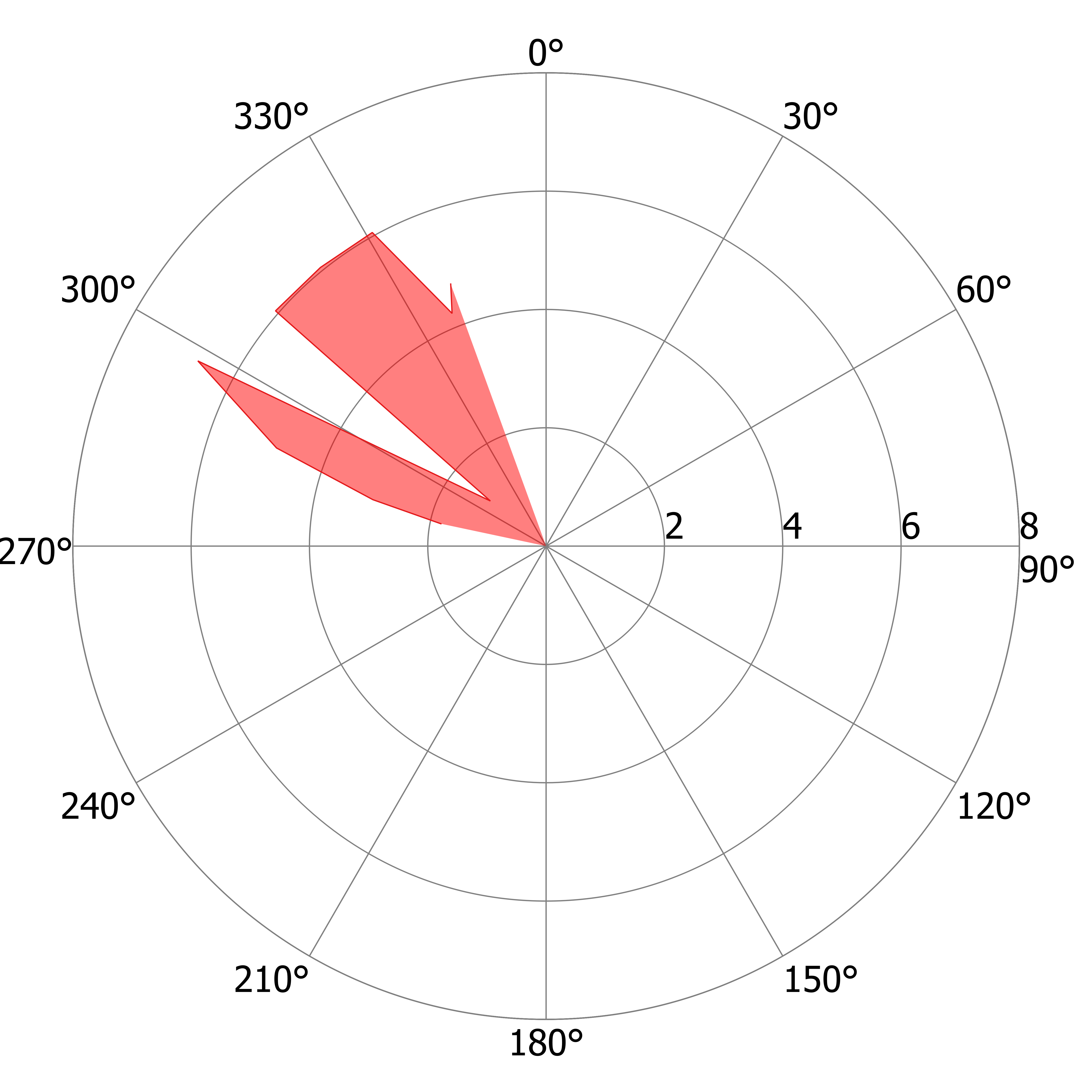

### R_DS_e.tiff

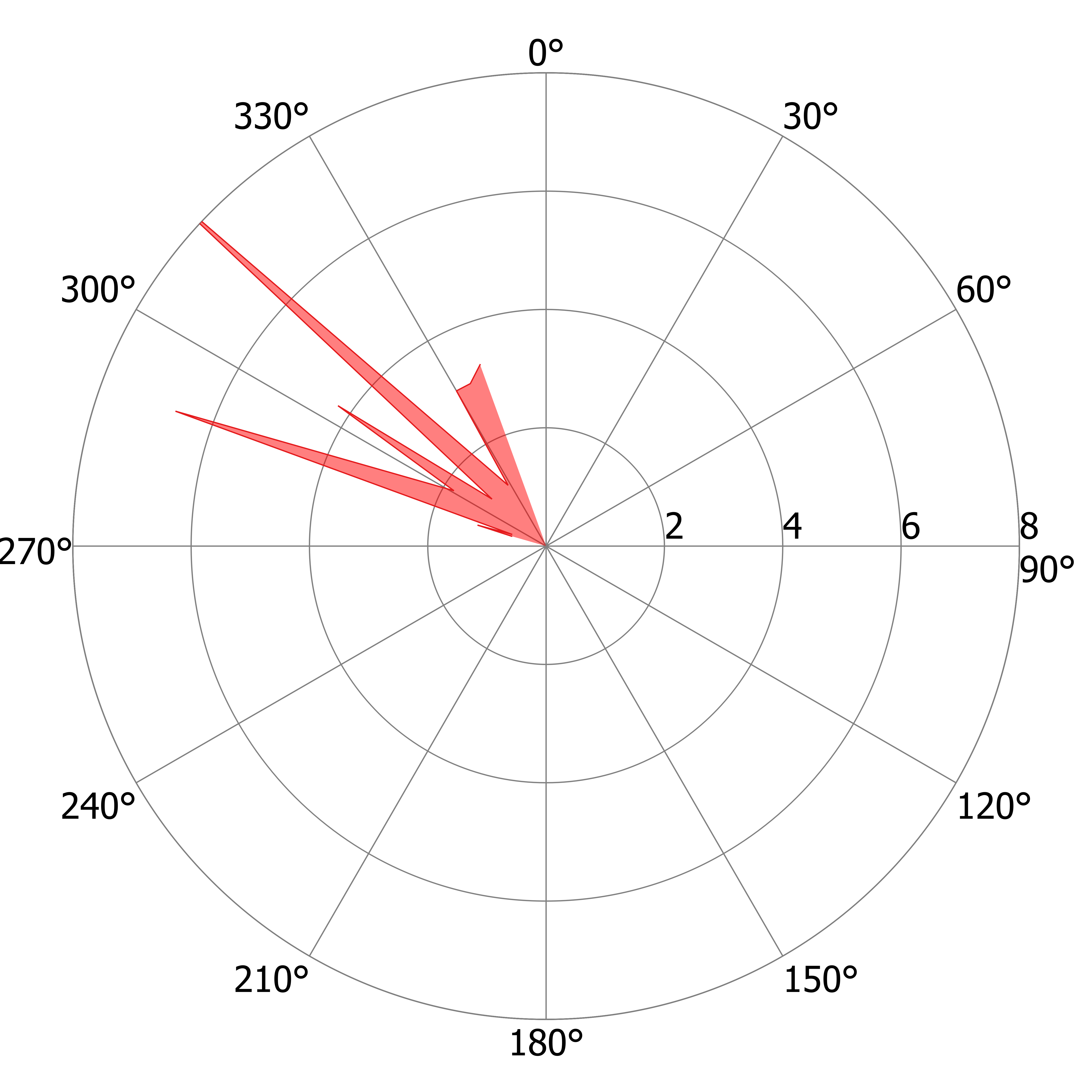

### R_DS_f.tiff

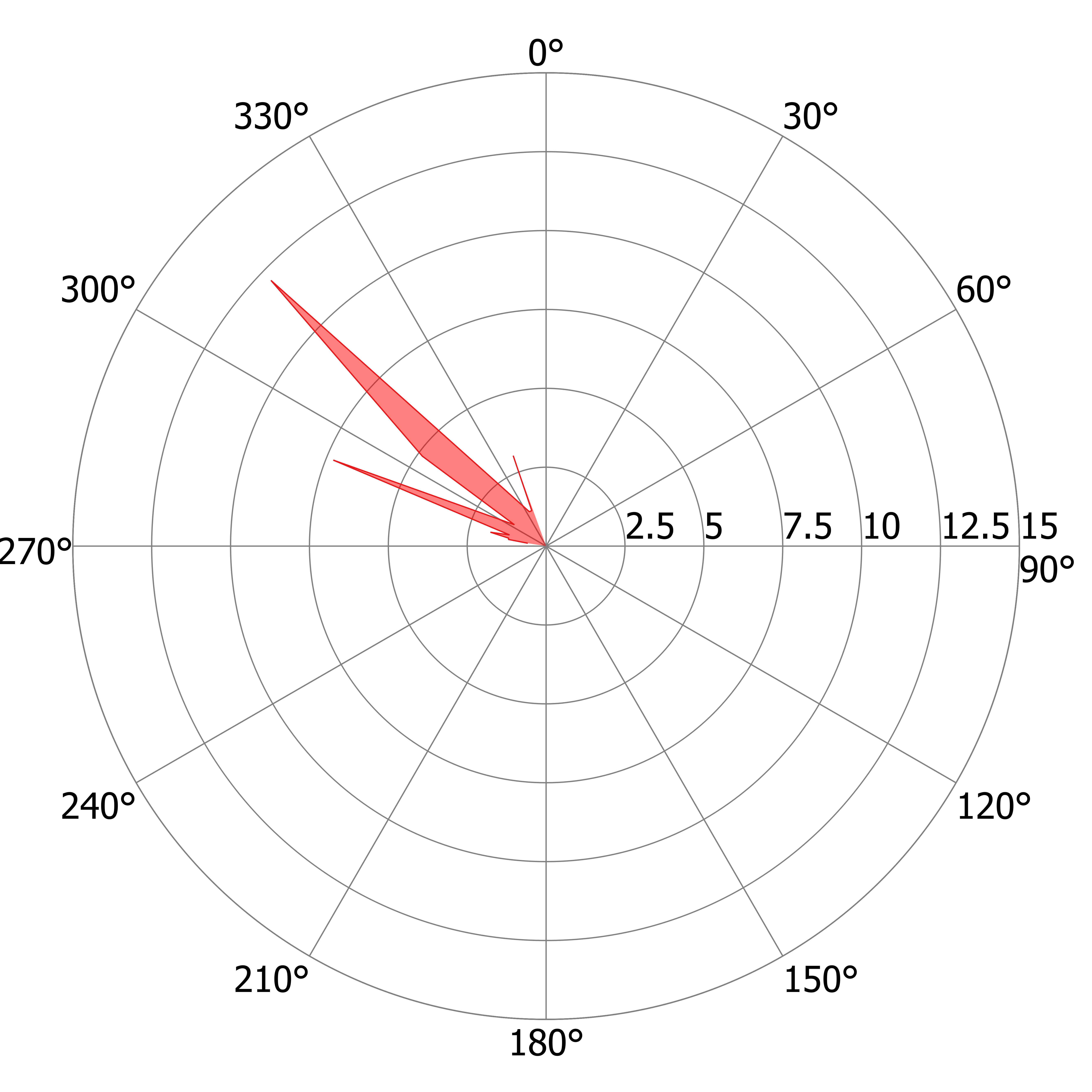

### R_DS_g.tiff

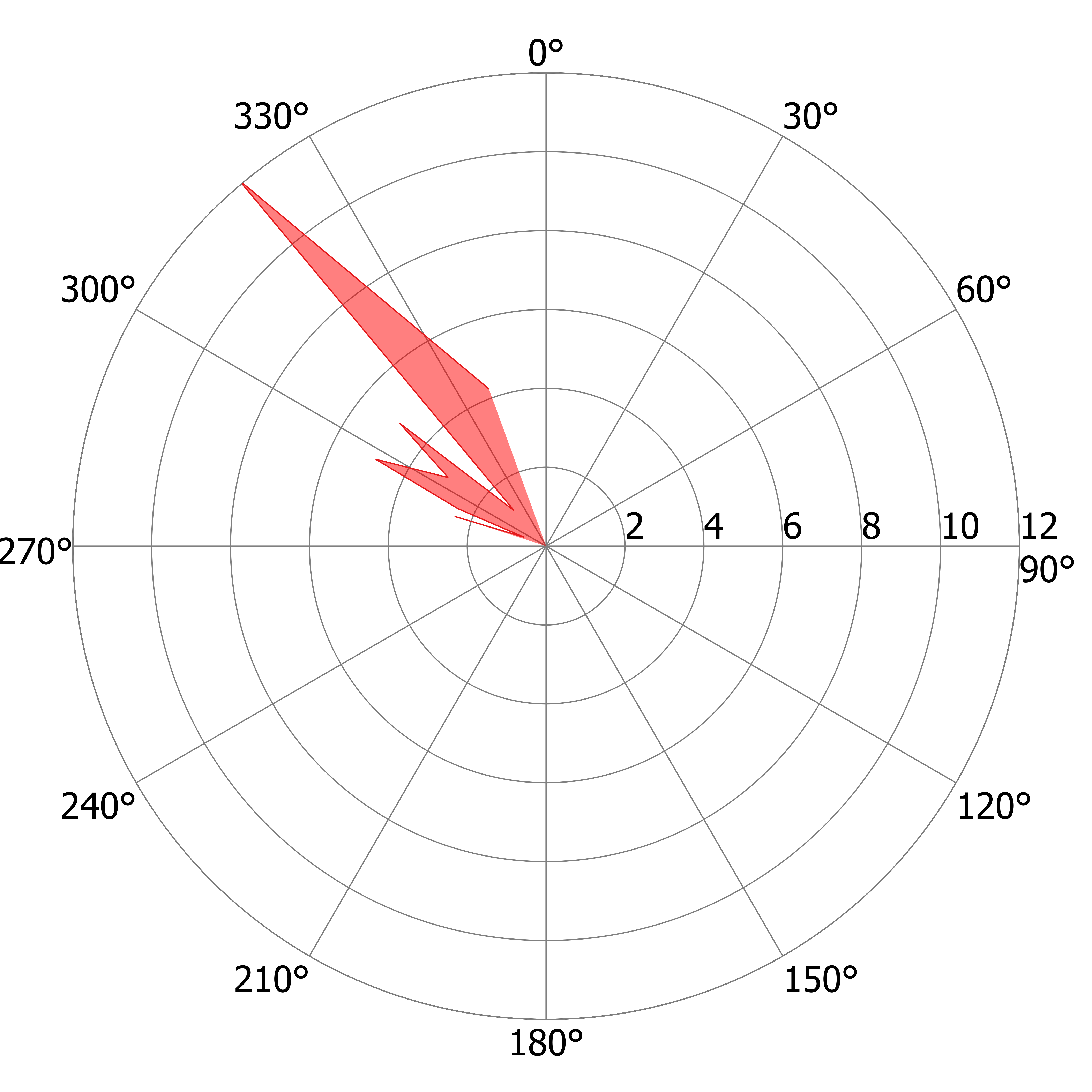

### R_DS_h.tiff

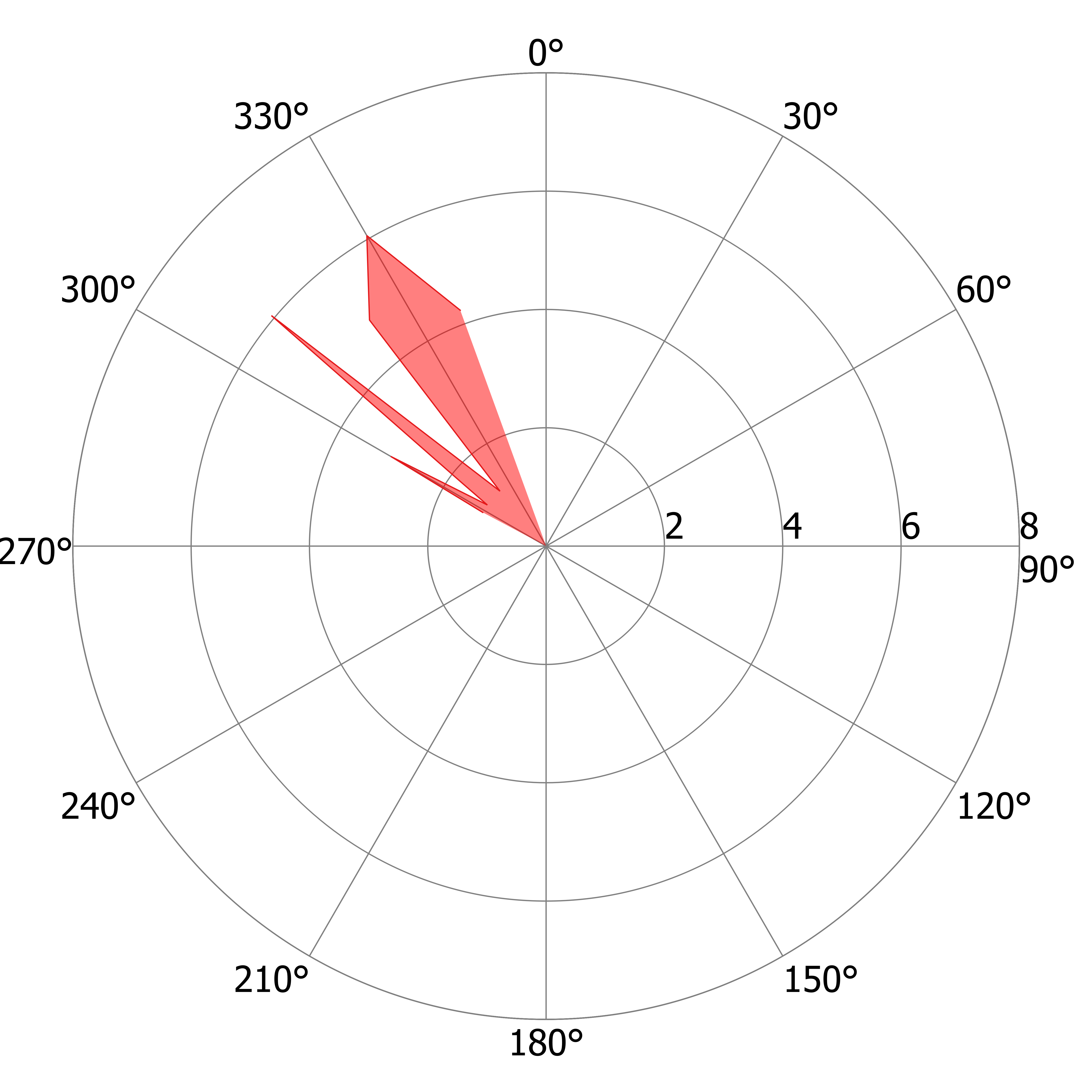

### R_DS_i.tiff

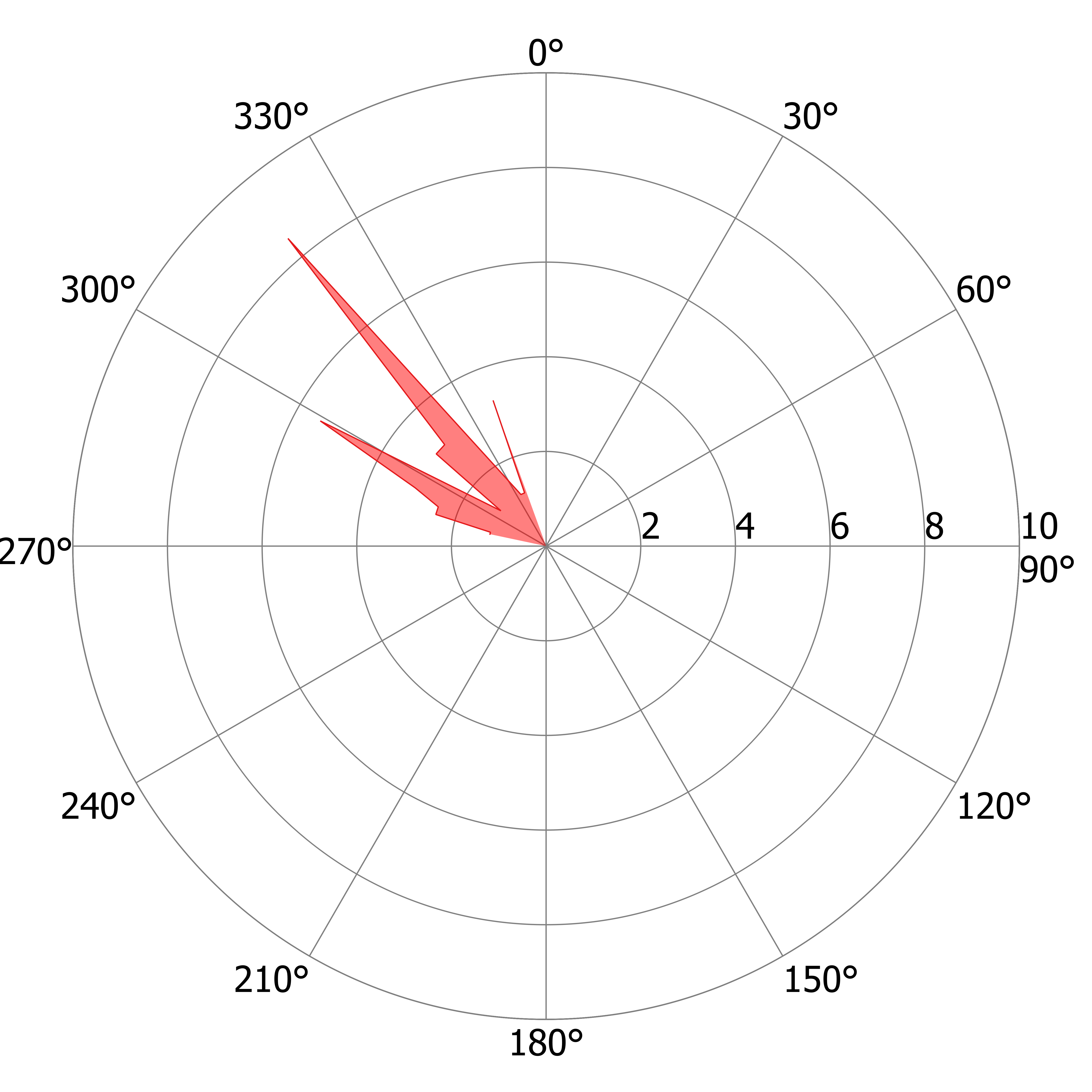

### R_DS_j.tiff

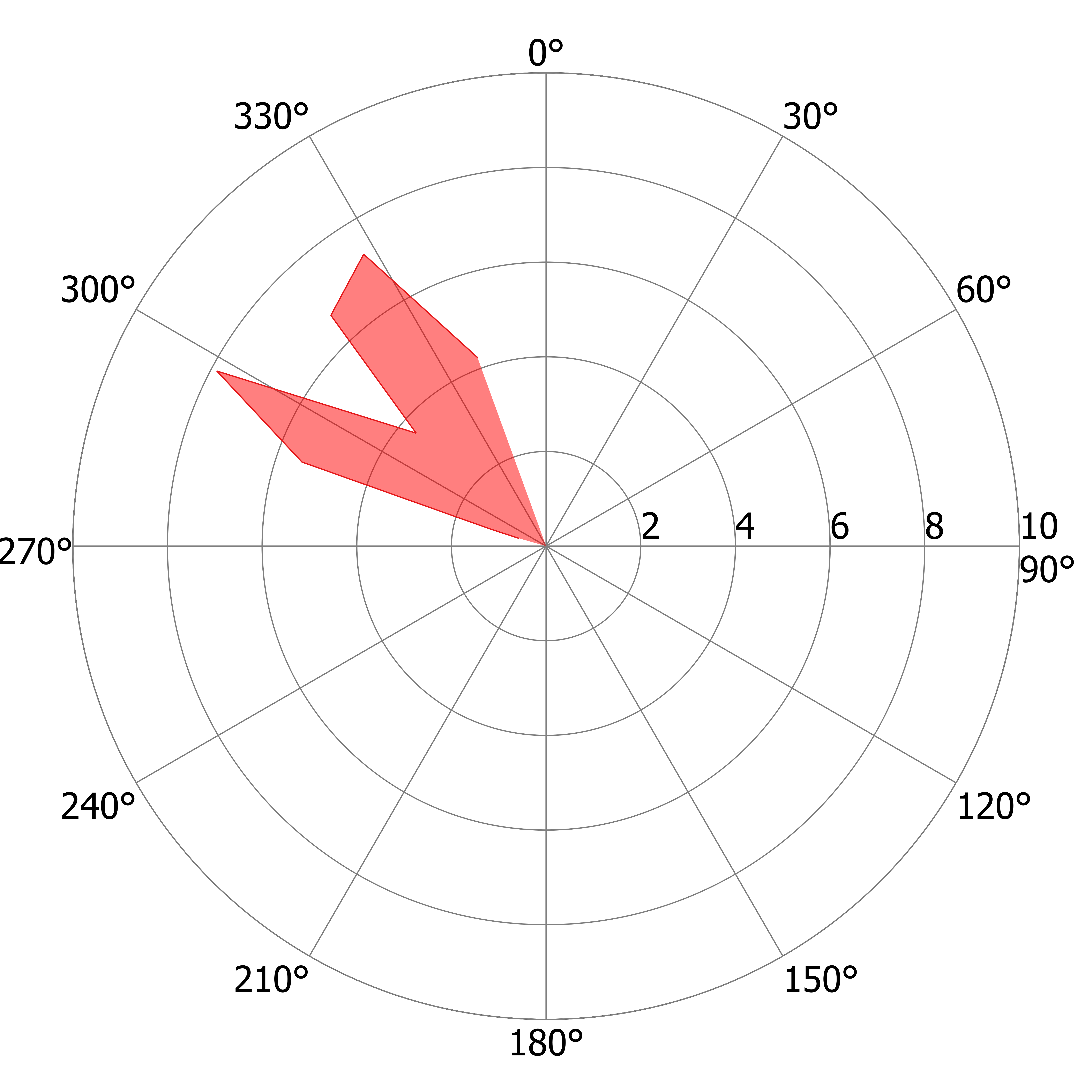

### R_DS_k.tiff

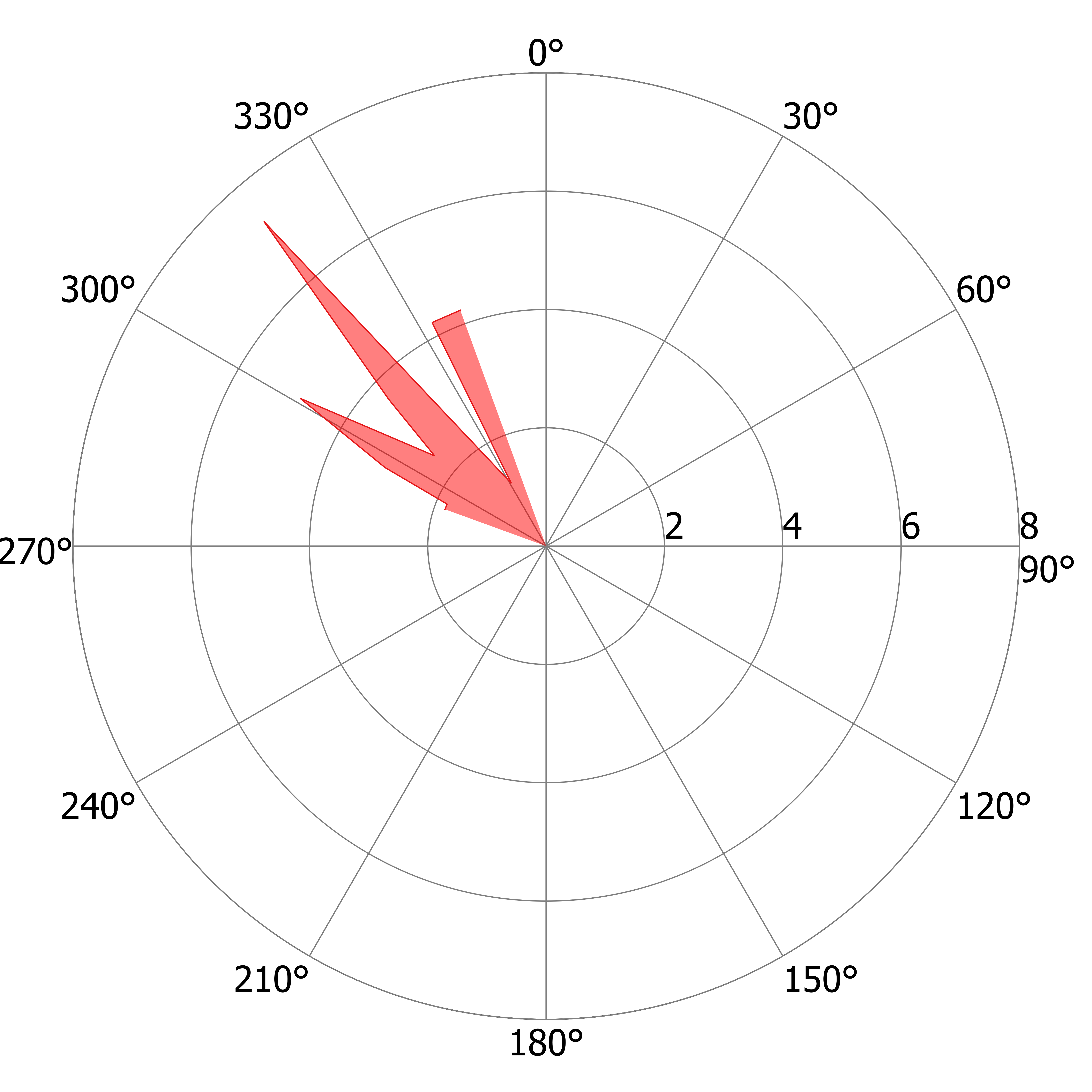

### R_US_a.tiff

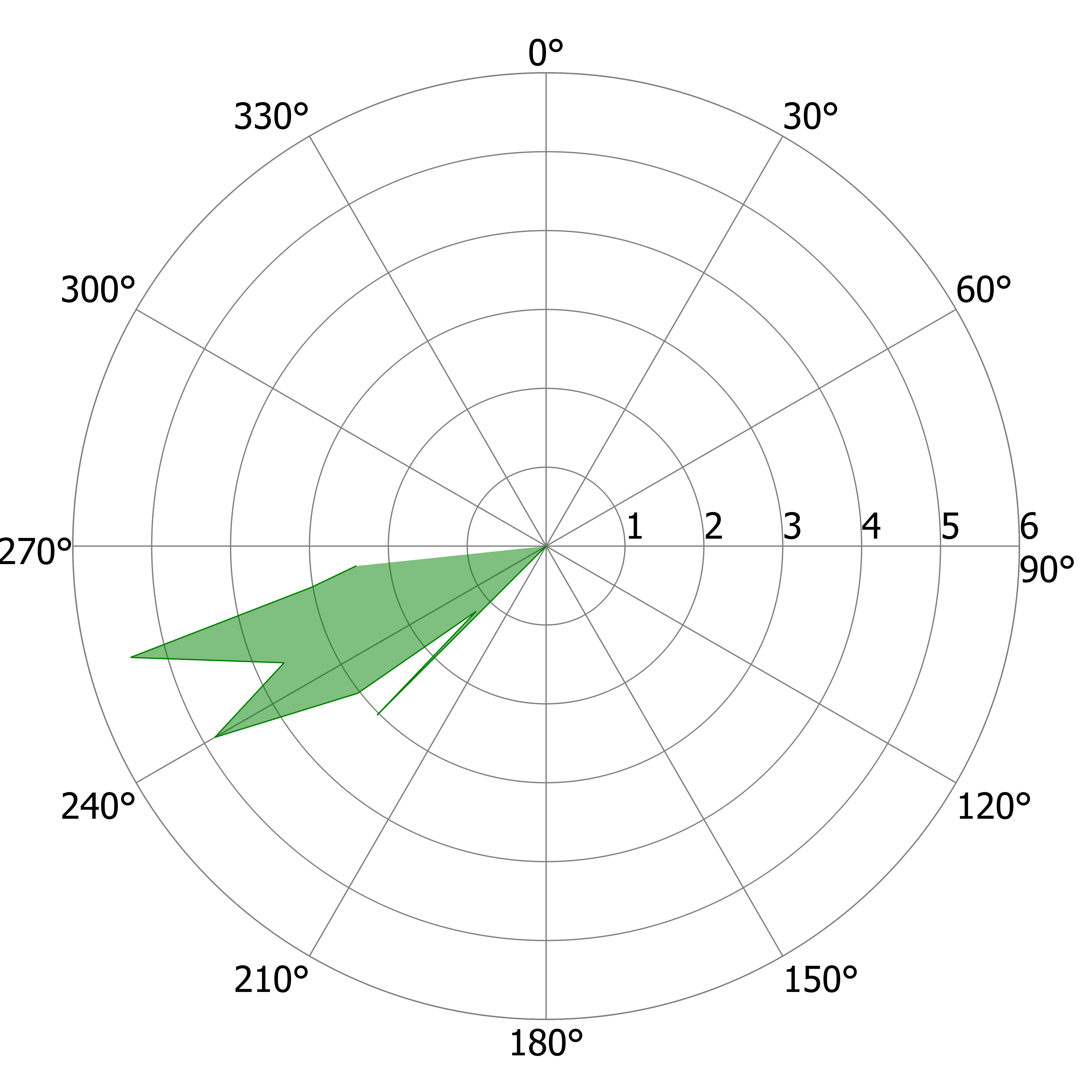

### R_US_a.tiff

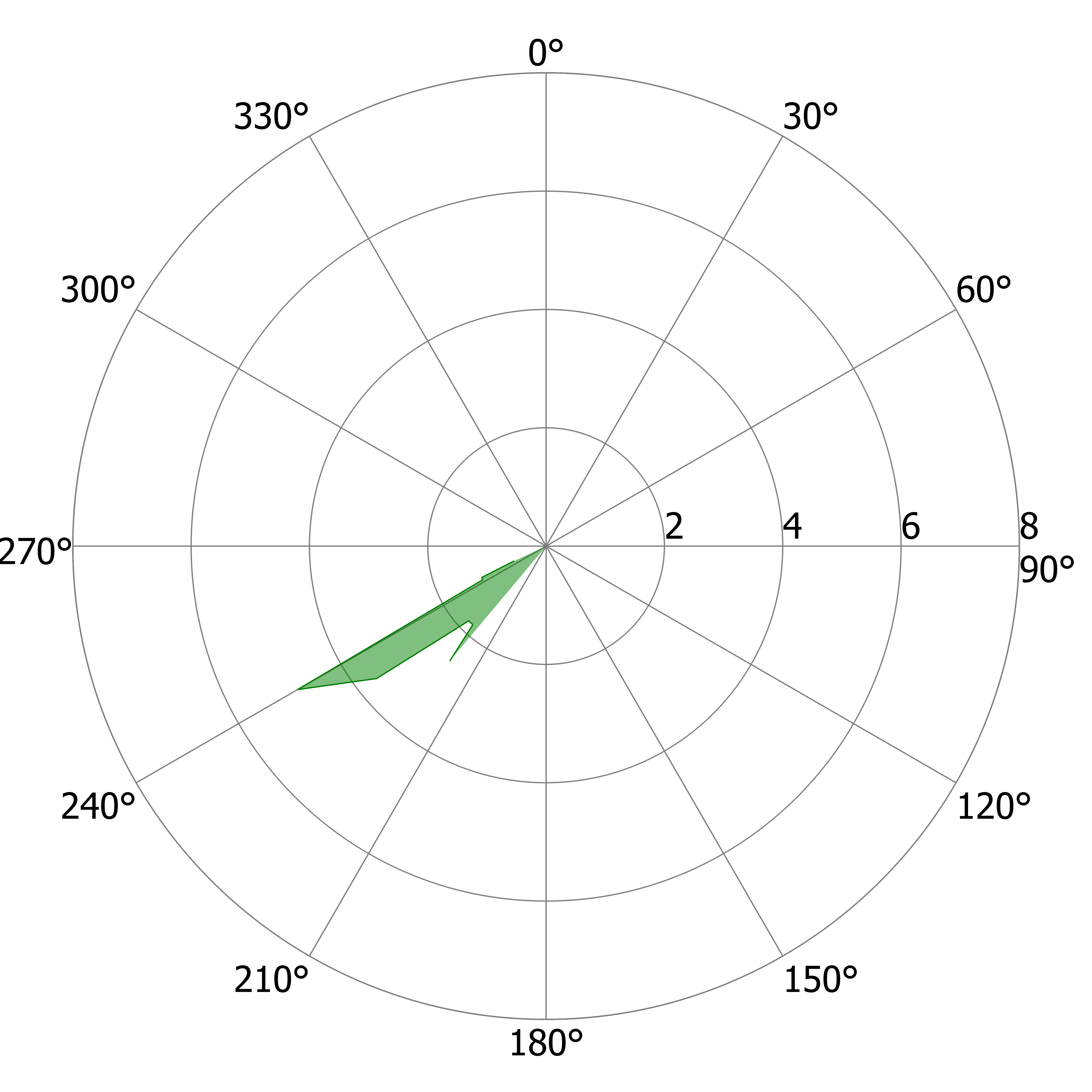

### R_US_b.tiff

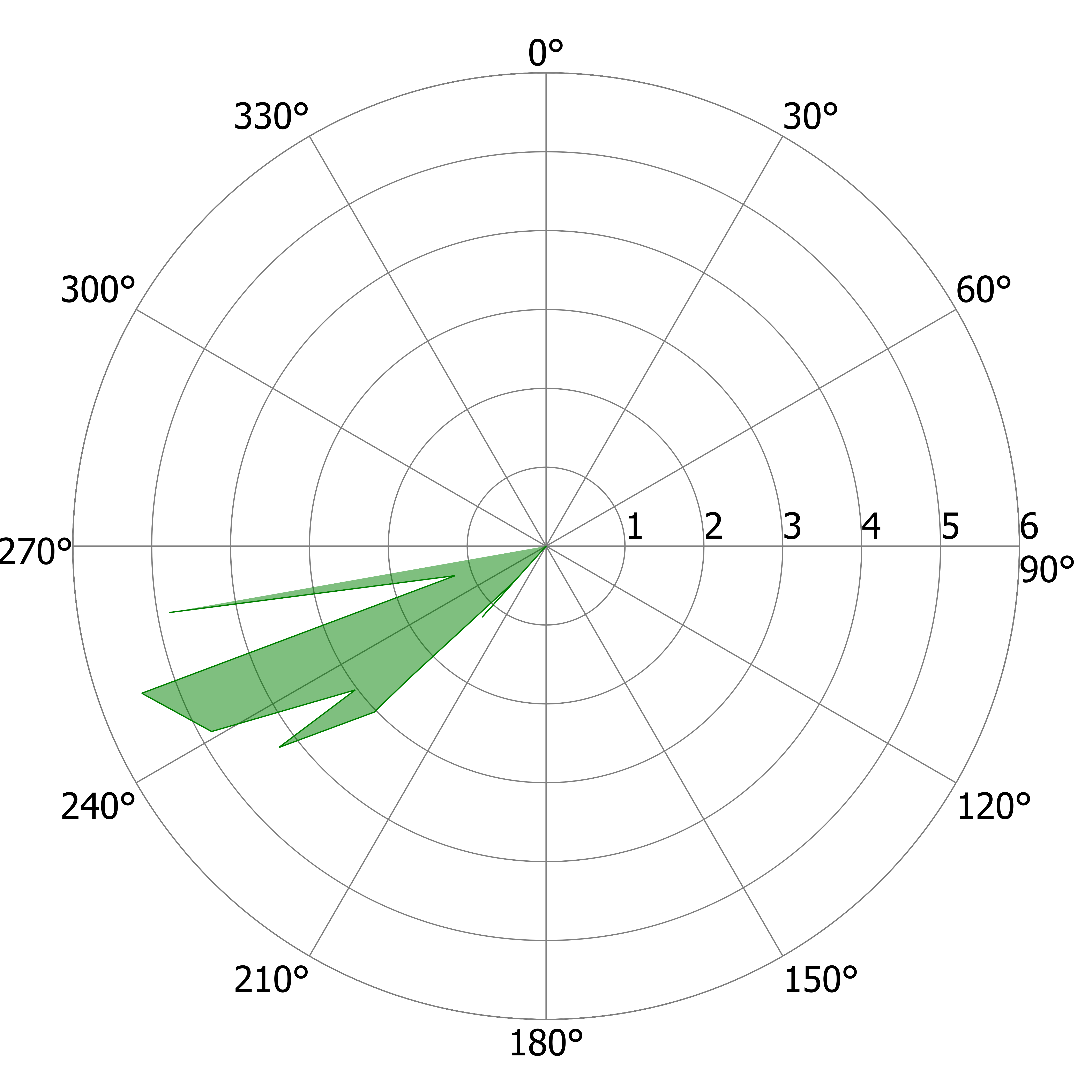

### R_US_b.tiff

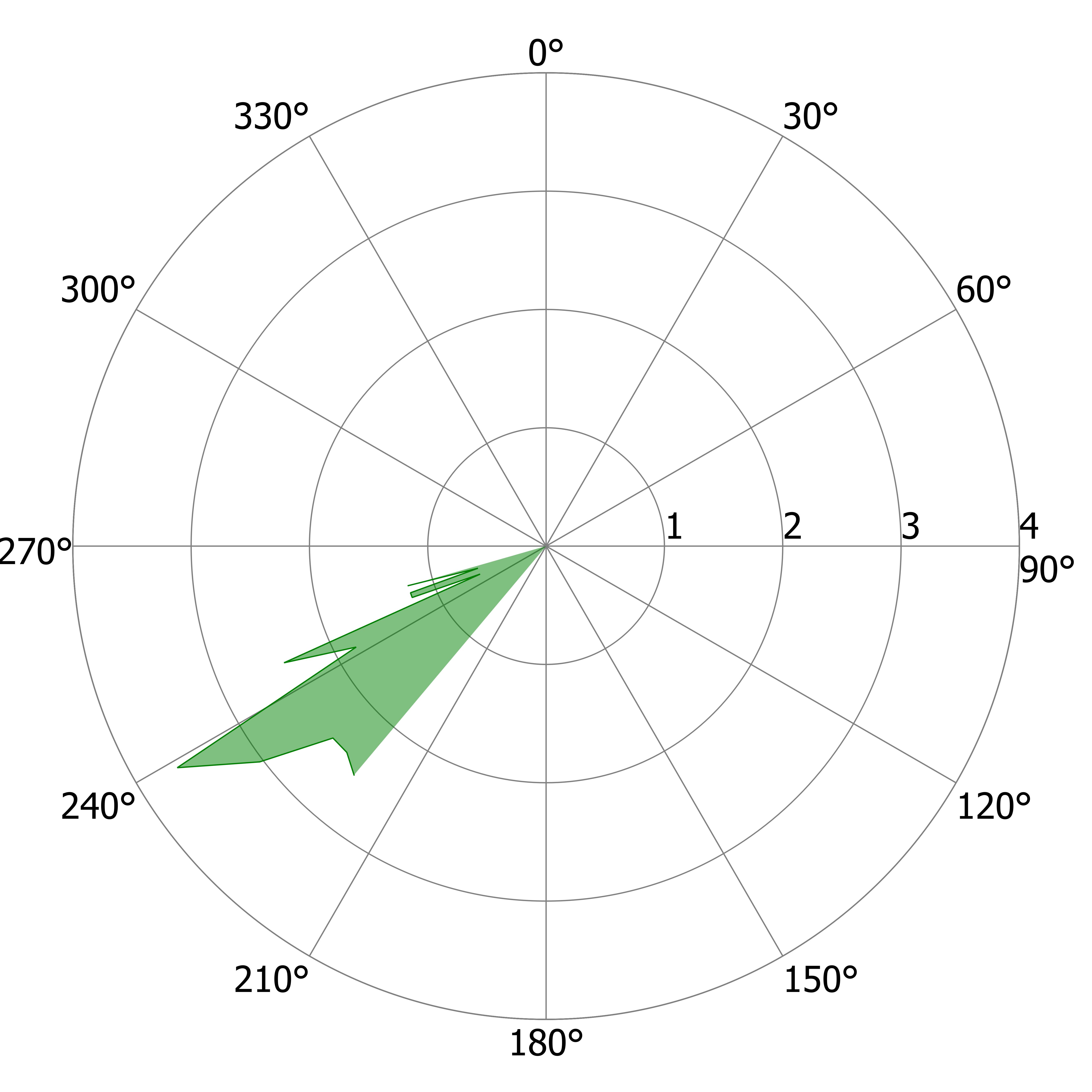

### R_US_c.tiff

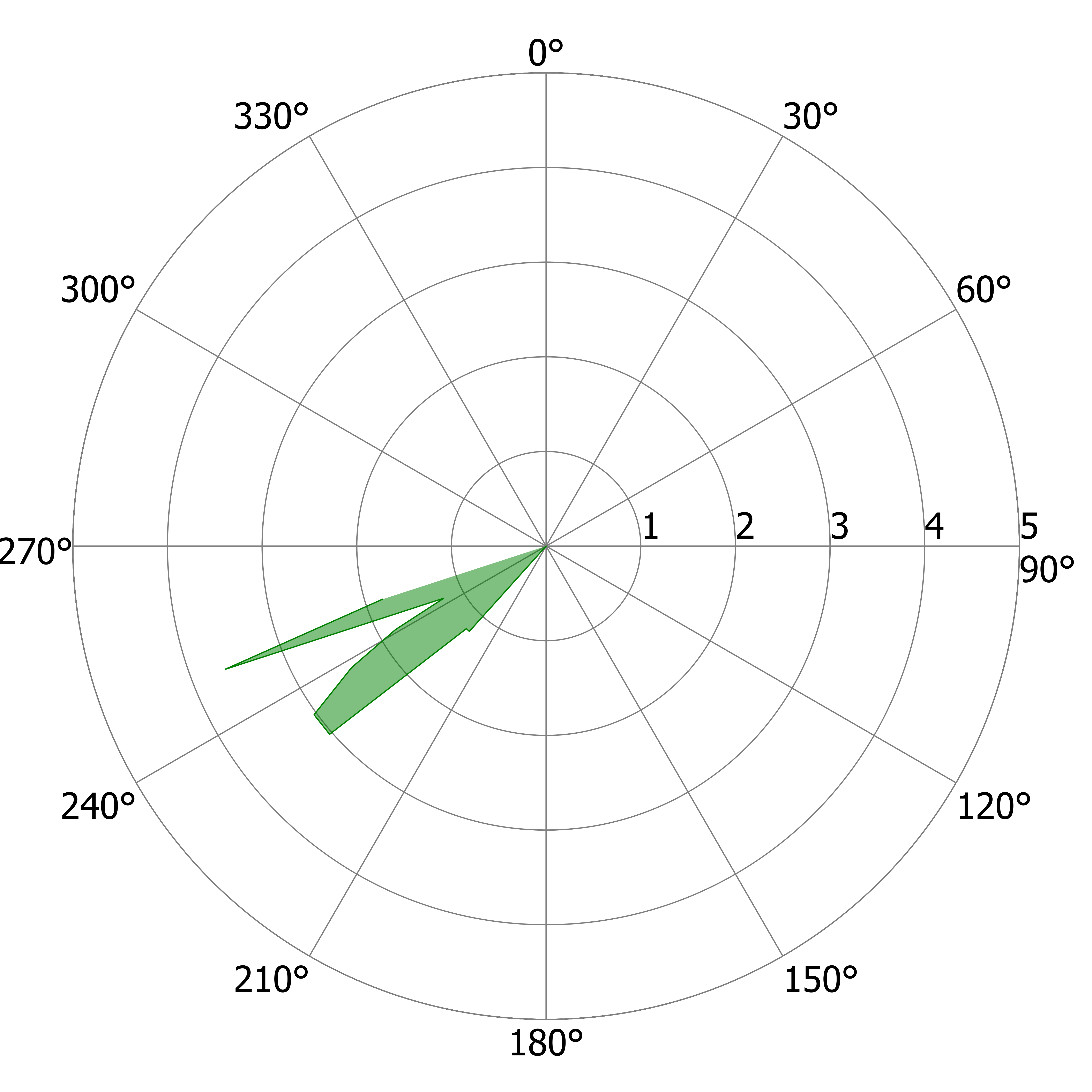

### R_US_c.tiff

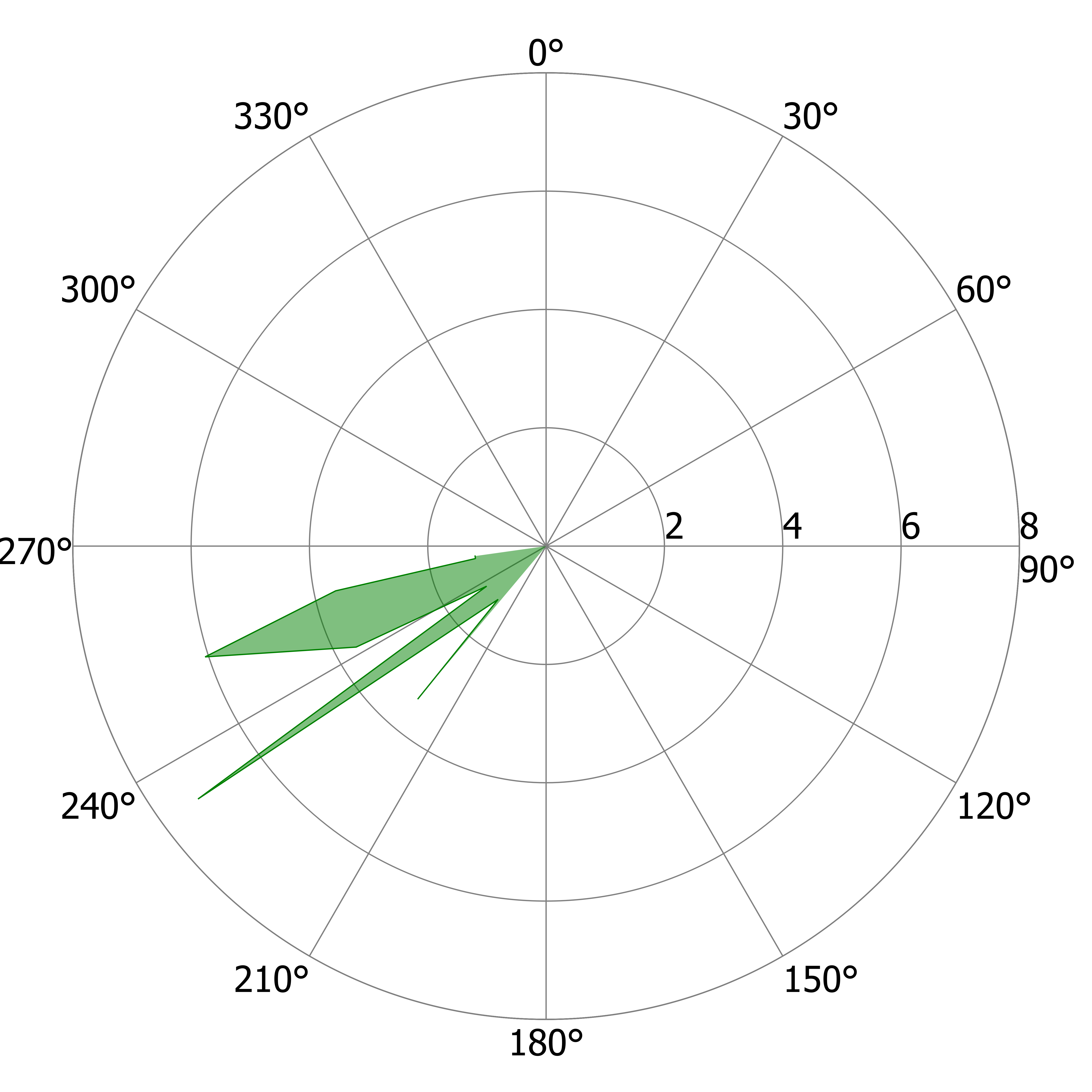

### R_US_d.tiff

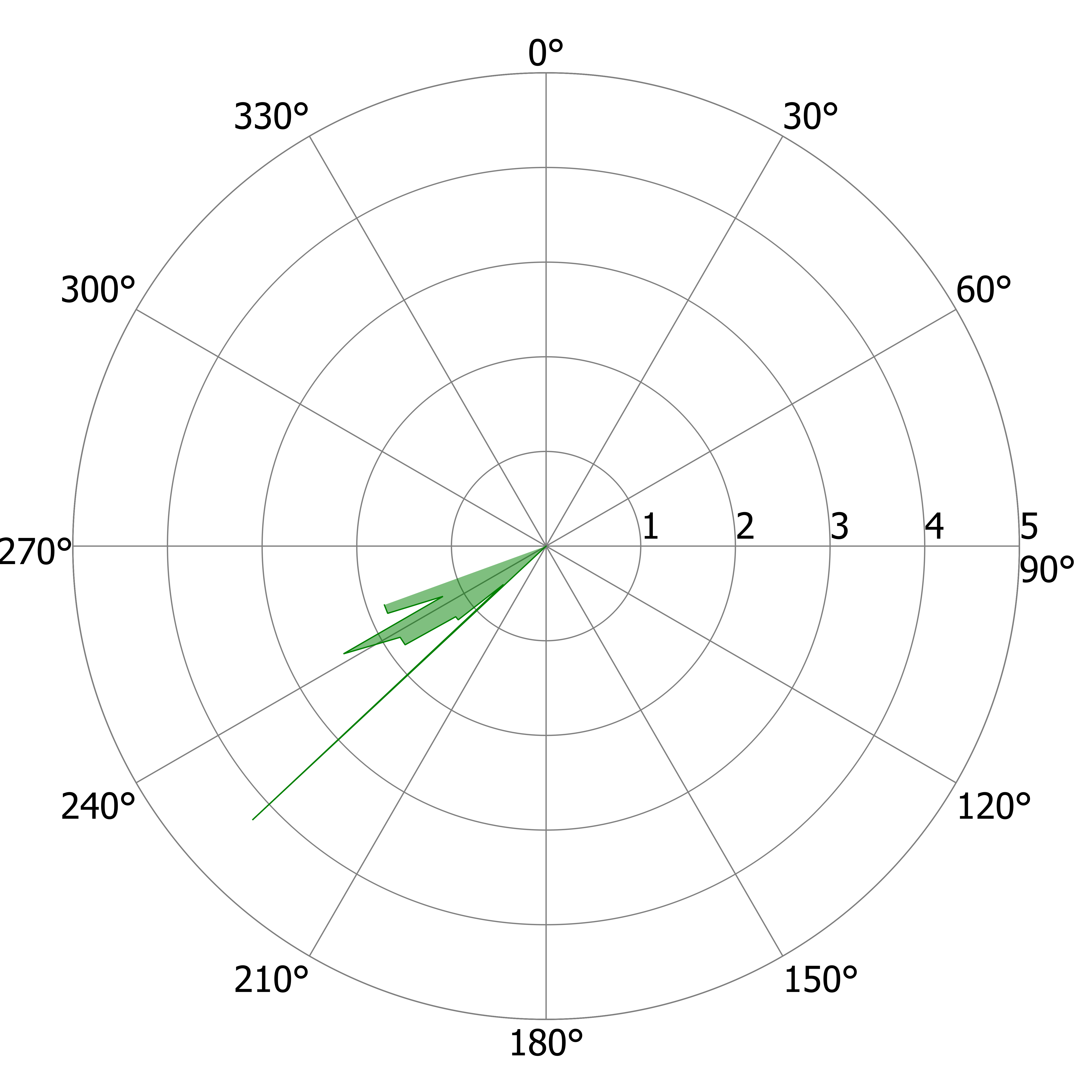

### R_US_d.tiff

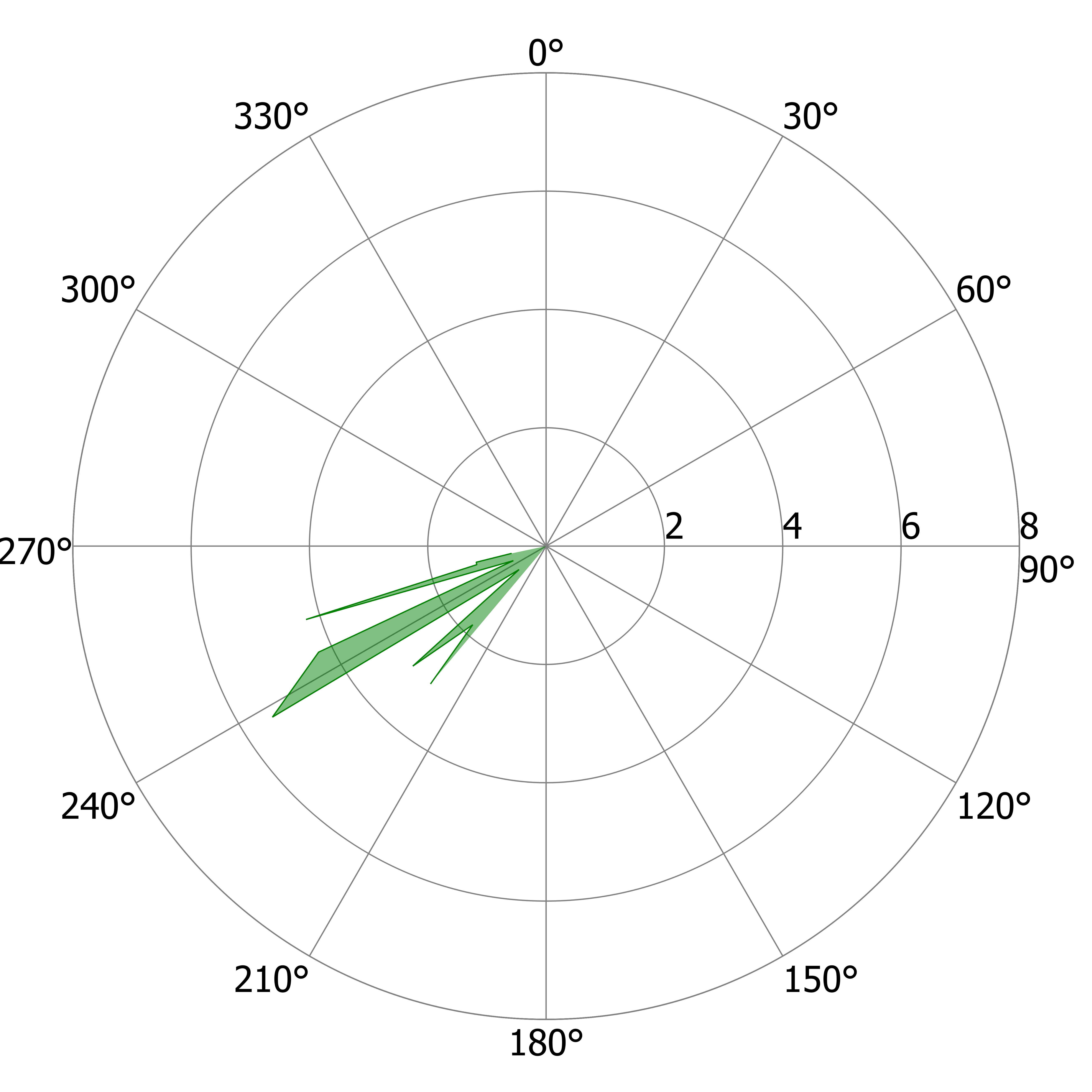

### R_US_e.tiff

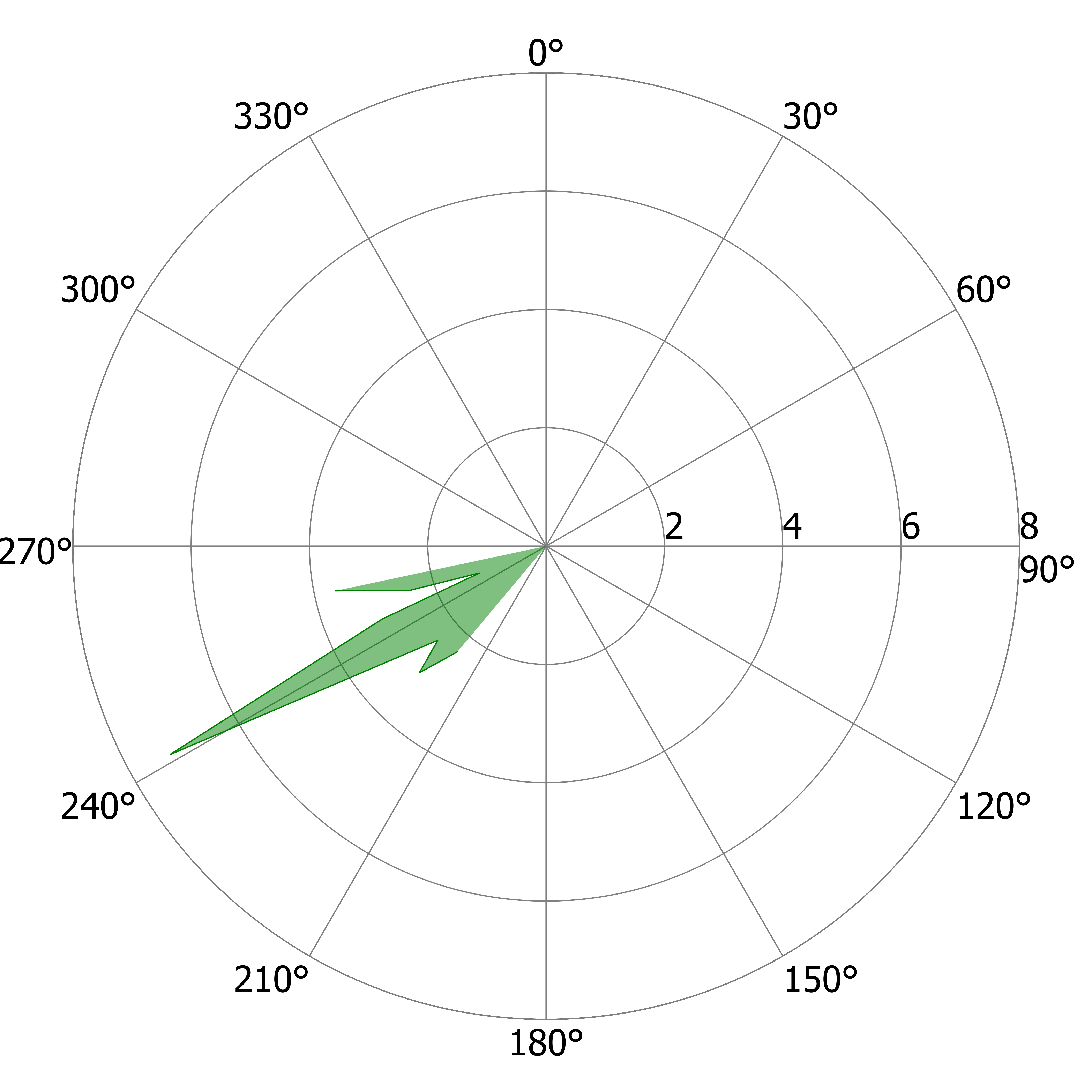

### R_US_f.tiff

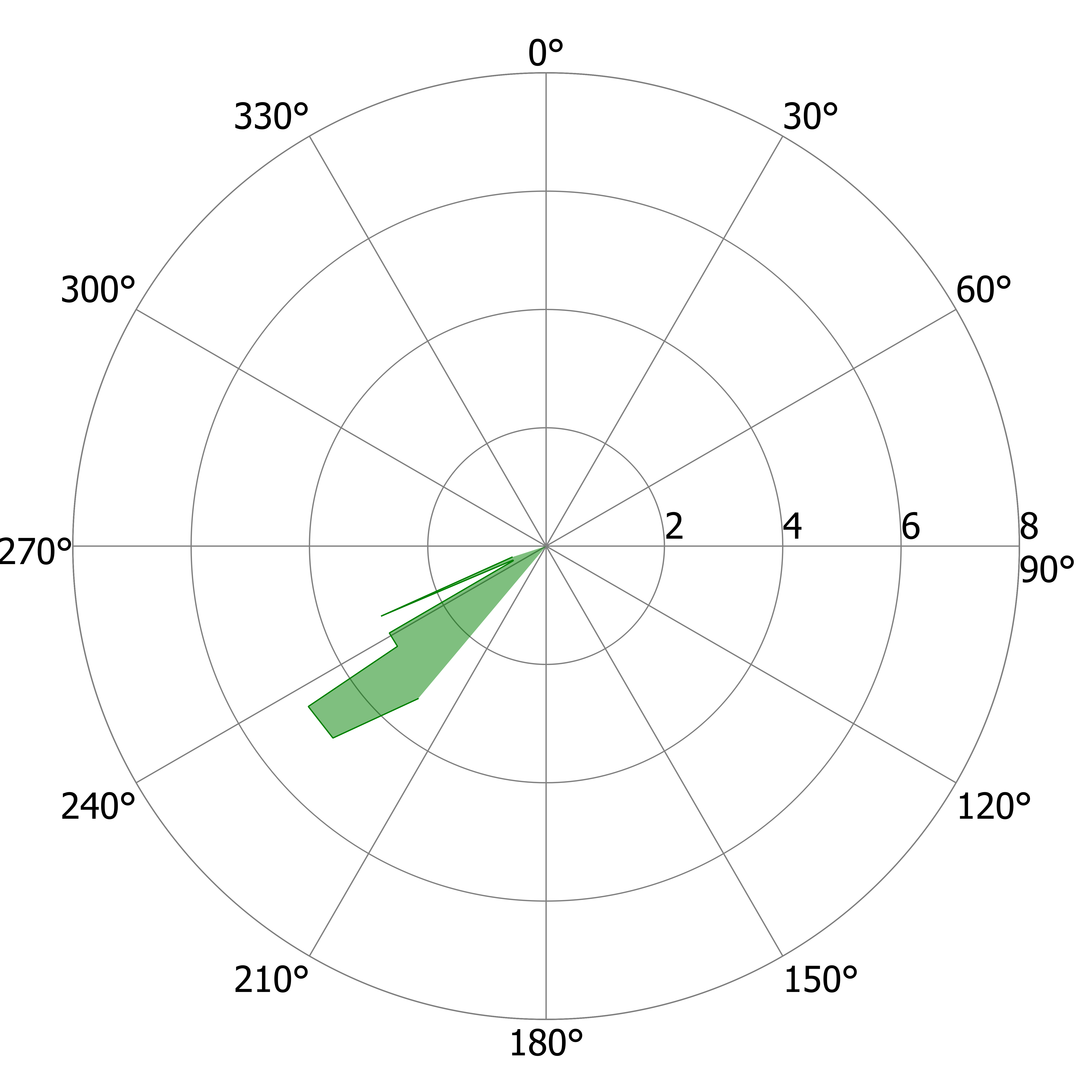

### R_US_g.tiff

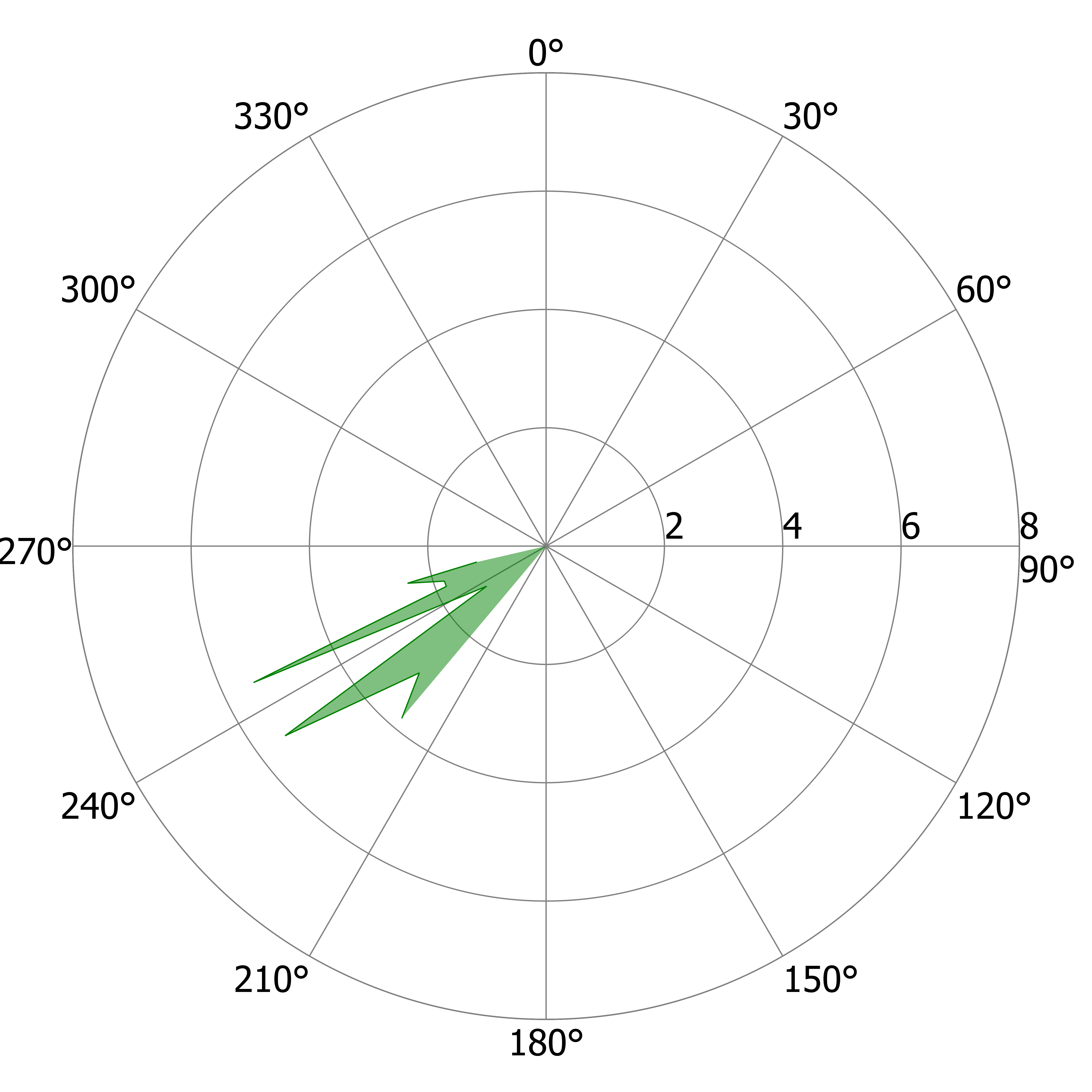

### R_US_h.tiff

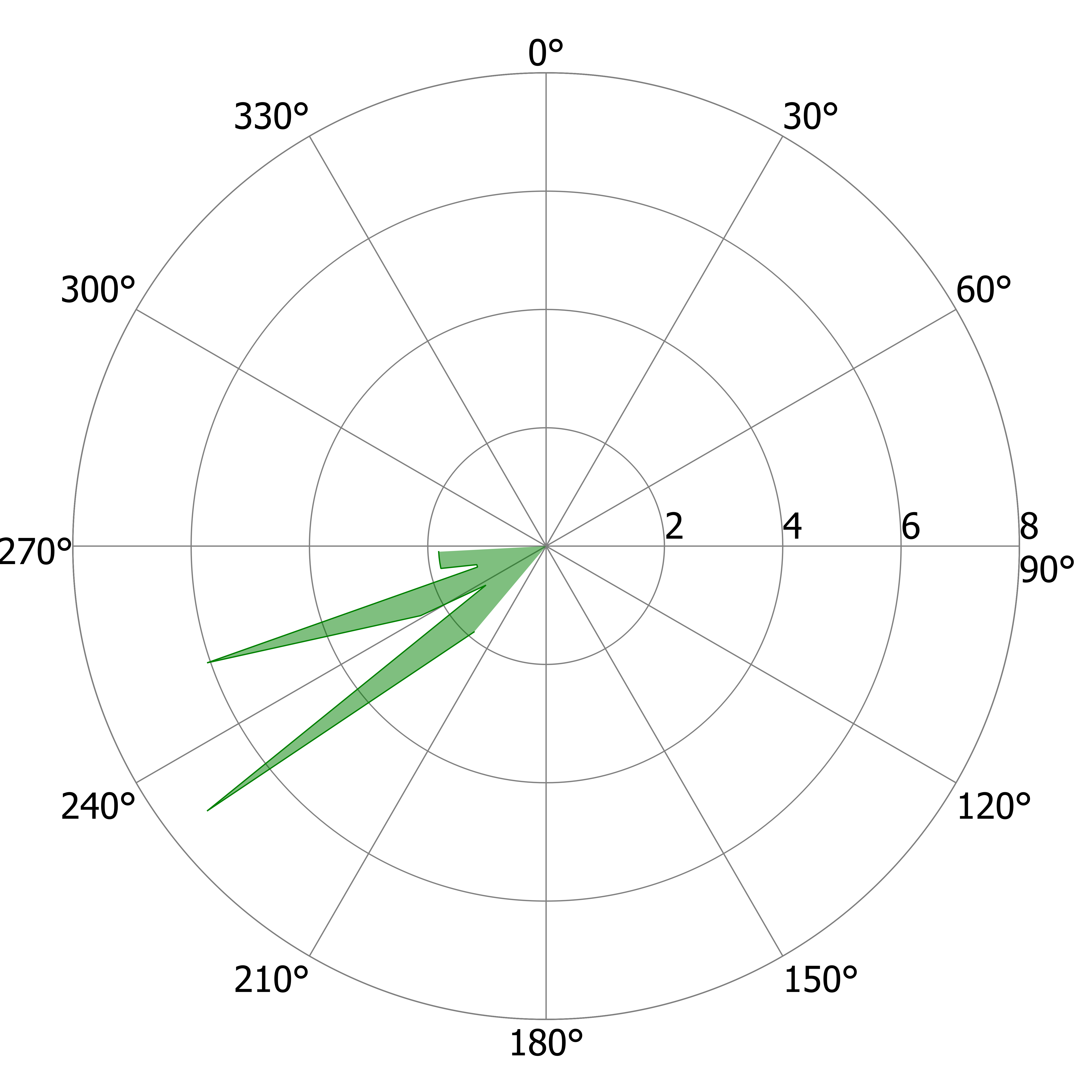

### R_US_i.tiff

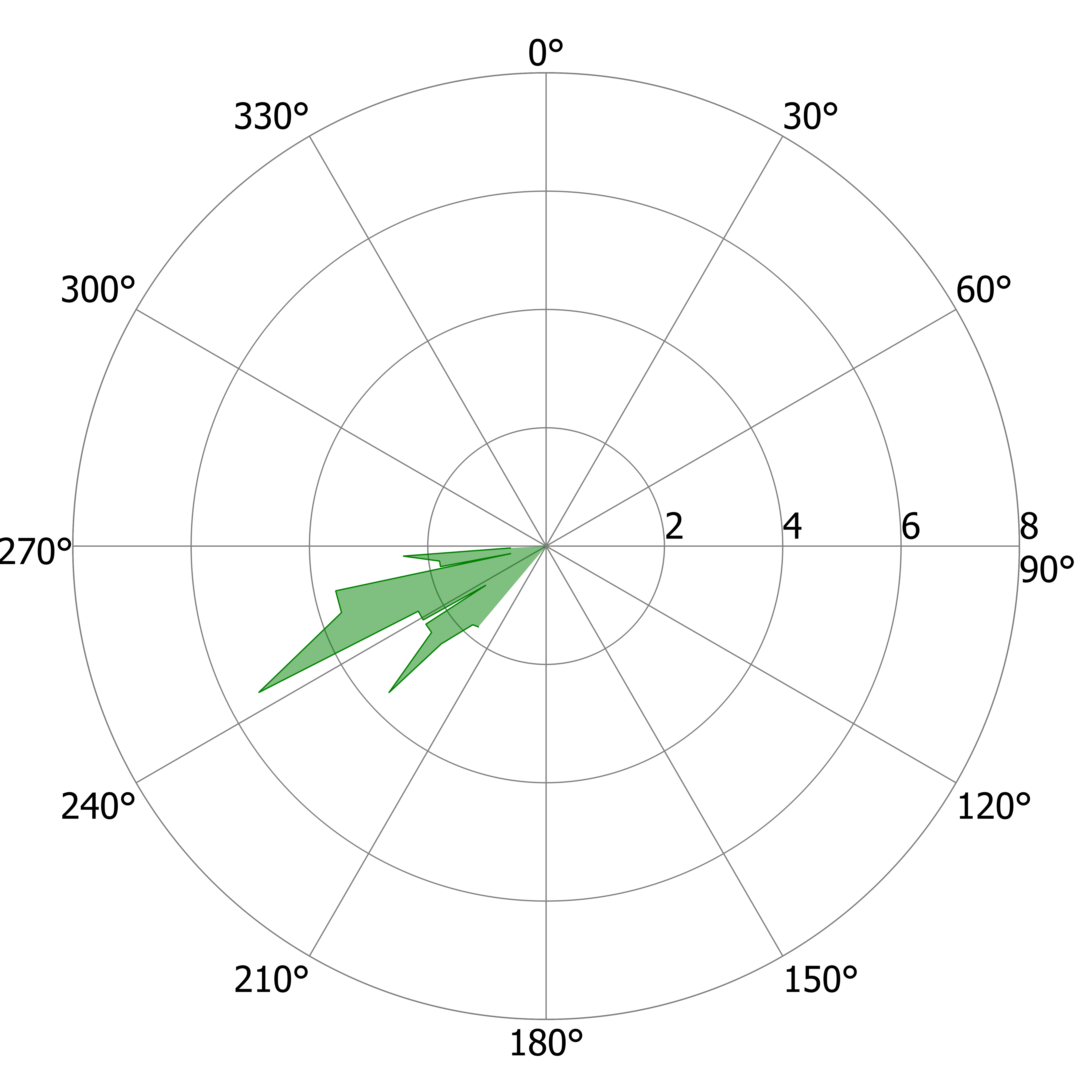
